# Statistical Optimization of Hydrazone-Crosslinked Hyaluronic Acid Hydrogels for Protein Delivery

**DOI:** 10.1101/2023.07.14.549125

**Authors:** Esther A. Mozipo, Alycia N. Galindo, Jenna D. Khachatourian, Conor G. Harris, Jonathan Dorogin, Veronica R. Spaulding, Madeleine R. Ford, Malvika Singhal, Kaitlin C. Fogg, Marian H. Hettiaratchi

## Abstract

Hydrazone-crosslinked hydrogels are attractive protein delivery vehicles for regenerative medicine. However, each regenerative medicine application requires unique hydrogel properties to achieve an ideal outcome. The properties of a hydrogel can be impacted by numerous factors involved in its fabrication. We used design of experiments (DoE) statistical modeling to efficiently optimize the physicochemical properties of a hyaluronic acid (HA) hydrazone-crosslinked hydrogel for protein delivery for bone regeneration. We modified HA with either adipic acid dihydrazide (HA-ADH) or aldehyde (HA-Ox) functional groups and used DoE to evaluate the interactions of three input variables, the molecular weight of HA (40 or 100 kDa), the concentration of HA-ADH (1-3% w/v), and the concentration of HA-Ox (1-3% w/v), on three output responses, gelation time, compressive modulus, and hydrogel stability over time. We identified 100 kDa HA-ADH_3.0_HA-Ox_2.33_ as an optimal hydrogel that met all of our design criteria, including displaying a gelation time of 3.7 minutes, compressive modulus of 62.1 Pa, and minimal mass change over 28 days. For protein delivery, we conjugated affinity proteins called affibodies that were specific to the osteogenic protein bone morphogenetic protein-2 (BMP-2) to HA hydrogels and demonstrated that our platform could control the release of BMP-2 over 28 days. Ultimately, our approach demonstrates the utility of DoE for optimizing hydrazone-crosslinked HA hydrogels for protein delivery.

**Graphical Abstract:** 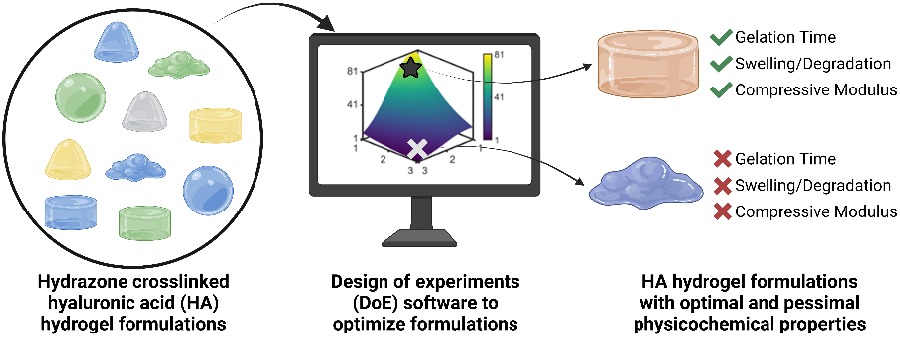

## Introduction

Recombinant protein delivery is a promising strategy for the repair of injured or damaged tissues including bone, muscle, nerves, vasculature, and skin.(1) Specifically in the case of bone repair, the delivery of bone morphogenetic protein-2 (BMP-2) has demonstrated success as a bone regeneration strategy in both animal models and human patients.(2) A wide variety of biomaterials have been investigated for protein delivery to the body, and their properties have been tuned to enable control over protein release rate, material degradation rate, material compressive modulus, and injectability. However, tuning all of these properties simultaneously to develop the ideal biomaterial for protein delivery is challenging. Furthermore, biomaterial delivery vehicles must be individually optimized for different clinical applications, as unique properties may be necessary to control the release of different proteins to different tissues. Hydrogels are chemically or physically crosslinked water-swollen polymer networks that can be effective vehicles for the delivery of a variety of cells, proteins, and drugs due to their ability to support cell viability, provide sustained release of cargo, and mimic the mechanical and biochemical properties of tissues.(1,3–5) The relative ease in which a variety of natural and synthetic polymers can be chemically modified with different functional groups has facilitated the fabrication of diverse hydrogels with a wide array of properties suitable for numerous applications. Varying individual input parameters to create a hydrogel, such as polymer concentration, crosslinking ratio, and molecular weight of the polymer, one at a time to determine the combination of parameters that yields a hydrogel with the optimal properties can be time-intensive and expensive. While extensive prior work has been performed to determine the ideal physicochemical properties of hydrogels for a variety of biomedical applications, there has been limited investigation of the collective, combinatorial effects of varying multiple input parameters on the properties of hydrogels in the context of protein delivery.

To tackle this challenge, we used a statistical optimization technique known as Design of Experiments (DoE) to efficiently screen the effects of multiple input parameters on several hydrogel properties and create a mathematical model to predict the optimal and pessimal hydrogel formulations given a specific set of design criteria. Design of experiments (DoE) is a method of experimental design in which a combination of factors are varied simultaneously instead of the typical “one factor at a time” approach. The responses to each set of input parameters is mapped to generate a model for how the different input parameters collectively influence each hydrogel property (i.e., the output responses), known as the response surface methodology.(6,7) Mathematical models can then be used to define the response surface and predict expected outcomes. DoE was originally developed in the 1920s for agricultural purposes, but has since been used in a broad array of applications, including the development and optimization of biomaterials.(7–10) It lends itself particularly well to applications such as hydrogel design, in which it is highly likely that multiple input parameters will have interactive effects on hydrogel properties, which makes individually varying input parameters less informative. We and others have used DoE to predict responses of hydrogels and cells within hydrogels in systems with multiple input variables.(8,11–14) This approach can be widely used in regenerative medicine applications such as scaffold design, cell culture optimization, and the development of in vitro model systems, accelerating the timeframe in which new platforms can be optimized.(15–17)

In this study, we were specifically interested in tuning the properties of hyaluronic acid hydrogels formed via hydrazone crosslinking. Hyaluronic acid (HA) is an abundant biopolymer found throughout the body. It is a critical extracellular matrix component of connective tissue, skin, and the brain that functions by creating a hydrated polymer matrix for cells to migrate and proliferate.(18–20) HA also exists at a range of molecular weights, which can activate different cell signaling pathways.(21,22) Although HA itself is soluble in aqueous media, modifications can be made to the carboxylic acid, adjacent alcohol, and secondary alcohol groups on the HA backbone to form crosslinks throughout the polymer, resulting in a stable three-dimensional, water-swollen network. HA hydrogels have been used successfully in several biomedical applications, including protein and cell delivery and tissue regeneration.(23–25)

While there are numerous criteria that may be considered important for the design of a hydrogel for protein delivery, we narrowed down our desired hydrogel properties to three essential criteria for optimization: compressive modulus, gelation time, and mass change over four weeks. Compressive modulus can have a profound effect on cell differentiation,(26,27) while the bulk stiffness of an implanted material must be matched to the target tissue to avoid eliciting mechanical irritation and subsequent inflammation. Hydrogel gelation time should allow for sufficient time to mix components (different polymers, proteins, cells) before gelation. Typically, hydrogel gelation times of less than 10 minutes are preferred.(28,29) Finally, the mass change of the hydrogel due to swelling or degradation of the crosslinked network can impact cellular responses over time, such cell spreading, cell-cycle progression, and differentiation.(30– 32)

We formed HA hydrogels using dynamic covalent hydrazone crosslinks as this approach can yield a wide range of physicochemical properties, making them an attractive hydrogel platform for numerous regenerative medicine applications.(33–40) The properties of hydrazone-crosslinked HA hydrogels can be easily tuned by adjusting the ratios of the two dynamic covalent crosslinking polymers: oxidized HA (HA-Ox) and HA adipic acid dihydrazide (HA-ADH). Thus, we narrowed our input parameters to HA molecular weight, HA-Ox concentration, and HA-ADH concentration. We then used DoE to identify the contributions and combinatorial effects of the concentration of HA-ADH, the concentration of HA-Ox, and the molecular weight of HA on hydrogel compressive modulus, gelation time, and mass change over time. We then created a statistical model that predicted physicochemical properties as a function of input parameters. This model was then validated by evaluating the hydrogels predicted to be the optimal and pessimal delivery vehicles. Finally, as a proof-of-concept for protein delivery, we conjugated a protein-binding peptide known as an affibody to the HA hydrogels using thiol-ene click chemistry to enable affinity-controlled delivery of bone morphogenetic protein-2 (BMP-2), which is a potent osteogenic protein that is clinically used as a bone graft substitute. Overall, we developed a hydrogel platform and used DoE analysis to optimize the vast tunability of hydrazone-crosslinked HA hydrogels for protein delivery for tissue regeneration.

## Materials and Methods

### Materials

Sodium Hyaluronate (HA, 40 kDa and 100 kDa) was purchased from Lifecore Biomedical LLC (Chaska, MN). Adipic acid dihydrazide (ADH) was purchased from Spectrum Chemical (Gardena, CA). Hydroxybenzotriazole (HOBt) was purchased from Chem Impex (Wood Dale, IL). 1-Ethyl-3-(3-dimethylaminopropyl)carbodiimide (EDC) was purchased from G Biosciences (St. Louis, MO). Sodium Periodate, AmberLite® MB Ion Exchange Resin, tert-butyl alcohol (TBA-OH), DMSO, 4-Dimethylaminopyridine (DMAP), Di-tert-butyl decarbonate (BoC_2_O) and 5-norbornene-2-carboxylic acid were purchased from Sigma Aldrich (St. Louis, MO). Irgacure 2959 was purchased from Advanced Biomatrix (Carlsbad, CA). Recombinant human bone morphogenetic protein-2 (BMP-2) and the human BMP-2 DuoSet enzyme-linked immunosorbent assay (ELISA) were purchased from R&D Systems (Minneapolis, MN).

### Synthesis of Adipic Acid Dihydrazide HA (HA-ADH)

HA (200 mg, 0.527 mmol) was dissolved in 20 mL of diH_2_O to form a 1% w/v solution. Adipic acid dihydrazide (183.75 mg, 1.05 mmol) and hydroxy-benzotriazole (142.53 mg, 1.05 mmol) were added to the HA solution and the pH was adjusted to 4.75. EDC was added (91.00 mg, 0.47 mmol) to the solution, and the pH was monitored and maintained at 4.75 for 4 hours using 1 M HCl and 1 M NaOH. The solution was stirred for 24 hours at room temperature and dialyzed against 0.1 M NaCl in diH_2_O for 2 days followed by diH_2_O for 2 days. The solution of HA functionalized with ADH (HA-ADH) was sterile filtered through a 0.22 μm filter, frozen at -80 °C, and lyophilized using a VirTis BenchTop Pro lyophilizer (SP Scientific, Warminster, PA).

### Synthesis of Oxidized HA (HA-Ox)

HA (200 mg, 0.561 mmol) was dissolved in 10 mL of diH20 to form a 2% w/v solution. Sodium periodate (322.00 mg, 0.281 mmol) was added to the solution and stirred for 4 hours at room temperature protected from light. The reaction was quenched with 1 mL of propylene glycol and dialyzed against diH_2_O for 3 days. The oxidized HA solution (HA-Ox) was sterile filtered through a 0.22 μm filter, frozen at -80 °C, and lyophilized.

### Synthesis of HA Norbornene (HA-Nor)

HA (1.01 g, 2.506 mmol) was dissolved in 50 mL diH_2_O to form a 2% w/v solution. AmberLite™ MB Ion Exchange Resin (3.03 g) was added to the reaction and stirred at room temperature for 5 hours. The resin was vacuum-filtered, and the filtrate was titrated to pH of 7 with TBA-OH and dialyzed against diH_2_O for 3 days. The HA-TBA solution was sterile filtered through a 0.22 μm filter, frozen at -80 °C, and lyophilized.

HA-TBA (153 mg, 0.35 mmol) was dissolved in 0.75 mL DMSO to form a 2% w/v solution. The flask was purged with N_2_ for 5 minutes, then 5-norbornene-2-carboxylic acid (145 mg, 1.05mmol) and DMAP (145 mg, 1.049 mmol) was added to the reaction flask. Boc_2_O was added via syringe (32 μL, 0.14 mmol). The solution was stirred at 45 °C for 20 hours, then quenched with cold diH_2_O (10 mL) before being dialyzed against diH_2_O for 3 days. Norbornene-functionalized HA (HA-Nor) was precipitated from the solution by adding NaCl and cold acetone (30 mL), then filtered and dialyzed against diH_2_O for 3 days. The solution was sterile filtered through a 0.22 μm filter, frozen at -80 °C, and lyophilized.

### Oxidization of HA Norbornene (HA-Nor-Ox)

HA-Nor (91.75 mg, 0.18 mmol) was dissolved in diH_2_O to form a 1% w/v solution. Sodium periodate (10.57 mg, 0.049 mmol) was dissolved in diH_2_O to create a 0.5 M solution and added to the HA-Nor solution. The solution was stirred overnight at room temperature protected from light. The reaction was quenched with 1 mL of propylene glycol and dialyzed against diH_2_O for 3 days. The oxidized HA-Nor solution (HA-Nor-Ox) was sterile filtered through a 0.22 μm filter, frozen at -80 °C, and lyophilized.

### Affibody Bioconjugation

BMP-2-specific affibodies were expressed in E. coli and purified as previously described.(41) HA-Nor-Ox (14.5 mg) was dissolved in 1.5 mL of 0.71 mg/mL affibody in PBS (1.45 ×10^−4^ mmol) to create a solution with 1:1 ratio of affibody to HA-Nor-Ox. 15 μL of 10% w/v Irgacure 2595 in methanol was added to the reaction vial, stirred, and illuminated with 365 nm light for 10 minutes. The solution was dialyzed against HEPES buffer pH 7.0 for 1 day and diH_2_O for 2 days. The affibody-conjugated solution of HA (HA-Nor-Ox-Affibody) was sterile filtered through a 0.22 μm filter, frozen at -80 °C, and lyophilized. The amount of affibody conjugated to the polymer was quantified by dissolving 2-3 mg of HA-Nor-Ox-Affibody in PBS at 2% w/v and determining the protein concentration of the solution using a Pierce 660 Protein Assay Kit (Thermo Fisher Scientific).

### Degree of Modification (DOM) Determination

The degree of chemical modification of HA-ADH and HA-Nor was quantified using proton nuclear magnetic resonance spectroscopy (^1^H NMR, 500Hz, Bruker, USA). 2-5 mg of modified polymer was dissolved in 600 μL of deuterium oxide (D_2_O) for ^1^H NMR (256 scans). To calculate the degree of modification for HA-ADH, the peaks from the aliphatic chain were integrated from 2.5-2.1 ppm and 1.7-1.44 ppm (8H) and normalized to the n-acetyl methyl group peaks from 1.8-2 ppm (3H) (**Figure S1**).(42) The degree of modification for HA-Nor was calculated by integrating the vinyl peaks from 5.7-6.3 ppm and normalizing to the n-acetyl methyl group from 1.8-2 ppm (3H) (**Figure S2**).(42)

The degree of oxidation for HA-Ox was determined by hydroxylamine hydrochloride titration.(43,44) Briefly, 100 mg of oxidized polymer was dissolved in 0.25 N hydroxylamine hydrochloride containing 0.05% w/v methyl orange reagent for 2 hours. The solution was then titrated with 0.1 M NaOH while monitoring the pH until the pH indicator changed from red to yellow at a pH of 4. The reaction of hydroxylamine hydrochloride with the aldehydes on oxidized HA is shown below.

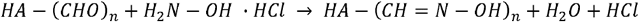

Equation 1 was used to calculate the percent degree of modification (% DOM) or the percentage of HA disaccharide units that contain aldehydes.

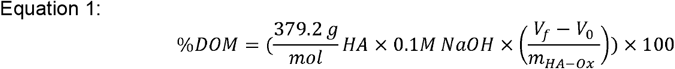

where 379.2 g/mol is the monomeric molecular weight of HA, 0.1 M is the molarity of NaOH used to adjust the solution pH, V_f_ is the final volume of NaOH in the burette after titration recorded in liters, V_0_ is the initial volume of NaOH in the burette before titration recorded in liters, and m_HA-Ox_ is the mass in g of HA-Ox used in the titration.

### Preparation of HA Hydrogels

HA-ADH and oxidized HA (HA-Ox, HA-Nor-Ox, or HA-Nor-Ox-Affibody) were each reconstituted at 1-3% w/v in PBS. Hydrogels were then prepared by mixing 50 μL of each polymer in 2 mL microcentrifuge tube or 8 mm diameter cylindrical silicone mold to create 100 μL hydrogels.

### Physicochemical Characterization of Hydrogels Gelation Time

The gelation time of each hydrogel was determined using the inverted tube test.(45) 50 μL each of HA-ADH and HA-Ox were pipetted into 2 mL microcentrifuge tubes to create 100 μL hydrogels and inverted every minute until gelation occurred. The time at which there was no viscous flow was recorded as the gelation time.

### Compressive Modulus

Compressive moduli of the hydrogels were evaluated using a Discovery Hybrid Rheometer-2 (DHR-2) (TA Instruments, New Castle, DE). 100 μL hydrogels were formed in 8 mm cylindrical silicone molds. The hydrogels were crosslinked overnight at room temperature before being removed from the molds and placed on the rheometer. Hydrogels were compressed within an 8 mm parallel plate to 15% of their original height at 37 °C, and the slope of the linear stress versus strain curve was used to calculate the compressive modulus.(46)

### Mass Change Over Time

Hydrogels were formed overnight in pre-weighed 2 mL microcentrifuge tubes and incubated in 100 μL of 1x PBS. At 0 hours, 24 hours, 4, 7, 10, 14, 21, and 28 days, the supernatant was removed, hydrogels were blotted of excess PBS with a KimWipe, and the tube containing the hydrogel was weighed. The mass of the hydrogel was recorded, and fresh PBS was added to the tube. The percentage mass change was calculated using Equation 2.

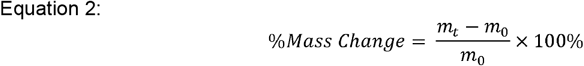

where m_t_ is the mass of the hydrogel at time t and m_0_ is the initial mass of the hydrogel immediately after it was formed.

### Release of BMP-2 from HA Hydrogels

Release of BMP-2 was assessed from HA hydrogels with and without conjugated BMP-2-specific affibodies over 28 days. To load BMP-2 into the hydrogels, BMP-2 was added to the HA-Ox and HA-Nor-Ox-Affibody solutions and then crosslinked with HA-ADH. For hydrogels without affibodies, 200 ng of BMP-2 were added to a 50 μL solution of 2% w/v HA-Ox. For affibody-conjugated HA hydrogels, BMP-2 was added to a 50 μL solution of 2% w/v HA-Nor-Ox-Affibody, which resulted in 1000 molar excess of affibody to BMP-2. For each hydrogel, 50 μL of HA-ADH were added to the HA-Nor-Ox or HA-Nor-Ox-Affibody solutions to create 100 μL HA hydrogels in 2 mL microcentrifuge tubes. 900 μL of 0.1% w/v BSA in PBS was added to each tube, and the hydrogels were incubated at 37 □C. 300 μL aliquots of supernatant were collected and replenished with fresh 0.1% w/v BSA in PBS at multiple time points over 28 days. BMP-2 in the supernatant was quantified using a BMP-2-specific ELISA. Cumulative BMP-2 release was calculated by adding the total mass of BMP-2 in each collected aliquot. Cumulative fractional release versus the square root of time was plotted, and the data were fit to a short time approximation of one-dimensional diffusion from a thin sheet, and the slope of Fickian diffusion, k, was determined using Equation 3.(47,48)

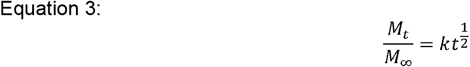

where M_t_ is the mass of BMP-2 released at time t, M_t_ is the total mass of BMP-2 released at the end of the experiment, and k is the slope of Fickian diffusion.

### Design of Experiments

Design of experiments (DoE) statistical optimization was used to investigate how the chemical composition of the hydrogels influenced their physicochemical properties. More specifically, D-optimal design was used to generate a design matrix of 15 hydrogel formulations with 1 centrally repeated condition (**Table 1**). The input variables for HA hydrogel composition were the concentration of HA-ADH (1-3% w/v), the concentration of HA-Ox (1-3% w/v), and the molecular weight of HA (40 kDa or 100 kDa) used to synthesize the hydrogel. The concentrations of HA-ADH and HA-Ox were continuous variables that were examined at equidistant low, medium, and high levels, whereas the molecular weight was a categorical variable that was examined at either 40 kDa or 100 kDa. The physicochemical properties quantified as response variables were gelation time, compressive modulus, and mass change at 1, 7, 14, 21, and 28 days after hydrogel fabrication.

**Table 1.** HA hydrogel formulations evaluated to generate D-optimal multilinear regression model. 15 unique hydrogel formulations were evaluated in triplicate with a centrally repeating condition at 100 kDa HA-ADH_2.0_HA-Ox_2.0_ evaluated with 9 replicates to determine the system’s variance. The centrally repeated condition is bolded.

| HA-ADH (% w/v) | HA-Ox (% w/v) | Molecular Weight |
| --- | --- | --- |
| 1 | 1 | 40 kDa |
| 1 | 3 | 40 kDa |
| 1.67 | 1 | 40 kDa |
| 2 | 2 | 40 kDa |
| 3 | 1 | 40 kDa |
| 3 | 3 | 40 kDa |
| 1 | 1 | 100 kDa |
| 1 | 3 | 100 kDa |
| 1 | 1.67 | 100 kDa |
| 1.67 | 3 | 100 kDa |
| 2 | 2 | 100 kDa |
| 2 | 2 | 100 kDa |
| **2** | **2** | **100 kDa** |
| 2.33 | 1 | 100 kDa |
| 3 | 1 | 100 kDa |
| 3 | 3 | 100 kDa |
| 3 | 2.33 | 100 kDa |

MODDE-Pro 13.0 software (Sartorius, Fremont, CA) was used to generate the D-optimal experimental matrix of 15 hydrogel formulations and analyze the experimental data obtained. Experimental data were collected from each of the hydrogel formulations then used to fit a multiple linear regression model for each response variable as a function of the statistically significant input variables. An input variable was considered statistically significant if it had a *p* value of < 0.05 as determined by ANOVA. The model performance was further assessed using R^2^, adjusted R^2^, and Q^2^. R^2^ describes the variation of the response explained by the model for every variable. Adjusted R^2^ describes the variation of the response explained by the model only for the variables that affect the response. Q^2^ describes the variation of the response predicted by the model for new data. R^2^ and adjusted R^2^ values close to 1.0 indicate a high correlation between observed and predicted values. Q^2^ above 0.50 and a difference between R^2^ and Q^2^ values below 0.30 validate that the model works independently of the specific data used to train the model.(49,50) Residual plots, Box-Cox plots, and ANOVA tables were used to diagnose outliers, trends, or lack of fit. The model was then refined by removing any outliers and input variables that were not statistically significant to increase the validity of our model.

Our overall engineering objective was to maximize the compressive modulus, set the gelation time to a value within 1-10 min, and set the mass change at all time points to 0%. Thus, to validate the predictive models, we generated hydrogels predicted to either fulfill these optimization parameters (optimal hydrogels) or fulfill reciprocal objectives (pessimal hydrogels). Specifically, the pessimal hydrogels were predicted to minimize the compressive modulus, maximize the gelation time, and either maximize or minimize the hydrogel mass change at all time points (i.e., induce hydrogel swelling or degradation).

## Results and Discussion

### Functionalization of HA polymers

HA was modified with either aldehyde (HA-Ox) or ADH functional groups (HA-ADH) for hydrazone crosslinking (**Figure 1**). Because polymer molecular weight can influence crosslinking density and subsequent physicochemical properties of the hydrogel,(28) we functionalized both 40 kDa and 100 kDa HA for hydrogel fabrication and evaluation. To expose aldehyde groups on HA, we oxidized the polymer in the presence of sodium periodate (NaIO_4_) for 4 h in the dark before quenching the reaction with propylene glycol. Using the same starting material, we added ADH to HA in the presence of EDC at a pH of 4.75 for 24 h to create HA-ADH. The degree of oxidization was characterized using titration with hydroxylamine hydrochloride and was found to be 27.5±5.9% and 41.4±12.8% of HA repeating disaccharide units for 40 kDa and 100 kDa HA-Ox, respectively. The degree of modification of the ADH functional groups on the polymer was 67.6 ±1.2% for 40 kDa HA-ADH and 50.6 ±10.1% for 100 kDa HA-ADH as characterized by ^1^H NMR (**Figure S1**). The degrees of modification and yields of all modified polymers are summarized in **Table S1**. Degrees of oxidation between 27-82%(33,51) and degrees of modification with ADH between 30-49%(52,53) have been previously reported, which is consistent with our results. We then used these modified HA polymers to fabricate the 15 hydrogels in **Table 1** and evaluated their gelation time, compressive moduli, and mass change over time to generate a predictive model for DoE analysis.

**Figure 1.**
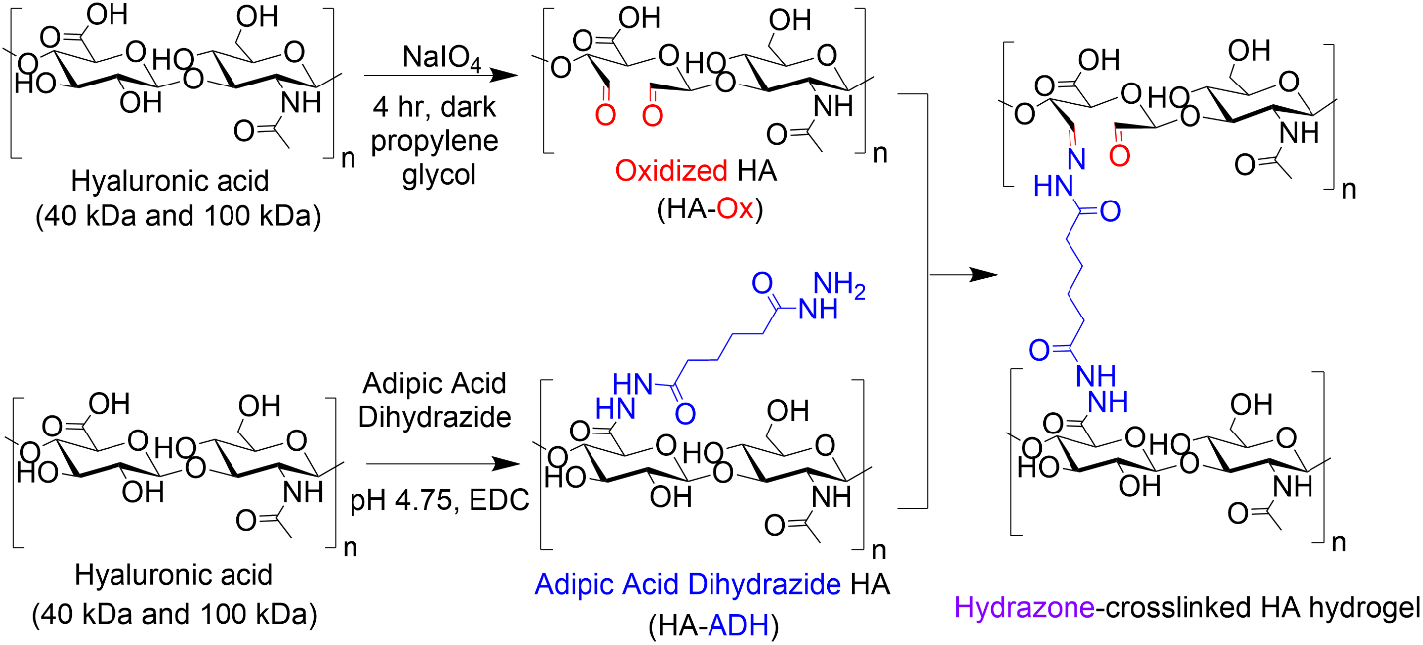
Functionalization of HA and hydrogel formation via hydrazone crosslinking. Hyaluronic acid (40 kDa or 100 kDa) was either stirred with sodium periodate in water for 4 h in the dark to oxidize the polymer and reveal aldehyde functional groups (HA-Ox) or mixed with adipic acid dihydrazide (ADH) and 1-ethyl-3-(3-dimethylaminopropyl)carbodiimide (EDC) in water and maintained at pH 4.75 for 24 h to create ADH-functionalized HA (HA-ADH). All polymers were dialyzed, filtered, and lyophilized before use. HA-Ox and HA-ADH were re-dissolved in phosphate buffered saline (PBS) and mixed at room temperature and pH 7.4 to form hydrazone-crosslinked HA hydrogels.

### Hydrogel Gelation Time

HA hydrogels were fabricated by mixing 50 μL of HA-ADH and 50 μL of HA-Ox at different polymer concentrations and molecular weights. Inversion tests were used to quantify the gelation time of each of the HA hydrogel formulations. As expected, increasing the polymer content of the hydrogels increased the crosslinking density of the polymer network and subsequently decreased the gelation time (**Figure 2A**). Changing the polymer concentrations of the 40 kDa hydrogels had a greater effect on gelation time compared to the 100 kDa hydrogels, such that the 40 kDa hydrogels had a larger range of average gelation times ranging from 4.5 to 292.7 min. Additionally, increasing the HA-ADH content in the 40 kDa hydrogels decreased the gelation time of the hydrogels, except for the HA-ADH_1.0_HA-OX_3.0_ hydrogel. The gelation times of the 100 kDa hydrogels all fell between 1.4 and 71.5 min. The 100 kDa HA hydrogels similarly exhibited decreasing gelation time in response to increasing HA-ADH content, except for the HA-ADH_1.67_HA-OX_3.0_ and HA-ADH_1.0_HA-OX_3.0_ hydrogels, which demonstrated faster gelation times than the HA-ADH_2.0_HA-OX_2.0_ hydrogel. Excess HA-Ox content compared to HA-ADH content also resulted in shorter gelation times in both the 40 kDa and 100 kDa hydrogels. Overall, the 100 kDa hydrogels gelled faster than 40 kDa hydrogels, potentially due to increased entanglement of the longer polymer chains.(54)

**Figure 2.**
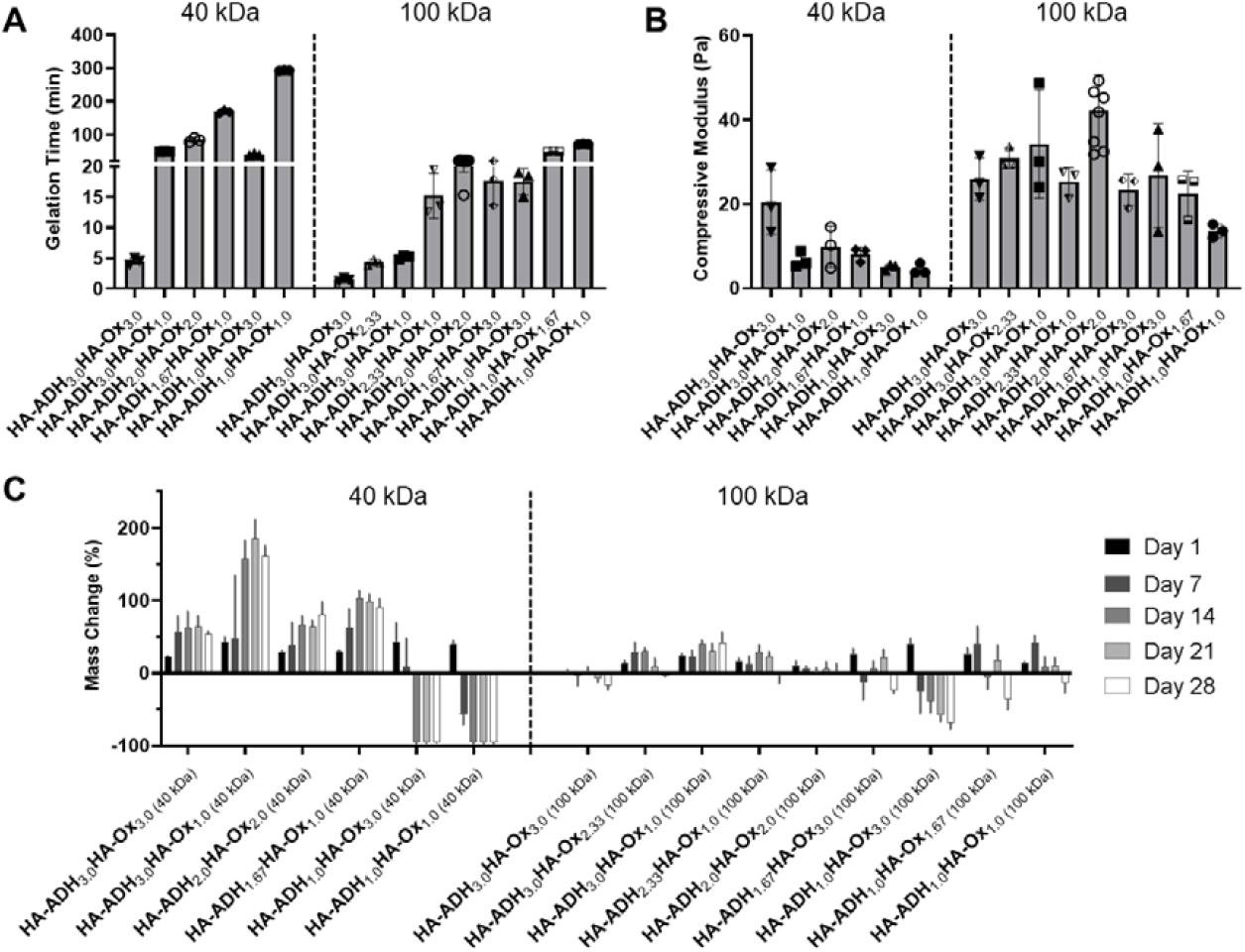
Physicochemical properties for each of the 15 hydrogel formulations for the D-optimal design. HA hydrogels were fabricated using 40 kDa or 100 kDa HA-Ox and HA-ADH at different polymer weight percentages. A) Gelation time of HA hydrogels. B) Compressive moduli of HA hydrogels. C) Mass change of HA hydrogels incubated in phosphate-buffered saline at room temperature for 28 days. Data presented as mean ± SD with 3 replicates for all hydrogel formulations, except for the centrally repeated condition, HA-ADH_2.0_HA-Ox_2.0_ (100 kDa), which had 9 replicates. A summary of statistical significance for gelation time, compressive modulus, and mass change can be found in **Tables S2, S3, and S4**, respectively.

### Hydrogel Compressive Modulus

The compressive moduli of the hydrogels ranged from 4.5 to 42.2 Pa (**Figure 2B**). The compressive moduli of all the 100 kDa HA hydrogels fell within a narrow range of 20-40 Pa except for the HA-ADH_1.0_HA-Ox_1.0_ hydrogel, which exhibited an average compressive modulus of 13.6 Pa, which was significantly lower than HA-ADH_3.0_HA-Ox_2.33_ and HA-ADH_2.0_HA-Ox_2.0_ hydrogels and likely due to the lower polymer concentration. In contrast, the compressive moduli of all of the 40 kDa HA hydrogels fell within 4.5 to 20 Pa. Increasing the overall polymer content and HA-ADH content of the 40 kDa HA hydrogels resulted in higher compressive moduli, which was similar to the trends of increasing polymer concentration and HA-ADH content decreasing gelation time.

### Hydrogel Mass Change Over Time

The hydrogels were incubated in PBS at room temperature for 28 days, and their masses were periodically measured to observe swelling and/or degradation over time. For the 40 kDa HA hydrogels, the HA-ADH_3.0_HA-Ox_1.0_ hydrogel exhibited the maximum swelling of 185% of its original mass at 21 days (**Figure 2C**). The HA-ADH_1.0_HA-Ox_3.0_ and HA-ADH_1.0_HA-Ox_1.0_ hydrogels demonstrated complete degradation (i.e., minimum mass change of -100%) at 14 days. For the 100 kDa HA hydrogels, the HA-ADH_1.0_HA-Ox_3.0_ hydrogel demonstrated the greatest degradation of -68% at 28 days, while the HA-ADH_2.33_HA-Ox_1.0_ hydrogel demonstrated the maximum swelling of 62% at 4 days (**Figure 2D**). Overall, the 40 kDa HA hydrogels demonstrated larger changes in mass (swelling or degradation) than the 100 kDa HA hydrogels. This may be due to the lower degree of modification achieved for 40 kDa HA-Ox compared to 100 kDa HA-Ox, which may have resulted in lower crosslinking densities of the polymer network and less stable hydrogels. Hydrogels that contained either excess ADH or aldehyde content also demonstrated increased degradation or swelling, likely due to lower crosslinking density and a less stable crosslinked polymer network.

Our results corroborate the findings of other groups who have fabricated hydrazone-crosslinked HA hydrogels. Similar to the results of our study, others have demonstrated a large range of gelation times, an inverse relationship between gelation time of their HA hydrogels and concentration of ADH crosslinker, and less stable hydrogels with excess aldehydes.(34) In another study exploring hydrazone-crosslinked HA hydrogels for a vitreous substitute, the use of higher amounts of free ADH crosslinker stabilized the hydrogels(33); however, we found that excess HA-ADH increased hydrogel swelling at early time points. A wide range of compressive moduli have been observed with hydrazone-crosslinked HA hydrogels with values ranging between 1-2000 Pa reported in the literature.(36) In comparison, our HA hydrogels exhibited relatively low compressive moduli within a narrow range of 4.5-42.2 Pa. Others have found that a higher degree of modification for HA-ADH led to stiffer hydrogels, which was similar to what we observed for the 40 kDa hydrogels.(35)

Ultimately, our method of fabricating hydrazone-crosslinked HA hydrogels yielded hydrogels with a wide range of physicochemical properties, including large ranges in hydrogel gelation time and mass change, with the capacity to swell, degrade, or maintain a relatively constant mass over 28 days depending on the molecular weight and concentrations of HA-ADH and HA-Ox used (**Table 2**).

**Table 2.** Ranges of physicochemical properties exhibited by HA hydrogels.

| Property | Range |
| --- | --- |
| Gelation Time | 1.4 – 292.7 min |
| Compressive Modulus | 4.5 – 42.2 Pa |
| Mass Change Day 1 | 1.0 – 43.1 % |
| Mass Change Day 7 | -56.3 – 62.2 % |
| Mass Change Day 14 | -100 – 156.9 % |
| Mass Change Day 21 | -100 – 185.3 % |
| Mass Change Day 28 | -100 – 160.7 % |

### Design of Experiments Analysis

After obtaining response data for our 15 hydrogel formulations, we used this data to generate a multivariate model for DoE analysis to better understand how HA-Ox concentration, HA-ADH concentration, and HA molecular weight interacted to produce a wide range of physicochemical properties demonstrated by our platform and to predict the hydrogel formulations that would yield responses that met our desired design criteria. We fit second order multiple linear regression models for each response (gelation time, compressive modulus, mass change at 1, 7, 14, 21, and 28 days) and eliminated the terms that were not statistically significant unless they contributed to square terms or interaction terms that were statistically significant (*p*<0.05) (**Table 3**). There was a significant effect of HA-ADH concentration on all responses except for mass change at day 7 and a significant effect of HA-Ox concentration on all responses except for mass change at day 1. The molecular weight of the HA had a significant effect on all responses. Notably, several squared and interaction terms also had significant effects on individual responses. We found that all of our models fit the success criteria of R^2^ and adjusted R^2^ values close to 1.0, Q^2^ above 0.50, and the difference between R^2^ and Q^2^ values below 0.30. The models with the lowest R^2^ and Q^2^ values, indicating the poorest fit, were for mass change at days 1 and 7. This may be due to large variations in the initial swelling of the hydrogel network when incubated in PBS resulting in unpredictable short-term changes in mass compared to hydrogels that have swollen to a stable, more predictable equilibrium at later time points (days 14, 21, and 28).(55)

**Table 3.** ANOVA assessment of regression models. Excluded non-significant values are indicated with a dash (-).

| Response | Gelation Time | Compressive Modulus | Mass Change Day 1 | Mass Change Day 7 | Mass Change Day 14 | Mass Change Day 21 | Mass Change Day 28 |
| --- | --- | --- | --- | --- | --- | --- | --- |
| p-values |  |  |  |  |  |  |  |
| [HA-ADH] | <0.0001 | 0.0258 | 0.0136 | 0.612 | <0.0001 | <0.0001 | <0.0001 |
| [HA-Ox] | <0.0001 | 0.173 | 0.725 | 0.000539 | <0.0001 | <0.0001 | <0.0001 |
| Molecular Weight (MW) | <0.0001 | 0.00122 | 0.000702 | 0.000222 | 0.000494 | 0.00107 | 0.000234 |
| [HA-ADH]<sup>2</sup> | 0.000985 | 0.00375 | 0.0399 | - | - | - | - |
| [HA-Ox]<sup>2</sup> | - | 0.000782 | - | - | - | - | - |
| MW<sup>2</sup> | - | - | - | - | - | - | - |
| [HA-ADH]*<br>[HA-Ox] | - | - | 0.00213 | 0.00237 | - | - | - |
| [HA-ADH]*<br>MW | - | - | - | - | 0.00166 | 0.000823 | 0.0125 |
| [HA-Ox]*<br>MW | - | - | - | - | 0.000964 | 0.00304 | 0.00763 |
| Lack of Fit | 0.177 | 0.363 | 0.948 | 0.213 | 0.410 | 0.472 | 0.548 |
| Total Sample Size (N) | 17 | 13 | 17 | 16 | 16 | 16 | 16 |
| Degrees of Freedom (DF) | 12 | 7 | 11 | 11 | 10 | 10 | 10 |
| R<sup>2</sup> | 0.964 | 0.970 | 0.828 | 0.852 | 0.950 | 0.937 | 0.936 |
| **R<sup>2</sup>-adjusted** | 0.952 | 0.903 | 0.750 | 0.799 | 0.926 | 0.905 | 0.904 |
| **Q<sup>2</sup>** | 0.918 | 0.949 | 0.601 | 0.683 | 0.710 | 0.840 | 0.792 |

Three-dimensional response surface plots of gelation time for the 40 kDa HA hydrogels demonstrated that higher polymer content resulted in faster gelation with the shortest gelation time occurring at HA-ADH_3.0_HA-Ox_3.0_ and longest gelation time occurring at HA-ADH_1.0_HA-Ox_1.0_ (**Figure 3A**). This response surface was identical but smaller in magnitude for the 100 kDa HA hydrogels (**Figure 3B**). These plots indicate that gelation time was dependent on the concentrations of both HA-ADH and HA-Ox. The response surfaces of compressive modulus for 40 kDa and 100 kDa HA hydrogels both depicted a negative quadratic dependence on the HA-Ox and HA-ADH concentrations with a maximum compressive modulus predicted to occur at HA-ADH_2.21_HA-Ox_2.08_. For the 40 kDa HA hydrogels, this maximum compressive modulus was predicted to be 39.2 Pa (**Figure 3C**), while, for the 100 kDa HA hydrogels, the maximum compressive modulus was predicted to be 27.3 Pa (**Figure 3D**). The relationship between mass change and input parameters was different at different time points. Mass change at day 1 was dependent on interactions between HA-ADH and HA-Ox concentrations and demonstrated a positive quadratic dependence on HA-ADH. The minimal mass change for both 40 kDa and 100 kDa hydrogels occurred at HA-ADH_3.0_HA-Ox_3.0_ and was found to be at 4.65% and 21.1%, respectively (**Figure 3E, Figure 3F)**. Mass change at day 7 demonstrated a linear dependence on HA-ADH and HA-Ox concentrations, suggesting there were no interactions between HA-ADH and HA-Ox concentrations for this response. For both 40 kDa and 100 kDa hydrogels, the minimum mass change occurred at HA-ADH_1.0_HA-Ox_3.0_; however, the predicted minimum was 9.49% for 40 kDa hydrogels (**Figure 3G**), and -22.61% for 100 kDa hydrogels (**Figure 3H**). At day 14, the model predicted a minimum mass change of -74.87% at HA-ADH_1.0_HA-OX_3.0_ and a maximum mass change of 174.16% at HA-ADH_3.0_HA-OX_1.0_ for 40 kDa hydrogels (**Figure 3I**).

**Figure 3.**
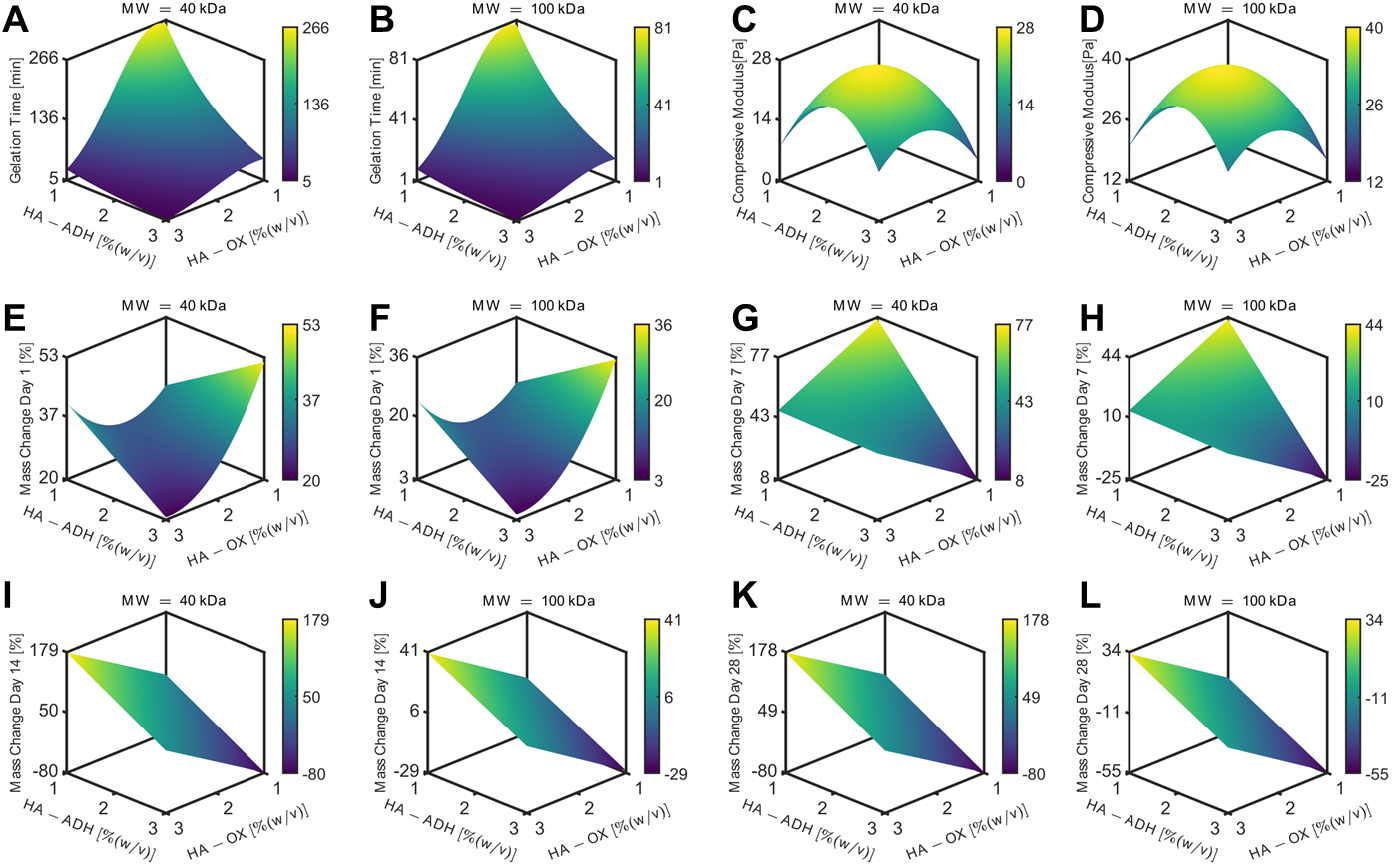
Three-dimensional response surfaces demonstrating the effect of input parameters on different output responses. Polymer weight percentages of HA-ADH and HA-Ox are on the x- and y-axes respectively, and output responses are on the z-axes. Responses for 40 kDa and 100 kDa HA hydrogels are plotted separately. The colored bars represent the magnitude of the response from low (blue) to high (yellow) for each response surface. Response surfaces for gelation time for A) 40 kDa HA and B) 100 kDa HA, compressive moduli for C) 40 kDa HA and D) 100 kDa HA, mass change at day 1 for E) 40 kDa HA and F) 100 kDa HA, mass change at day 7 for G) 40 kDa HA and H) 100 kDa HA, mass change at day 14 for I) 40 kDa HA and J) 100 kDa HA, and mass change at day 28 for, K) 40 kDa HA and L) 100 kDa HA.

Minimum and maximum mass changes occurred at the same polymer compositions for 100 kDa hydrogels on day 14, with a minimum at -26.63% and a maximum of 40.23% (**Figure 3J**). The response surfaces for 40 kDa and 100 kDa HA hydrogels at day 28 demonstrated linear dependences on HA-ADH and HA-Ox concentrations similar to response surfaces at day 14 (**Figure 3K, Figure 3L**). Overall, the response surfaces and ANOVA table demonstrate that gelation time, compressive modulus, and mass change at day 1 were dependent on interactions between HA-ADH concentration and HA-Ox concentration, while no interactions of the input parameters were observed with mass changes at the other time points.

### Optimal and Pessimal Hydrogel Predictions

Our overall objective was to use DoE to optimize HA hydrogels for protein delivery for bone regeneration. We aimed to formulate a hydrogel that exhibited a gelation time between 1-10 minutes, minimal mass change (i.e., 0%) over 28 days, and the maximum possible compressive modulus. D-optimal design generated predictive models describing how HA-ADH polymer concentration, HA-Ox polymer concentration, and HA molecular weight influenced compressive modulus, gelation time, and mass change.

To validate this model, we generated a hydrogel formulation that was predicted to optimize our target parameters (Optimal) as well as two hydrogels that were optimized with opposite objectives (Pessimal 1 and 2) (**Table 4**). More specifically, Pessimal 1 was set to degrade over 28 days, while Pessimal 2 was set to swell over 28 days. Our optimal formulation had a molecular weight of 100 kDa while both pessimal formulations had a molecular weight of 40 kDa. This may be due to the longer gelation times, softer compressive moduli, and larger mass changes over time that were observed in our 40 kDa hydrogels. The optimal formulation, HA-ADH_3.0_HA-OX_2.33_, met our targeted responses with a gelation time of 3.7 min (**Figure 4A**), compressive modulus of 62.1 Pa (**Figure 4B**), and minimal mass change ranging between -22.2 to 5.5% over 28 days (**Figure 4C**). Conversely, our Pessimal 1 formulation, HA-ADH_1.17_HA-OX_1.02_, displayed a gelation time of 262.2 min, compressive modulus of 23.1 Pa, and swelling up to 85.3% of its initial mass on day 4, followed by full degradation by day 21. The Pessimal 2 formulation, HA-ADH_1.70_HA-OX_1.80_ displayed a gelation time of 100 min, a compressive modulus of 70.6 Pa, and swelling up to 172.3% by day 28 with no degradation.

**Table 4.** Formulations for optimal and pessimal hydrogels.

|  | HA-ADH (%w/v) | HA-Ox (%w/v) | Molecular Weight |
| --- | --- | --- | --- |
| Optimal | 3.00 | 2.33 | 100 kDa |
| Pessimal 1 | 1.17 | 1.02 | 40 kDa |
| Pessimal 2 | 1.70 | 1.80 | 40 kDa |

**Figure 4.**
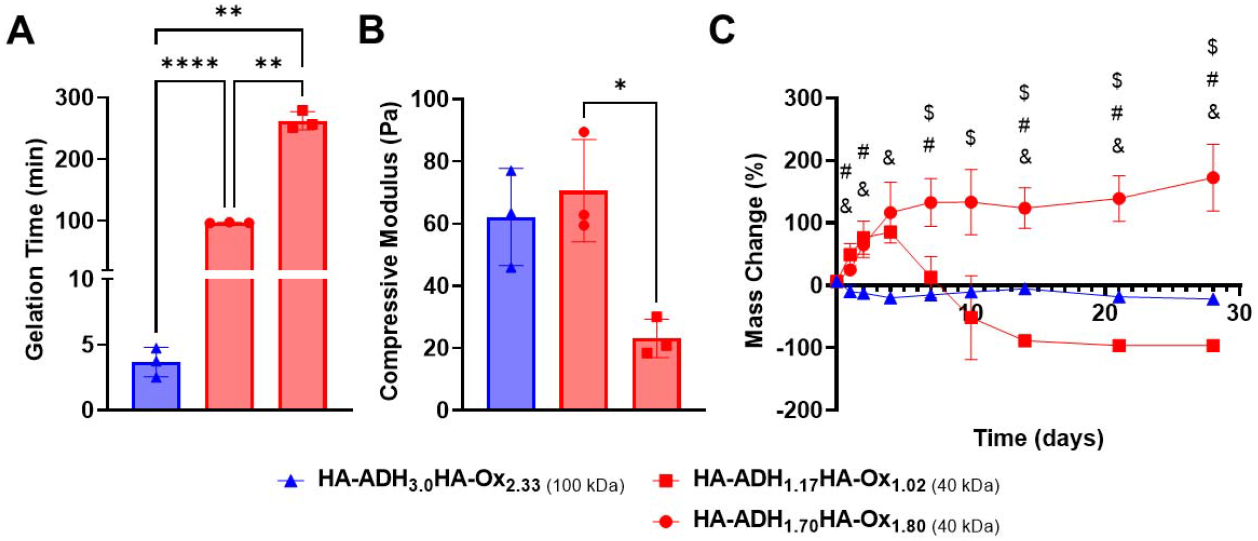
Validation of predictive models to optimize HA hydrogels. A) Gelation time of optimal (blue) and pessimal (red) hydrogels. One-way ANOVA, post-hoc Dunnett’s T3 multiple comparisons. n = 3; ** < p 0.01, **** p < 0.0001. B) Compressive modulus of optimal and pessimal hydrogels. One-way ANOVA, post-hoc Dunnett’s T3 multiple comparisons. n = 3; * p < 0.05. C) Mass change over 28 days of optimal and pessimal hydrogels. Two-way ANOVA, post-hoc Tukey’s multiple comparisons. n = 3; & p < 0.05 HA-ADH_3.0_HA-Ox_2.33 (100 kDa)_ vs. HA-ADH_1.17_HA-Ox_1.02 (40 kDa)_, # p < 0.05 HA-ADH_3.0_HA-Ox_2.33 (100 kDa)_ vs. HA-ADH_1.70_HA-Ox_1.80 (40 kDa)_, and $ p < 0.05 HA-ADH_1.17_HA-Ox_1.02 (40 kDa)_ vs. HA-ADH_1.70_HA-Ox_1.80 (40 kDa)_.

The gelation time of the optimal formulation was significantly lower than that of the pessimal formulations. Mass change for the optimal formulation was closer to the 0% target mass change at all timepoints, while both pessimal formulations exhibited initial swelling before either degrading completely or continuing to swell excessively. However, there were no significant differences between the compressive moduli of the optimal formulation and the two pessimal formulations. Thus, trying to independently target each of our desired physicochemical properties with our hydrogel formulation would not have been successful, highlighting the importance of using DoE to simultaneously optimize multiple output responses.

### BMP-2 Release from Affibody-Containing Hydrogels

As proof-of-concept of affinity-controlled protein delivery from our HA hydrogels, we modified HA-Ox to contain a norbornene functional group for bioconjugation of a previously discovered BMP-2-specific affibody with high affinity for BMP-2 (dissociation constant, K_D_ = 10.7 nM).(41) Affibodies are small proteins with 58 amino acids folded into a bundle of three α-helices.(56,57) Affibodies can be engineered to bind to different proteins with a range of affinities and can be used to release proteins at various rates by tuning the strength of the affinity interactions.(41,58,59) HA was functionalized with norbornene through BoC_2_O activated coupling (**Figure 5A**). The resulting product was then modified a second time by the previously described oxidation reaction, resulting in oxidized, norbornene-functionalized HA (HA-Nor-Ox; **Figure 5B**). Two times molar excess affibody to HA was then added to HA-Nor-Ox in the presence of a photoinitiator and exposed to light at 365nm to create affibody-conjugated HA-Nor-Ox (HA-Nor-Ox-Affibody). The average concentration of affibody conjugated to HA-Nor-Ox was 2.80±1.95 nmol/mg HA polymer (**Figure S3**). This modified polymer was crosslinked with HA-ADH to form a hydrazone-crosslinked, affibody-containing HA hydrogel.

**Figure 5.**
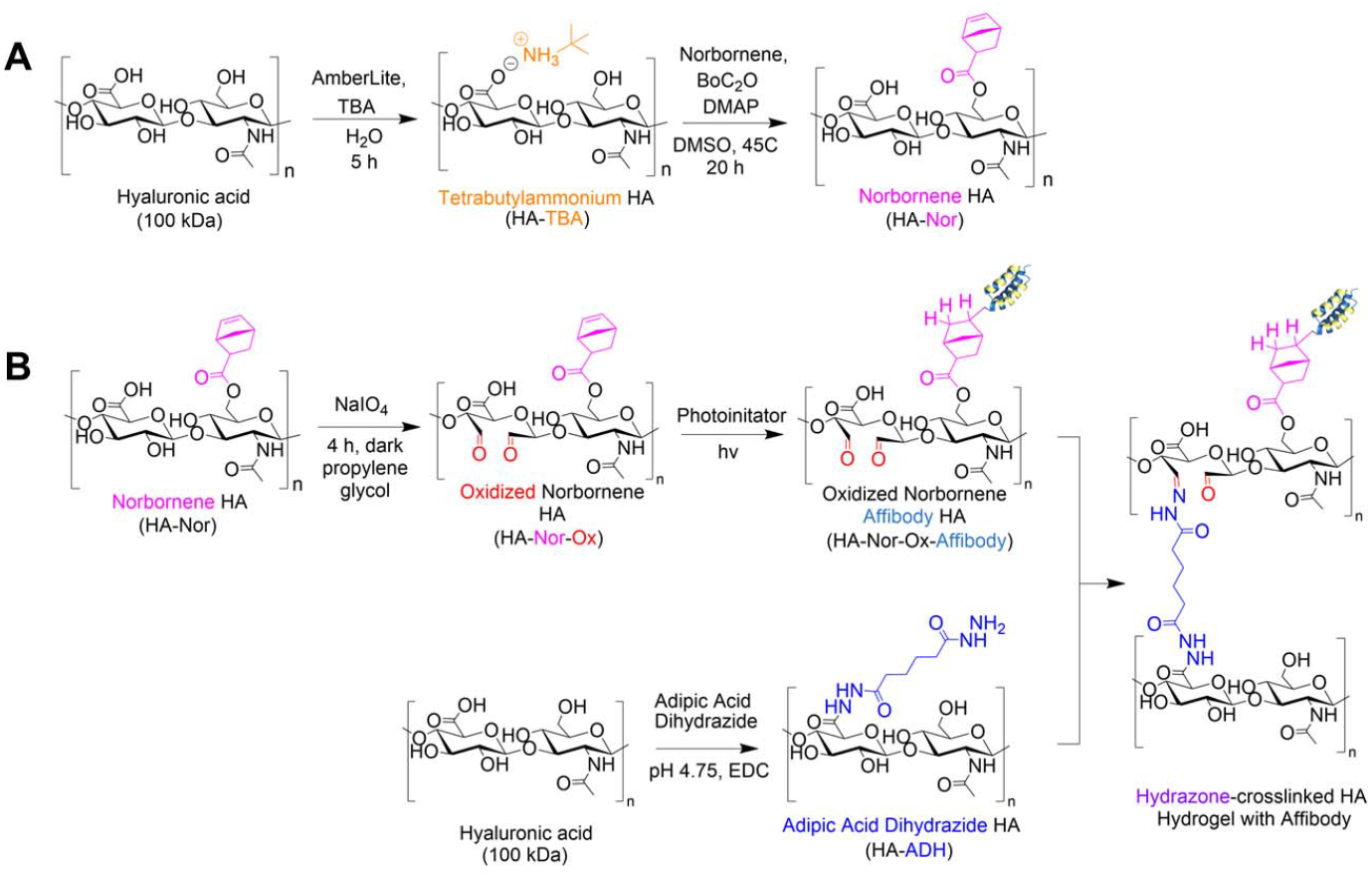
Synthesis of HA-Nor, HA-Nor-Ox, and HA-Nor-Ox-Affibody. A) HA was functionalized with a norbornene group through BoC_2_O activated coupling prior to oxidation resulting in HA-Nor-Ox. B) HA-Nor-Ox was then used to bioconjugate our BMP-2 affibody to the norbornene group through a photoinitiated reaction and thiol-ene click chemistry. The product was then crosslinked with HA-ADH to form a hydrazone-crosslinked hydrogel containing affibodies.

Release of BMP-2 was evaluated over 28 days from unmodified HA hydrogels or hydrogels conjugated with affibodies with high affinity and specificity for BMP-2 (dissociation constant, K_D_ = 10.7 nM).(41) Hydrogels were formed with BMP-2 and incubated in 1% w/v BSA in PBS. Aliquots were taken from the supernatant over 28 days to evaluate a release profile of BMP-2 from the HA hydrogel. The HA hydrogel with BMP-2 affibody reduced the release of BMP-2 compared to a HA hydrogel without affibody (**Figure 6A**). After 28 days, 11.7% of BMP-2 was released from the hydrogel without the affibodies, whereas 3.5% of BMP-2 was released from the hydrogel with the BMP-2-specific affibodies. **Figure 6B** depicts the cumulative fraction release at each time point (M_t_/M∞) as a function of the square root of time of the linear region of the BMP-2 release profile. The diffusivity (i.e., release rate) or the Fickian diffusion slope, k, was calculated using the slope of the cumulative fractional release (**Equation 3**), demonstrating that the release rate of BMP-2 was significantly slower in the affibody-containing hydrogel compared to the hydrogel without affibodies (**Figure 6C**).

**Figure 6.**
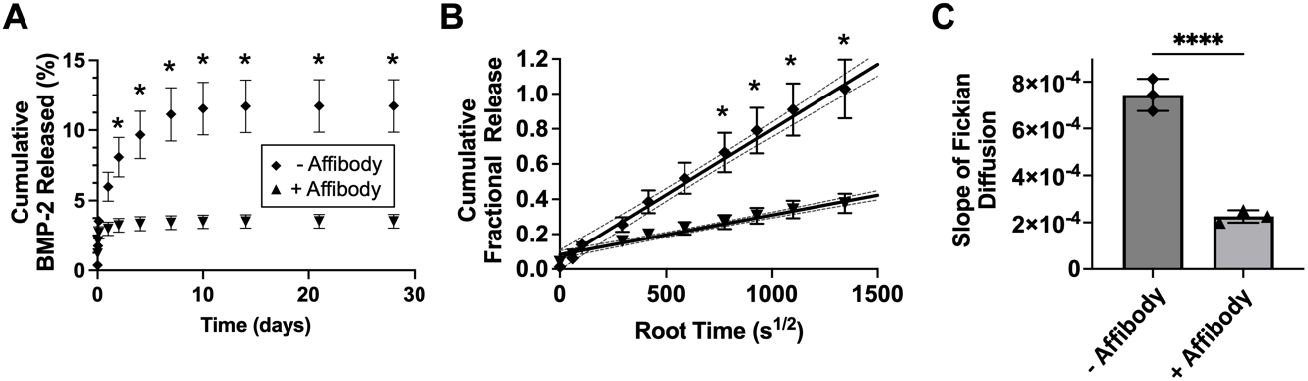
Release of BMP-2 from HA hydrogel. A) Cumulative release of BMP-2 from HA hydrogel without affibody (diamond) and with affibody (triangle). Two-way ANOVA, post-hoc Dunnett’s T3 multiple comparisons. n = 3; * p< 0.05. B) Linearized Release of BMP-2 affibody from hydrogels with and without affibody. Two-way ANOVA, post-hoc Dunnett’s T3 multiple comparisons. n = 3; * p< 0.05. C) Slope of linearized release of BMP-2 from HA hydrogels. One-way ANOVA, post-hoc Dunnett’s T3 multiple comparisons. n = 3; * p< 0.05, **** p < 0.0001.

## Conclusion

Although hydrazone chemistry is similar in many applications, each hydrogel requires distinct properties for a specific use. Designing an appropriate hydrogel for a biomedical application requires the consideration of many design criteria, such as gelation time, compressive modulus, and stability over time, that can be affected by many factors, including polymer molecular weight and concentration. In this work, we optimized hydrazone-crosslinked HA hydrogels for protein delivery for bone regeneration using DoE to better understand how various input parameters interacted to affect physicochemical properties, while reducing the number of experiments required to obtain this information. We generated models to describe how HA-ADH concentration, HA-Ox concentration, and HA molecular weight affected hydrogel compressive modulus, gelation time, and mass change over time. We identified an optimized hydrogel formulation that simultaneously maximized compressive modulus (62.1 Pa), achieved a target gelation time between 1-10 minutes (3.7 minutes), and was stable over 28 days with minimal mass change. We confirmed the validity of our model by evaluating two pessimal hydrogel formulations that were predicted to provide responses opposite to our optimal formulation. We also demonstrated that protein release from hydrazone-crosslinked HA hydrogels could be sustained over 28 days by conjugated BMP-2-specific affibodies to the hydrogels to control the release of BMP-2. Ultimately, hydrazone-crosslinked HA hydrogels are a promising platform for protein delivery and DoE is a valuable tool to optimize hydrogels for this application.

## Acknowledgements

We are grateful for funding from the Collins Medical Trust, NIH (R21-EB032112), and Wu Tsai Human Performance Alliance. JD is supported by a doctoral-level post-graduate scholarship (PGS-D) from the Natural Sciences and Engineering Council of Canada. EAM and JK were supported by the Knight Campus Undergraduate Scholars Program. MF was supported by the University of Oregon’s Summer Program in Undergraduate Research (SPUR) and Vice President for Research and Innovation (VPRI) Fellowship. We would like to thank Dr. Casey Check for his guidance on rheology. We would also like to thank Chandler Asnes and Payton Jefferis for their thoughtful feedback on this work.

## Conflict of Interest

A.N.G., J.D., V.R.S., and M.H.H. are authors on a pending patent application encompassing some of this work (U.S. Patent Application No. 18/340,754).

