## Supplemental Information for "Statistical Optimization of Hydrazone-Crosslinked Hyaluronic Acid Hydrogels for Protein Delivery"

*Degree of Modification*

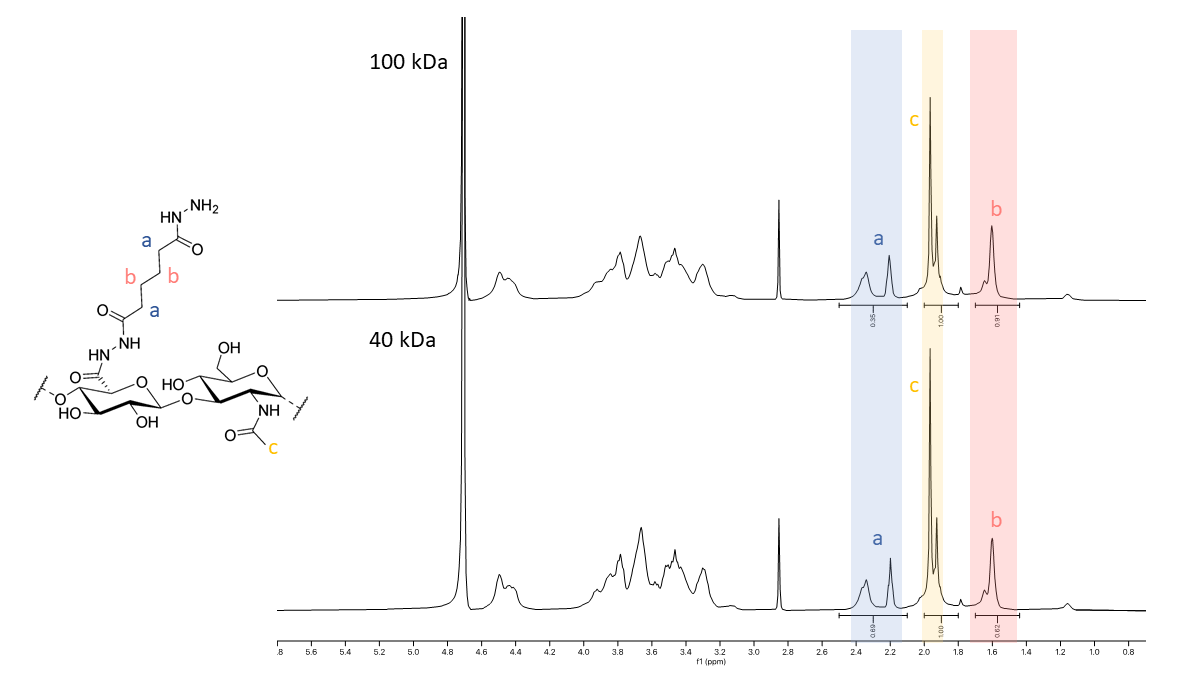

**Figure S1. ^1^H NMR of 40 kDa and 100 kDa HA-ADH***.* HA-ADH functionalization was confirmed by ^1^H NMR. The percentage of ADH functional groups per polymer chain was calculated by integrating the hydrogen peaks from the aliphatic chain on ADH (8H, a-b) and normalizing by the hydrogen peaks from the on the n-actyl group on HA (3H, c).

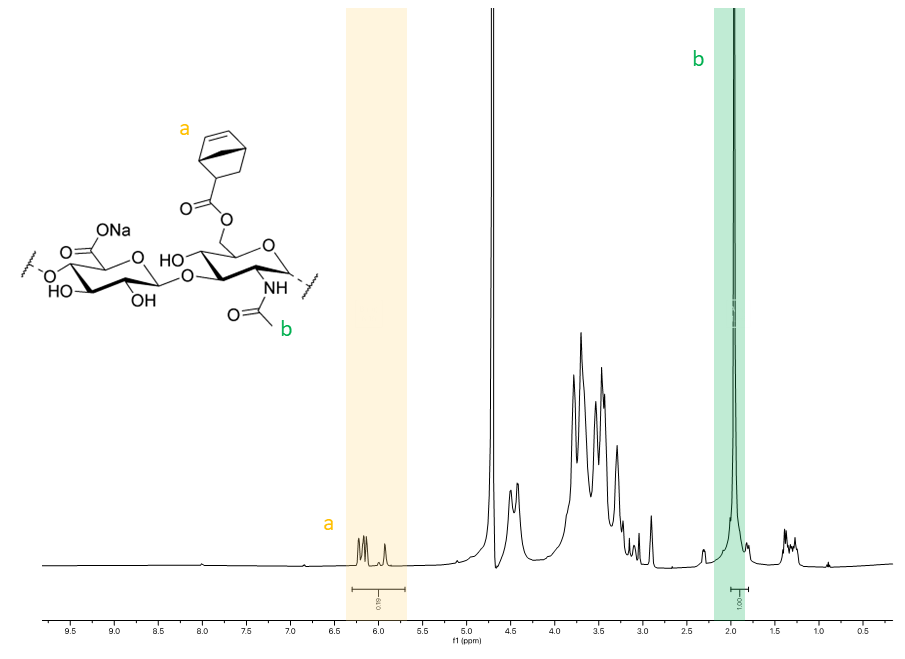

**Figure S2. ^1^H NMR of HA-Nor.** The functionalization of HA-Nor was confirmed by 1H NMR. The degree of modification or the percentage of norbornene groups per HA polymer chain was calculated by integrating the vinyl hydrogen peaks on the norbornene functional group (2H, a) and normalizing to the hydrogen peaks from the on the n-acetyl group on HA (3H, b).

|  | **%Degree of Modification** | | **%Yield** | |
| --- | --- | --- | --- | --- |
| **Modification** | **40 kDa** | **100 kDa** | **40 kDa** | **100 kDa** |
| **HA-Ox** | 22.5% | 56.0% | 42.0% | 68.3% |
|  | 25.9% | 35.5% | 21.8% | 89.2% |
|  | 34.0% | 32.6% | 27.6% | 83.4% |
|  | **27.5 ± 5.9%** | **41.4±12.8%** | **30.5±10.4%** | **80.3±10.8%** |
| **HA-ADH** | 69.3% | 45.1% | 45.1% | 44.9% |
|  | 67.5% | 52.7% | 52.7% | 46.4% |
|  | 66.8% | 40.6% | 40.6% | 49.5% |
|  | 66.9% | 63.7% | 52.8% | 46.6% |
|  | **67.6±1.2%** | **50.6±10.1%** | **47.8±6.0%** | **46.8±1.9%** |
| **HA-Nor** | - | 30% | - | 34% |
|  | - | 27% | - | 54% |
|  | - | 33.3% | - | 31% |
|  | **-** | **30.1±3.2%** | **-** | **38.7±12.5%** |

**Table S1. Degree of Modification and Yield for Functionalized HA Polymers.** A summary of the average degrees of modification and average yields for each functionalized HA polymer, including HA-Ox, HA-ADH, and HA-Nor.

*Gelation Time*

| **Gelation Time Significance** | | |
| --- | --- | --- |
| **Dunnett's T3 multiple comparisons test** | Summary | Adjusted P Value |
| HA-ADH_1.0_HA-Ox_1.0 (40 kDa)_ vs. HA-ADH_3.0_HA-Ox_1.0 (40 kDa)_ | **** | <0.0001 |
| HA-ADH_1.0_HA-Ox_1.0 (40 kDa)_ vs. HA-ADH_1.0_HA-Ox_3.0 (40 kDa)_ | *** | 0.0005 |
| HA-ADH_1.0_HA-Ox_1.0 (40 kDa)_ vs. HA-ADH_3.0_HA-Ox_3.0 (40 kDa)_ | **** | <0.0001 |
| HA-ADH_1.0_HA-Ox_1.0 (40 kDa)_ vs. HA-ADH_1.67_HA-Ox_1.0 (40 kDa)_ | ** | 0.0043 |
| HA-ADH_1.0_HA-Ox_1.0 (40 kDa)_ vs. HA-ADH_2.0_HA-Ox_2.0 (40 kDa)_ | ** | 0.0026 |
| HA-ADH_1.0_HA-Ox_1.0 (40 kDa)_ vs. HA-ADH_1.0_HA-Ox_1.0 (100 kDa)_ | **** | <0.0001 |
| HA-ADH_1.0_HA-Ox_1.0 (40 kDa)_ vs. HA-ADH_3.0_HA-Ox_1.0 (100 kDa)_ | **** | <0.0001 |
| HA-ADH_1.0_HA-Ox_1.0 (40 kDa)_ vs. HA-ADH_1.0_HA-Ox_3.0 (100 kDa)_ | *** | 0.0002 |
| HA-ADH_1.0_HA-Ox_1.0 (40 kDa)_ vs. HA-ADH_3.0_HA-Ox_3.0 (100 kDa)_ | **** | <0.0001 |
| HA-ADH_1.0_HA-Ox_1.0 (40 kDa)_ vs. HA-ADH_1.0_HA-Ox_1.67 (100 kDa)_ | **** | <0.0001 |
| HA-ADH_1.0_HA-Ox_1.0 (40 kDa)_ vs. HA-ADH_3.0_HA-Ox_2.33 (100 kDa)_ | **** | <0.0001 |
| HA-ADH_1.0_HA-Ox_1.0 (40 kDa)_ vs. HA-ADH_2.33_HA-Ox_1.0 (100 kDa)_ | *** | 0.0005 |
| HA-ADH_1.0_HA-Ox_1.0 (40 kDa)_ vs. HA-ADH_1.67_HA-Ox_3.0 (100 kDa)_ | *** | 0.0005 |
| HA-ADH_1.0_HA-Ox_1.0 (40 kDa)_ vs. HA-ADH_2.0_HA-Ox_2.0 (100 kDa)_ | **** | <0.0001 |
| HA-ADH_3.0_HA-Ox_1.0 (40 kDa)_ vs. HA-ADH_1.0_HA-Ox_3.0 (40 kDa)_ | ns | 0.3702 |
| HA-ADH_3.0_HA-Ox_1.0 (40 kDa)_ vs. HA-ADH_3.0_HA-Ox_3.0 (40 kDa)_ | *** | 0.0002 |
| HA-ADH_3.0_HA-Ox_1.0 (40 kDa)_ vs. HA-ADH_1.67_HA-Ox_1.0 (40 kDa)_ | ** | 0.0048 |
| HA-ADH_3.0_HA-Ox_1.0 (40 kDa)_ vs. HA-ADH_2.0_HA-Ox_2.0 (40 kDa)_ | ns | 0.0888 |
| HA-ADH_3.0_HA-Ox_1.0 (40 kDa)_ vs. HA-ADH_1.0_HA-Ox_1.0 (100 kDa)_ | *** | 0.0004 |
| HA-ADH_3.0_HA-Ox_1.0 (40 kDa)_ vs. HA-ADH_3.0_HA-Ox_1.0 (100 kDa)_ | ** | 0.0021 |
| HA-ADH_3.0_HA-Ox_1.0 (40 kDa)_ vs. HA-ADH_1.0_HA-Ox_3.0 (100 kDa)_ | ** | 0.0027 |
| HA-ADH_3.0_HA-Ox_1.0 (40 kDa)_ vs. HA-ADH_3.0_HA-Ox_3.0 (100 kDa)_ | ** | 0.0018 |
| HA-ADH_3.0_HA-Ox_1.0 (40 kDa)_ vs. HA-ADH_1.0_HA-Ox_1.67 (100 kDa)_ | ns | >0.9999 |
| HA-ADH_3.0_HA-Ox_1.0 (40 kDa)_ vs. HA-ADH_3.0_HA-Ox_2.33 (100 kDa)_ | *** | 0.0002 |
| HA-ADH_3.0_HA-Ox_1.0 (40 kDa)_ vs. HA-ADH_2.33_HA-Ox_1.0 (100 kDa)_ | * | 0.0335 |
| HA-ADH_3.0_HA-Ox_1.0 (40 kDa)_ vs. HA-ADH_1.67_HA-Ox_3.0 (100 kDa)_ | * | 0.0427 |
| HA-ADH_3.0_HA-Ox_1.0 (40 kDa)_ vs. HA-ADH_2.0_HA-Ox_2.0 (100 kDa)_ | **** | <0.0001 |
| HA-ADH_1.0_HA-Ox_3.0 (40 kDa)_ vs. HA-ADH_3.0_HA-Ox_3.0 (40 kDa)_ | * | 0.0246 |
| HA-ADH_1.0_HA-Ox_3.0 (40 kDa)_ vs. HA-ADH_1.67_HA-Ox_1.0 (40 kDa)_ | **** | <0.0001 |
| HA-ADH_1.0_HA-Ox_3.0 (40 kDa)_ vs. HA-ADH_2.0_HA-Ox_2.0 (40 kDa)_ | * | 0.0269 |
| HA-ADH_1.0_HA-Ox_3.0 (40 kDa)_ vs. HA-ADH_1.0_HA-Ox_1.0 (100 kDa)_ | * | 0.0359 |
| HA-ADH_1.0_HA-Ox_3.0 (40 kDa)_ vs. HA-ADH_3.0_HA-Ox_1.0 (100 kDa)_ | * | 0.0252 |
| HA-ADH_1.0_HA-Ox_3.0 (40 kDa)_ vs. HA-ADH_1.0_HA-Ox_3.0 (100 kDa)_ | * | 0.0305 |
| HA-ADH_1.0_HA-Ox_3.0 (40 kDa)_ vs. HA-ADH_3.0_HA-Ox_3.0 (100 kDa)_ | * | 0.0206 |
| HA-ADH_1.0_HA-Ox_3.0 (40 kDa)_ vs. HA-ADH_1.0_HA-Ox_1.67 (100 kDa)_ | ns | 0.3194 |
| HA-ADH_1.0_HA-Ox_3.0 (40 kDa)_ vs. HA-ADH_3.0_HA-Ox_2.33 (100 kDa)_ | * | 0.0243 |
| HA-ADH_1.0_HA-Ox_3.0 (40 kDa)_ vs. HA-ADH_2.33_HA-Ox_1.0 (100 kDa)_ | * | 0.0204 |
| HA-ADH_1.0_HA-Ox_3.0 (40 kDa)_ vs. HA-ADH_1.67_HA-Ox_3.0 (100 kDa)_ | * | 0.0327 |
| HA-ADH_1.0_HA-Ox_3.0 (40 kDa)_ vs. HA-ADH_2.0_HA-Ox_2.0 (100 kDa)_ | * | 0.0465 |
| HA-ADH_3.0_HA-Ox_3.0 (40 kDa)_ vs. HA-ADH_1.67_HA-Ox_1.0 (40 kDa)_ | ** | 0.0024 |
| HA-ADH_3.0_HA-Ox_3.0 (40 kDa)_ vs. HA-ADH_2.0_HA-Ox_2.0 (40 kDa)_ | * | 0.0173 |
| HA-ADH_3.0_HA-Ox_3.0 (40 kDa)_ vs. HA-ADH_1.0_HA-Ox_1.0 (100 kDa)_ | **** | <0.0001 |
| HA-ADH_3.0_HA-Ox_3.0 (40 kDa)_ vs. HA-ADH_3.0_HA-Ox_1.0 (100 kDa)_ | ns | 0.8765 |
| HA-ADH_3.0_HA-Ox_3.0 (40 kDa)_ vs. HA-ADH_1.0_HA-Ox_3.0 (100 kDa)_ | ns | 0.0698 |
| HA-ADH_3.0_HA-Ox_3.0 (40 kDa)_ vs. HA-ADH_3.0_HA-Ox_3.0 (100 kDa)_ | * | 0.0367 |
| HA-ADH_3.0_HA-Ox_3.0 (40 kDa)_ vs. HA-ADH_1.0_HA-Ox_1.67 (100 kDa)_ | **** | <0.0001 |
| HA-ADH_3.0_HA-Ox_3.0 (40 kDa)_ vs. HA-ADH_3.0_HA-Ox_2.33 (100 kDa)_ | ns | >0.9999 |
| HA-ADH_3.0_HA-Ox_3.0 (40 kDa)_ vs. HA-ADH_2.33_HA-Ox_1.0 (100 kDa)_ | ns | 0.2687 |
| HA-ADH_3.0_HA-Ox_3.0 (40 kDa)_ vs. HA-ADH_1.67_HA-Ox_3.0 (100 kDa)_ | ns | 0.2078 |
| HA-ADH_3.0_HA-Ox_3.0 (40 kDa)_ vs. HA-ADH_2.0_HA-Ox_2.0 (100 kDa)_ | **** | <0.0001 |
| HA-ADH_1.67_HA-Ox_1.0 (40 kDa)_ vs. HA-ADH_2.0_HA-Ox_2.0 (40 kDa)_ | ** | 0.0013 |
| HA-ADH_1.67_HA-Ox_1.0 (40 kDa)_ vs. HA-ADH_1.0_HA-Ox_1.0 (100 kDa)_ | ** | 0.0072 |
| HA-ADH_1.67_HA-Ox_1.0 (40 kDa)_ vs. HA-ADH_3.0_HA-Ox_1.0 (100 kDa)_ | ** | 0.0024 |
| HA-ADH_1.67_HA-Ox_1.0 (40 kDa)_ vs. HA-ADH_1.0_HA-Ox_3.0 (100 kDa)_ | *** | 0.0003 |
| HA-ADH_1.67_HA-Ox_1.0 (40 kDa)_ vs. HA-ADH_3.0_HA-Ox_3.0 (100 kDa)_ | ** | 0.0023 |
| HA-ADH_1.67_HA-Ox_1.0 (40 kDa)_ vs. HA-ADH_1.0_HA-Ox_1.67 (100 kDa)_ | ** | 0.0047 |
| HA-ADH_1.67_HA-Ox_1.0 (40 kDa)_ vs. HA-ADH_3.0_HA-Ox_2.33 (100 kDa)_ | ** | 0.0024 |
| HA-ADH_1.67_HA-Ox_1.0 (40 kDa)_ vs. HA-ADH_2.33_HA-Ox_1.0 (100 kDa)_ | **** | <0.0001 |
| HA-ADH_1.67_HA-Ox_1.0 (40 kDa)_ vs. HA-ADH_1.67_HA-Ox_3.0 (100 kDa)_ | **** | <0.0001 |
| HA-ADH_1.67_HA-Ox_1.0 (40 kDa)_ vs. HA-ADH_2.0_HA-Ox_2.0 (100 kDa)_ | ** | 0.0033 |
| HA-ADH_2.0_HA-Ox_2.0 (40 kDa)_ vs. HA-ADH_1.0_HA-Ox_1.0 (100 kDa)_ | ns | 0.4885 |
| HA-ADH_2.0_HA-Ox_2.0 (40 kDa)_ vs. HA-ADH_3.0_HA-Ox_1.0 (100 kDa)_ | * | 0.0176 |
| HA-ADH_2.0_HA-Ox_2.0 (40 kDa)_ vs. HA-ADH_1.0_HA-Ox_3.0 (100 kDa)_ | * | 0.027 |
| HA-ADH_2.0_HA-Ox_2.0 (40 kDa)_ vs. HA-ADH_3.0_HA-Ox_3.0 (100 kDa)_ | * | 0.016 |
| HA-ADH_2.0_HA-Ox_2.0 (40 kDa)_ vs. HA-ADH_1.0_HA-Ox_1.67 (100 kDa)_ | ns | 0.0889 |
| HA-ADH_2.0_HA-Ox_2.0 (40 kDa)_ vs. HA-ADH_3.0_HA-Ox_2.33 (100 kDa)_ | * | 0.0172 |
| HA-ADH_2.0_HA-Ox_2.0 (40 kDa)_ vs. HA-ADH_2.33_HA-Ox_1.0 (100 kDa)_ | ** | 0.0074 |
| HA-ADH_2.0_HA-Ox_2.0 (40 kDa)_ vs. HA-ADH_1.67_HA-Ox_3.0 (100 kDa)_ | ** | 0.0086 |
| HA-ADH_2.0_HA-Ox_2.0 (40 kDa)_ vs. HA-ADH_2.0_HA-Ox_2.0 (100 kDa)_ | * | 0.0297 |
| HA-ADH_1.0_HA-Ox_1.0 (100 kDa)_ vs. HA-ADH_3.0_HA-Ox_1.0 (100 kDa)_ | *** | 0.0008 |
| HA-ADH_1.0_HA-Ox_1.0 (100 kDa)_ vs. HA-ADH_1.0_HA-Ox_3.0 (100 kDa)_ | *** | 0.0005 |
| HA-ADH_1.0_HA-Ox_1.0 (100 kDa)_ vs. HA-ADH_3.0_HA-Ox_3.0 (100 kDa)_ | *** | 0.0007 |
| HA-ADH_1.0_HA-Ox_1.0 (100 kDa)_ vs. HA-ADH_1.0_HA-Ox_1.67 (100 kDa)_ | ** | 0.0012 |
| HA-ADH_1.0_HA-Ox_1.0 (100 kDa)_ vs. HA-ADH_3.0_HA-Ox_2.33 (100 kDa)_ | **** | <0.0001 |
| HA-ADH_1.0_HA-Ox_1.0 (100 kDa)_ vs. HA-ADH_2.33_HA-Ox_1.0 (100 kDa)_ | * | 0.0122 |
| HA-ADH_1.0_HA-Ox_1.0 (100 kDa)_ vs. HA-ADH_1.67_HA-Ox_3.0 (100 kDa)_ | * | 0.0147 |
| HA-ADH_1.0_HA-Ox_1.0 (100 kDa)_ vs. HA-ADH_2.0_HA-Ox_2.0 (100 kDa)_ | **** | <0.0001 |
| HA-ADH_3.0_HA-Ox_1.0 (100 kDa)_ vs. HA-ADH_1.0_HA-Ox_3.0 (100 kDa)_ | ns | 0.0748 |
| HA-ADH_3.0_HA-Ox_1.0 (100 kDa)_ vs. HA-ADH_3.0_HA-Ox_3.0 (100 kDa)_ | ** | 0.0023 |
| HA-ADH_3.0_HA-Ox_1.0 (100 kDa)_ vs. HA-ADH_1.0_HA-Ox_1.67 (100 kDa)_ | **** | <0.0001 |
| HA-ADH_3.0_HA-Ox_1.0 (100 kDa)_ vs. HA-ADH_3.0_HA-Ox_2.33 (100 kDa)_ | ns | 0.657 |
| HA-ADH_3.0_HA-Ox_1.0 (100 kDa)_ vs. HA-ADH_2.33_HA-Ox_1.0 (100 kDa)_ | ns | 0.2964 |
| HA-ADH_3.0_HA-Ox_1.0 (100 kDa)_ vs. HA-ADH_1.67_HA-Ox_3.0 (100 kDa)_ | ns | 0.2257 |
| HA-ADH_3.0_HA-Ox_1.0 (100 kDa)_ vs. HA-ADH_2.0_HA-Ox_2.0 (100 kDa)_ | **** | <0.0001 |
| HA-ADH_1.0_HA-Ox_3.0 (100 kDa)_ vs. HA-ADH_3.0_HA-Ox_3.0 (100 kDa)_ | * | 0.0446 |
| HA-ADH_1.0_HA-Ox_3.0 (100 kDa)_ vs. HA-ADH_1.0_HA-Ox_1.67 (100 kDa)_ | * | 0.0121 |
| HA-ADH_1.0_HA-Ox_3.0 (100 kDa)_ vs. HA-ADH_3.0_HA-Ox_2.33 (100 kDa)_ | ns | 0.0674 |
| HA-ADH_1.0_HA-Ox_3.0 (100 kDa)_ vs. HA-ADH_2.33_HA-Ox_1.0 (100 kDa)_ | ns | 0.9998 |
| HA-ADH_1.0_HA-Ox_3.0 (100 kDa)_ vs. HA-ADH_1.67_HA-Ox_3.0 (100 kDa)_ | ns | >0.9999 |
| HA-ADH_1.0_HA-Ox_3.0 (100 kDa)_ vs. HA-ADH_2.0_HA-Ox_2.0 (100 kDa)_ | ns | 0.5425 |
| HA-ADH_3.0_HA-Ox_3.0 (100 kDa)_ vs. HA-ADH_1.0_HA-Ox_1.67 (100 kDa)_ | **** | <0.0001 |
| HA-ADH_3.0_HA-Ox_3.0 (100 kDa)_ vs. HA-ADH_3.0_HA-Ox_2.33 (100 kDa)_ | * | 0.0417 |
| HA-ADH_3.0_HA-Ox_3.0 (100 kDa)_ vs. HA-ADH_2.33_HA-Ox_1.0 (100 kDa)_ | ns | 0.1714 |
| HA-ADH_3.0_HA-Ox_3.0 (100 kDa)_ vs. HA-ADH_1.67_HA-Ox_3.0 (100 kDa)_ | ns | 0.141 |
| HA-ADH_3.0_HA-Ox_3.0 (100 kDa)_ vs. HA-ADH_2.0_HA-Ox_2.0 (100 kDa)_ | **** | <0.0001 |
| HA-ADH_1.0_HA-Ox_1.67 (100 kDa)_ vs. HA-ADH_3.0_HA-Ox_2.33 (100 kDa)_ | **** | <0.0001 |
| HA-ADH_1.0_HA-Ox_1.67 (100 kDa)_ vs. HA-ADH_2.33_HA-Ox_1.0 (100 kDa)_ | * | 0.0303 |
| HA-ADH_1.0_HA-Ox_1.67 (100 kDa)_ vs. HA-ADH_1.67_HA-Ox_3.0 (100 kDa)_ | * | 0.0389 |
| HA-ADH_1.0_HA-Ox_1.67 (100 kDa)_ vs. HA-ADH_2.0_HA-Ox_2.0 (100 kDa)_ | **** | <0.0001 |
| HA-ADH_3.0_HA-Ox_2.33 (100 kDa)_ vs. HA-ADH_2.33_HA-Ox_1.0 (100 kDa)_ | ns | 0.2597 |
| HA-ADH_3.0_HA-Ox_2.33 (100 kDa)_ vs. HA-ADH_1.67_HA-Ox_3.0 (100 kDa)_ | ns | 0.2018 |
| HA-ADH_3.0_HA-Ox_2.33 (100 kDa)_ vs. HA-ADH_2.0_HA-Ox_2.0 (100 kDa)_ | **** | <0.0001 |
| HA-ADH_2.33_HA-Ox_1.0 (100 kDa)_ vs. HA-ADH_1.67_HA-Ox_3.0 (100 kDa)_ | ns | >0.9999 |
| HA-ADH_2.33_HA-Ox_1.0 (100 kDa)_ vs. HA-ADH_2.0_HA-Ox_2.0 (100 kDa)_ | ns | 0.5742 |
| HA-ADH_1.67_HA-Ox_3.0 (100 kDa)_ vs. HA-ADH_2.0_HA-Ox_2.0 (100 kDa)_ | ns | 0.9285 |

**Table S2. Summary of significance for 40 kDa HA gelation times and 100 kDa HA gelation time.** A summary of significance for the gelation times for both 40 kDa and 100 kDa is listed, a one-way ANOVA post-hoc Dunnett’s T3 multiple comparisons was performed. n = 3; * p < 0.05, ** < p 0.01, *** p < 0.001, **** p < 0.0001.

*Compressive Modulus*

| **Compressive Modulus Significance** | | |
| --- | --- | --- |
| **Dunnett's T3 multiple comparisons test** | Summary | Adjusted P Value |
| HA-ADH_1.0_HA-Ox_1.0 (40 kDa)_ vs. HA-ADH_3.0_HA-Ox_1.0 (40 kDa)_ | ns | 0.9591 |
| HA-ADH_1.0_HA-Ox_1.0 (40 kDa)_ vs. HA-ADH_1.0_HA-Ox_3.0 (40 kDa)_ | ns | >0.9999 |
| HA-ADH_1.0_HA-Ox_1.0 (40 kDa)_ vs. HA-ADH_3.0_HA-Ox_3.0 (40 kDa)_ | ns | 0.4543 |
| HA-ADH_1.0_HA-Ox_1.0 (40 kDa)_ vs. HA-ADH_1.67_HA-Ox_1.0 (40 kDa)_ | ns | 0.5122 |
| HA-ADH_1.0_HA-Ox_1.0 (40 kDa)_ vs. HA-ADH_2.0_HA-Ox_2.0 (40 kDa)_ | ns | 0.8766 |
| HA-ADH_1.0_HA-Ox_1.0 (40 kDa)_ vs. HA-ADH_1.0_HA-Ox_1.0 (100 kDa)_ | * | 0.0297 |
| HA-ADH_1.0_HA-Ox_1.0 (40 kDa)_ vs. HA-ADH_3.0_HA-Ox_1.0 (100 kDa)_ | ns | 0.3815 |
| HA-ADH_1.0_HA-Ox_1.0 (40 kDa)_ vs. HA-ADH_1.0_HA-Ox_3.0 (100 kDa)_ | ns | 0.5412 |
| HA-ADH_1.0_HA-Ox_1.0 (40 kDa)_ vs. HA-ADH_3.0_HA-Ox_3.0 (100 kDa)_ | ns | 0.1398 |
| HA-ADH_1.0_HA-Ox_1.0 (40 kDa)_ vs. HA-ADH_1.0_HA-Ox_1.67 (100 kDa)_ | ns | 0.2201 |
| HA-ADH_1.0_HA-Ox_1.0 (40 kDa)_ vs. HA-ADH_3.0_HA-Ox_2.33 (100 kDa)_ | ** | 0.0065 |
| HA-ADH_1.0_HA-Ox_1.0 (40 kDa)_ vs. HA-ADH_2.33_HA-Ox_1.0 (100 kDa)_ | * | 0.0296 |
| HA-ADH_1.0_HA-Ox_1.0 (40 kDa)_ vs. HA-ADH_1.67_HA-Ox_3.0 (100 kDa)_ | ns | 0.1101 |
| HA-ADH_1.0_HA-Ox_1.0 (40 kDa)_ vs. HA-ADH_2.0_HA-Ox_2.0 (100 kDa)_ | *** | 0.0001 |
| HA-ADH_3.0_HA-Ox_1.0 (40 kDa)_ vs. HA-ADH_1.0_HA-Ox_3.0 (40 kDa)_ | ns | 0.9702 |
| HA-ADH_3.0_HA-Ox_1.0 (40 kDa)_ vs. HA-ADH_3.0_HA-Ox_3.0 (40 kDa)_ | ns | 0.5643 |
| HA-ADH_3.0_HA-Ox_1.0 (40 kDa)_ vs. HA-ADH_1.67_HA-Ox_1.0 (40 kDa)_ | ns | 0.9999 |
| HA-ADH_3.0_HA-Ox_1.0 (40 kDa)_ vs. HA-ADH_2.0_HA-Ox_2.0 (40 kDa)_ | ns | 0.9989 |
| HA-ADH_3.0_HA-Ox_1.0 (40 kDa)_ vs. HA-ADH_1.0_HA-Ox_1.0 (100 kDa)_ | ns | 0.144 |
| HA-ADH_3.0_HA-Ox_1.0 (40 kDa)_ vs. HA-ADH_3.0_HA-Ox_1.0 (100 kDa)_ | ns | 0.431 |
| HA-ADH_3.0_HA-Ox_1.0 (40 kDa)_ vs. HA-ADH_1.0_HA-Ox_3.0 (100 kDa)_ | ns | 0.6178 |
| HA-ADH_3.0_HA-Ox_1.0 (40 kDa)_ vs. HA-ADH_3.0_HA-Ox_3.0 (100 kDa)_ | ns | 0.1057 |
| HA-ADH_3.0_HA-Ox_1.0 (40 kDa)_ vs. HA-ADH_1.0_HA-Ox_1.67 (100 kDa)_ | ns | 0.2891 |
| HA-ADH_3.0_HA-Ox_1.0 (40 kDa)_ vs. HA-ADH_3.0_HA-Ox_2.33 (100 kDa)_ | ** | 0.0036 |
| HA-ADH_3.0_HA-Ox_1.0 (40 kDa)_ vs. HA-ADH_2.33_HA-Ox_1.0 (100 kDa)_ | * | 0.0491 |
| HA-ADH_3.0_HA-Ox_1.0 (40 kDa)_ vs. HA-ADH_1.67_HA-Ox_3.0 (100 kDa)_ | ns | 0.0833 |
| HA-ADH_3.0_HA-Ox_1.0 (40 kDa)_ vs. HA-ADH_2.0_HA-Ox_2.0 (100 kDa)_ | *** | 0.0002 |
| HA-ADH_1.0_HA-Ox_3.0 (40 kDa)_ vs. HA-ADH_3.0_HA-Ox_3.0 (40 kDa)_ | ns | 0.4708 |
| HA-ADH_1.0_HA-Ox_3.0 (40 kDa)_ vs. HA-ADH_1.67_HA-Ox_1.0 (40 kDa)_ | ns | 0.5468 |
| HA-ADH_1.0_HA-Ox_3.0 (40 kDa)_ vs. HA-ADH_2.0_HA-Ox_2.0 (40 kDa)_ | ns | 0.9131 |
| HA-ADH_1.0_HA-Ox_3.0 (40 kDa)_ vs. HA-ADH_1.0_HA-Ox_1.0 (100 kDa)_ | * | 0.0434 |
| HA-ADH_1.0_HA-Ox_3.0 (40 kDa)_ vs. HA-ADH_3.0_HA-Ox_1.0 (100 kDa)_ | ns | 0.3904 |
| HA-ADH_1.0_HA-Ox_3.0 (40 kDa)_ vs. HA-ADH_1.0_HA-Ox_3.0 (100 kDa)_ | ns | 0.5559 |
| HA-ADH_1.0_HA-Ox_3.0 (40 kDa)_ vs. HA-ADH_3.0_HA-Ox_3.0 (100 kDa)_ | ns | 0.1407 |
| HA-ADH_1.0_HA-Ox_3.0 (40 kDa)_ vs. HA-ADH_1.0_HA-Ox_1.67 (100 kDa)_ | ns | 0.2246 |
| HA-ADH_1.0_HA-Ox_3.0 (40 kDa)_ vs. HA-ADH_3.0_HA-Ox_2.33 (100 kDa)_ | * | 0.0243 |
| HA-ADH_1.0_HA-Ox_3.0 (40 kDa)_ vs. HA-ADH_2.33_HA-Ox_1.0 (100 kDa)_ | ns | 0.0734 |
| HA-ADH_1.0_HA-Ox_3.0 (40 kDa)_ vs. HA-ADH_1.67_HA-Ox_3.0 (100 kDa)_ | ns | 0.1083 |
| HA-ADH_1.0_HA-Ox_3.0 (40 kDa)_ vs. HA-ADH_2.0_HA-Ox_2.0 (100 kDa)_ | *** | 0.0003 |
| HA-ADH_3.0_HA-Ox_3.0 (40 kDa)_ vs. HA-ADH_1.67_HA-Ox_1.0 (40 kDa)_ | ns | 0.638 |
| HA-ADH_3.0_HA-Ox_3.0 (40 kDa)_ vs. HA-ADH_2.0_HA-Ox_2.0 (40 kDa)_ | ns | 0.8503 |
| HA-ADH_3.0_HA-Ox_3.0 (40 kDa)_ vs. HA-ADH_1.0_HA-Ox_1.0 (100 kDa)_ | ns | 0.9537 |
| HA-ADH_3.0_HA-Ox_3.0 (40 kDa)_ vs. HA-ADH_3.0_HA-Ox_1.0 (100 kDa)_ | ns | 0.9545 |
| HA-ADH_3.0_HA-Ox_3.0 (40 kDa)_ vs. HA-ADH_1.0_HA-Ox_3.0 (100 kDa)_ | ns | >0.9999 |
| HA-ADH_3.0_HA-Ox_3.0 (40 kDa)_ vs. HA-ADH_3.0_HA-Ox_3.0 (100 kDa)_ | ns | 0.9992 |
| HA-ADH_3.0_HA-Ox_3.0 (40 kDa)_ vs. HA-ADH_1.0_HA-Ox_1.67 (100 kDa)_ | ns | >0.9999 |
| HA-ADH_3.0_HA-Ox_3.0 (40 kDa)_ vs. HA-ADH_3.0_HA-Ox_2.33 (100 kDa)_ | ns | 0.7687 |
| HA-ADH_3.0_HA-Ox_3.0 (40 kDa)_ vs. HA-ADH_2.33_HA-Ox_1.0 (100 kDa)_ | ns | 0.9996 |
| HA-ADH_3.0_HA-Ox_3.0 (40 kDa)_ vs. HA-ADH_1.67_HA-Ox_3.0 (100 kDa)_ | ns | >0.9999 |
| HA-ADH_3.0_HA-Ox_3.0 (40 kDa)_ vs. HA-ADH_2.0_HA-Ox_2.0 (100 kDa)_ | ns | 0.2458 |
| HA-ADH_1.67_HA-Ox_1.0 (40 kDa)_ vs. HA-ADH_2.0_HA-Ox_2.0 (40 kDa)_ | ns | >0.9999 |
| HA-ADH_1.67_HA-Ox_1.0 (40 kDa)_ vs. HA-ADH_1.0_HA-Ox_1.0 (100 kDa)_ | ns | 0.2323 |
| HA-ADH_1.67_HA-Ox_1.0 (40 kDa)_ vs. HA-ADH_3.0_HA-Ox_1.0 (100 kDa)_ | ns | 0.4638 |
| HA-ADH_1.67_HA-Ox_1.0 (40 kDa)_ vs. HA-ADH_1.0_HA-Ox_3.0 (100 kDa)_ | ns | 0.6686 |
| HA-ADH_1.67_HA-Ox_1.0 (40 kDa)_ vs. HA-ADH_3.0_HA-Ox_3.0 (100 kDa)_ | ns | 0.203 |
| HA-ADH_1.67_HA-Ox_1.0 (40 kDa)_ vs. HA-ADH_1.0_HA-Ox_1.67 (100 kDa)_ | ns | 0.3314 |
| HA-ADH_1.67_HA-Ox_1.0 (40 kDa)_ vs. HA-ADH_3.0_HA-Ox_2.33 (100 kDa)_ | ** | 0.0039 |
| HA-ADH_1.67_HA-Ox_1.0 (40 kDa)_ vs. HA-ADH_2.33_HA-Ox_1.0 (100 kDa)_ | ns | 0.057 |
| HA-ADH_1.67_HA-Ox_1.0 (40 kDa)_ vs. HA-ADH_1.67_HA-Ox_3.0 (100 kDa)_ | ns | 0.0997 |
| HA-ADH_1.67_HA-Ox_1.0 (40 kDa)_ vs. HA-ADH_2.0_HA-Ox_2.0 (100 kDa)_ | *** | 0.0003 |
| HA-ADH_2.0_HA-Ox_2.0 (40 kDa)_ vs. HA-ADH_1.0_HA-Ox_1.0 (100 kDa)_ | ns | 0.9845 |
| HA-ADH_2.0_HA-Ox_2.0 (40 kDa)_ vs. HA-ADH_3.0_HA-Ox_1.0 (100 kDa)_ | ns | 0.5079 |
| HA-ADH_2.0_HA-Ox_2.0 (40 kDa)_ vs. HA-ADH_1.0_HA-Ox_3.0 (100 kDa)_ | ns | 0.7857 |
| HA-ADH_2.0_HA-Ox_2.0 (40 kDa)_ vs. HA-ADH_3.0_HA-Ox_3.0 (100 kDa)_ | ns | 0.2589 |
| HA-ADH_2.0_HA-Ox_2.0 (40 kDa)_ vs. HA-ADH_1.0_HA-Ox_1.67 (100 kDa)_ | ns | 0.5083 |
| HA-ADH_2.0_HA-Ox_2.0 (40 kDa)_ vs. HA-ADH_3.0_HA-Ox_2.33 (100 kDa)_ | ns | 0.0877 |
| HA-ADH_2.0_HA-Ox_2.0 (40 kDa)_ vs. HA-ADH_2.33_HA-Ox_1.0 (100 kDa)_ | ns | 0.1852 |
| HA-ADH_2.0_HA-Ox_2.0 (40 kDa)_ vs. HA-ADH_1.67_HA-Ox_3.0 (100 kDa)_ | ns | 0.2961 |
| HA-ADH_2.0_HA-Ox_2.0 (40 kDa)_ vs. HA-ADH_2.0_HA-Ox_2.0 (100 kDa)_ | ** | 0.0047 |
| HA-ADH_1.0_HA-Ox_1.0 (100 kDa)_ vs. HA-ADH_3.0_HA-Ox_1.0 (100 kDa)_ | ns | 0.6264 |
| HA-ADH_1.0_HA-Ox_1.0 (100 kDa)_ vs. HA-ADH_1.0_HA-Ox_3.0 (100 kDa)_ | ns | 0.8819 |
| HA-ADH_1.0_HA-Ox_1.0 (100 kDa)_ vs. HA-ADH_3.0_HA-Ox_3.0 (100 kDa)_ | ns | 0.3723 |
| HA-ADH_1.0_HA-Ox_1.0 (100 kDa)_ vs. HA-ADH_1.0_HA-Ox_1.67 (100 kDa)_ | ns | 0.6428 |
| HA-ADH_1.0_HA-Ox_1.0 (100 kDa)_ vs. HA-ADH_3.0_HA-Ox_2.33 (100 kDa)_ | * | 0.0263 |
| HA-ADH_1.0_HA-Ox_1.0 (100 kDa)_ vs. HA-ADH_2.33_HA-Ox_1.0 (100 kDa)_ | ns | 0.1541 |
| HA-ADH_1.0_HA-Ox_1.0 (100 kDa)_ vs. HA-ADH_1.67_HA-Ox_3.0 (100 kDa)_ | ns | 0.2902 |
| HA-ADH_1.0_HA-Ox_1.0 (100 kDa)_ vs. HA-ADH_2.0_HA-Ox_2.0 (100 kDa)_ | *** | 0.001 |
| HA-ADH_3.0_HA-Ox_1.0 (100 kDa)_ vs. HA-ADH_1.0_HA-Ox_3.0 (100 kDa)_ | ns | >0.9999 |
| HA-ADH_3.0_HA-Ox_1.0 (100 kDa)_ vs. HA-ADH_3.0_HA-Ox_3.0 (100 kDa)_ | ns | 0.9991 |
| HA-ADH_3.0_HA-Ox_1.0 (100 kDa)_ vs. HA-ADH_1.0_HA-Ox_1.67 (100 kDa)_ | ns | 0.9743 |
| HA-ADH_3.0_HA-Ox_1.0 (100 kDa)_ vs. HA-ADH_3.0_HA-Ox_2.33 (100 kDa)_ | ns | >0.9999 |
| HA-ADH_3.0_HA-Ox_1.0 (100 kDa)_ vs. HA-ADH_2.33_HA-Ox_1.0 (100 kDa)_ | ns | 0.9915 |
| HA-ADH_3.0_HA-Ox_1.0 (100 kDa)_ vs. HA-ADH_1.67_HA-Ox_3.0 (100 kDa)_ | ns | 0.9682 |
| HA-ADH_3.0_HA-Ox_1.0 (100 kDa)_ vs. HA-ADH_2.0_HA-Ox_2.0 (100 kDa)_ | ns | 0.9996 |
| HA-ADH_1.0_HA-Ox_3.0 (100 kDa)_ vs. HA-ADH_3.0_HA-Ox_3.0 (100 kDa)_ | ns | >0.9999 |
| HA-ADH_1.0_HA-Ox_3.0 (100 kDa)_ vs. HA-ADH_1.0_HA-Ox_1.67 (100 kDa)_ | ns | >0.9999 |
| HA-ADH_1.0_HA-Ox_3.0 (100 kDa)_ vs. HA-ADH_3.0_HA-Ox_2.33 (100 kDa)_ | ns | >0.9999 |
| HA-ADH_1.0_HA-Ox_3.0 (100 kDa)_ vs. HA-ADH_2.33_HA-Ox_1.0 (100 kDa)_ | ns | >0.9999 |
| HA-ADH_1.0_HA-Ox_3.0 (100 kDa)_ vs. HA-ADH_1.67_HA-Ox_3.0 (100 kDa)_ | ns | >0.9999 |
| HA-ADH_1.0_HA-Ox_3.0 (100 kDa)_ vs. HA-ADH_2.0_HA-Ox_2.0 (100 kDa)_ | ns | 0.8547 |
| HA-ADH_3.0_HA-Ox_3.0 (100 kDa)_ vs. HA-ADH_1.0_HA-Ox_1.67 (100 kDa)_ | ns | >0.9999 |
| HA-ADH_3.0_HA-Ox_3.0 (100 kDa)_ vs. HA-ADH_3.0_HA-Ox_2.33 (100 kDa)_ | ns | 0.9655 |
| HA-ADH_3.0_HA-Ox_3.0 (100 kDa)_ vs. HA-ADH_2.33_HA-Ox_1.0 (100 kDa)_ | ns | >0.9999 |
| HA-ADH_3.0_HA-Ox_3.0 (100 kDa)_ vs. HA-ADH_1.67_HA-Ox_3.0 (100 kDa)_ | ns | >0.9999 |
| HA-ADH_3.0_HA-Ox_3.0 (100 kDa)_ vs. HA-ADH_2.0_HA-Ox_2.0 (100 kDa)_ | ns | 0.1859 |
| HA-ADH_1.0_HA-Ox_1.67 (100 kDa)_ vs. HA-ADH_3.0_HA-Ox_2.33 (100 kDa)_ | ns | 0.6977 |
| HA-ADH_1.0_HA-Ox_1.67 (100 kDa)_ vs. HA-ADH_2.33_HA-Ox_1.0 (100 kDa)_ | ns | >0.9999 |
| HA-ADH_1.0_HA-Ox_1.67 (100 kDa)_ vs. HA-ADH_1.67_HA-Ox_3.0 (100 kDa)_ | ns | >0.9999 |
| HA-ADH_1.0_HA-Ox_1.67 (100 kDa)_ vs. HA-ADH_2.0_HA-Ox_2.0 (100 kDa)_ | ns | 0.1135 |
| HA-ADH_3.0_HA-Ox_2.33 (100 kDa)_ vs. HA-ADH_2.33_HA-Ox_1.0 (100 kDa)_ | ns | 0.7478 |
| HA-ADH_3.0_HA-Ox_2.33 (100 kDa)_ vs. HA-ADH_1.67_HA-Ox_3.0 (100 kDa)_ | ns | 0.5603 |
| HA-ADH_3.0_HA-Ox_2.33 (100 kDa)_ vs. HA-ADH_2.0_HA-Ox_2.0 (100 kDa)_ | ns | 0.2718 |
| HA-ADH_2.33_HA-Ox_1.0 (100 kDa)_ vs. HA-ADH_1.67_HA-Ox_3.0 (100 kDa)_ | ns | >0.9999 |
| HA-ADH_2.33_HA-Ox_1.0 (100 kDa)_ vs. HA-ADH_2.0_HA-Ox_2.0 (100 kDa)_ | ns | 0.0538 |
| HA-ADH_1.67_HA-Ox_3.0 (100 kDa)_ vs. HA-ADH_2.0_HA-Ox_2.0 (100 kDa)_ | * | 0.044 |

**Table 3. Summary of significance for 40 kDa HA compressive modulus and 100 kDa HA compressive modulus.** A summary of significance for the compressive modulus for both 40 kDa and 100 kDa is listed, a one-way ANOVA post-hoc Dunnett’s T3 multiple comparisons was performed. n = 3; * p < 0.05, ** < p 0.01, *** p < 0.001, **** p < 0.0001.

*Mass Change*

| **Mass Change Significance** | | |
| --- | --- | --- |
| **Tukey's multiple comparisons test** | Summary | Adjusted P Value |
| **Day 1** | | |
| HA-ADH_3.0_HA-Ox_3.0 (40 kDa)_ vs. HA-ADH_3.0_HA-Ox_1.0 (40 kDa)_ | ns | 0.484 |
| HA-ADH_3.0_HA-Ox_3.0 (40 kDa)_ vs. HA-ADH_2.0_HA-Ox_2.0 (40 kDa)_ | ns | 0.5557 |
| HA-ADH_3.0_HA-Ox_3.0 (40 kDa)_ vs. HA-ADH_1.67_HA-Ox_1.0 (40 kDa)_ | ns | 0.6338 |
| HA-ADH_3.0_HA-Ox_3.0 (40 kDa)_ vs. HA-ADH_1.0_HA-Ox_3.0 (40 kDa)_ | ns | 0.9969 |
| HA-ADH_3.0_HA-Ox_3.0 (40 kDa)_ vs. HA-ADH_1.0_HA-Ox_1.0 (40 kDa)_ | ns | 0.3629 |
| HA-ADH_3.0_HA-Ox_3.0 (40 kDa)_ vs. HA-ADH_3.0_HA-Ox_3.0 (100 kDa)_ | ns | 0.1086 |
| HA-ADH_3.0_HA-Ox_3.0 (40 kDa)_ vs. HA-ADH_3.0_HA-Ox_2.33 (100 kDa)_ | ns | 0.7109 |
| HA-ADH_3.0_HA-Ox_3.0 (40 kDa)_ vs. HA-ADH_3.0_HA-Ox_1.0 (100 kDa)_ | ns | >0.9999 |
| HA-ADH_3.0_HA-Ox_3.0 (40 kDa)_ vs. HA-ADH_2.33_HA-Ox_1.0 (100 kDa)_ | ns | 0.9294 |
| HA-ADH_3.0_HA-Ox_3.0 (40 kDa)_ vs. HA-ADH_2.0_HA-Ox_2.0 (100 kDa)_ | ns | 0.7845 |
| HA-ADH_3.0_HA-Ox_3.0 (40 kDa)_ vs. HA-ADH_1.67_HA-Ox_3.0 (100 kDa)_ | ns | >0.9999 |
| HA-ADH_3.0_HA-Ox_3.0 (40 kDa)_ vs. HA-ADH_1.0_HA-Ox_3.0 (100 kDa)_ | ns | 0.683 |
| HA-ADH_3.0_HA-Ox_3.0 (40 kDa)_ vs. HA-ADH_1.0_HA-Ox_1.67 (100 kDa)_ | ns | >0.9999 |
| HA-ADH_3.0_HA-Ox_3.0 (40 kDa)_ vs. HA-ADH_1.0_HA-Ox_1.0 (100 kDa)_ | ns | 0.3514 |
| HA-ADH_3.0_HA-Ox_1.0 (40 kDa)_ vs. HA-ADH_2.0_HA-Ox_2.0 (40 kDa)_ | ns | 0.7532 |
| HA-ADH_3.0_HA-Ox_1.0 (40 kDa)_ vs. HA-ADH_1.67_HA-Ox_1.0 (40 kDa)_ | ns | 0.7932 |
| HA-ADH_3.0_HA-Ox_1.0 (40 kDa)_ vs. HA-ADH_1.0_HA-Ox_3.0 (40 kDa)_ | ns | >0.9999 |
| HA-ADH_3.0_HA-Ox_1.0 (40 kDa)_ vs. HA-ADH_1.0_HA-Ox_1.0 (40 kDa)_ | ns | >0.9999 |
| HA-ADH_3.0_HA-Ox_1.0 (40 kDa)_ vs. HA-ADH_3.0_HA-Ox_3.0 (100 kDa)_ | ns | 0.1164 |
| HA-ADH_3.0_HA-Ox_1.0 (40 kDa)_ vs. HA-ADH_3.0_HA-Ox_2.33 (100 kDa)_ | ns | 0.2618 |
| HA-ADH_3.0_HA-Ox_1.0 (40 kDa)_ vs. HA-ADH_3.0_HA-Ox_1.0 (100 kDa)_ | ns | 0.5469 |
| HA-ADH_3.0_HA-Ox_1.0 (40 kDa)_ vs. HA-ADH_2.33_HA-Ox_1.0 (100 kDa)_ | ns | 0.3309 |
| HA-ADH_3.0_HA-Ox_1.0 (40 kDa)_ vs. HA-ADH_2.0_HA-Ox_2.0 (100 kDa)_ | ns | 0.2637 |
| HA-ADH_3.0_HA-Ox_1.0 (40 kDa)_ vs. HA-ADH_1.67_HA-Ox_3.0 (100 kDa)_ | ns | 0.855 |
| HA-ADH_3.0_HA-Ox_1.0 (40 kDa)_ vs. HA-ADH_1.0_HA-Ox_3.0 (100 kDa)_ | ns | >0.9999 |
| HA-ADH_3.0_HA-Ox_1.0 (40 kDa)_ vs. HA-ADH_1.0_HA-Ox_1.67 (100 kDa)_ | ns | 0.8929 |
| HA-ADH_3.0_HA-Ox_1.0 (40 kDa)_ vs. HA-ADH_1.0_HA-Ox_1.0 (100 kDa)_ | ns | 0.2722 |
| HA-ADH_2.0_HA-Ox_2.0 (40 kDa)_ vs. HA-ADH_1.67_HA-Ox_1.0 (40 kDa)_ | ns | >0.9999 |
| HA-ADH_2.0_HA-Ox_2.0 (40 kDa)_ vs. HA-ADH_1.0_HA-Ox_3.0 (40 kDa)_ | ns | 0.9999 |
| HA-ADH_2.0_HA-Ox_2.0 (40 kDa)_ vs. HA-ADH_1.0_HA-Ox_1.0 (40 kDa)_ | ns | 0.7066 |
| HA-ADH_2.0_HA-Ox_2.0 (40 kDa)_ vs. HA-ADH_3.0_HA-Ox_3.0 (100 kDa)_ | ns | 0.0524 |
| HA-ADH_2.0_HA-Ox_2.0 (40 kDa)_ vs. HA-ADH_3.0_HA-Ox_2.33 (100 kDa)_ | ns | 0.302 |
| HA-ADH_2.0_HA-Ox_2.0 (40 kDa)_ vs. HA-ADH_3.0_HA-Ox_1.0 (100 kDa)_ | ns | 0.9163 |
| HA-ADH_2.0_HA-Ox_2.0 (40 kDa)_ vs. HA-ADH_2.33_HA-Ox_1.0 (100 kDa)_ | ns | 0.424 |
| HA-ADH_2.0_HA-Ox_2.0 (40 kDa)_ vs. HA-ADH_2.0_HA-Ox_2.0 (100 kDa)_ | ns | 0.489 |
| HA-ADH_2.0_HA-Ox_2.0 (40 kDa)_ vs. HA-ADH_1.67_HA-Ox_3.0 (100 kDa)_ | ns | >0.9999 |
| HA-ADH_2.0_HA-Ox_2.0 (40 kDa)_ vs. HA-ADH_1.0_HA-Ox_3.0 (100 kDa)_ | ns | 0.9259 |
| HA-ADH_2.0_HA-Ox_2.0 (40 kDa)_ vs. HA-ADH_1.0_HA-Ox_1.67 (100 kDa)_ | ns | >0.9999 |
| HA-ADH_2.0_HA-Ox_2.0 (40 kDa)_ vs. HA-ADH_1.0_HA-Ox_1.0 (100 kDa)_ | ns | 0.068 |
| HA-ADH_1.67_HA-Ox_1.0 (40 kDa)_ vs. HA-ADH_1.0_HA-Ox_3.0 (40 kDa)_ | ns | >0.9999 |
| HA-ADH_1.67_HA-Ox_1.0 (40 kDa)_ vs. HA-ADH_1.0_HA-Ox_1.0 (40 kDa)_ | ns | 0.773 |
| HA-ADH_1.67_HA-Ox_1.0 (40 kDa)_ vs. HA-ADH_3.0_HA-Ox_3.0 (100 kDa)_ | * | 0.0444 |
| HA-ADH_1.67_HA-Ox_1.0 (40 kDa)_ vs. HA-ADH_3.0_HA-Ox_2.33 (100 kDa)_ | ns | 0.2954 |
| HA-ADH_1.67_HA-Ox_1.0 (40 kDa)_ vs. HA-ADH_3.0_HA-Ox_1.0 (100 kDa)_ | ns | 0.9206 |
| HA-ADH_1.67_HA-Ox_1.0 (40 kDa)_ vs. HA-ADH_2.33_HA-Ox_1.0 (100 kDa)_ | ns | 0.4266 |
| HA-ADH_1.67_HA-Ox_1.0 (40 kDa)_ vs. HA-ADH_2.0_HA-Ox_2.0 (100 kDa)_ | ns | 0.4784 |
| HA-ADH_1.67_HA-Ox_1.0 (40 kDa)_ vs. HA-ADH_1.67_HA-Ox_3.0 (100 kDa)_ | ns | >0.9999 |
| HA-ADH_1.67_HA-Ox_1.0 (40 kDa)_ vs. HA-ADH_1.0_HA-Ox_3.0 (100 kDa)_ | ns | 0.9473 |
| HA-ADH_1.67_HA-Ox_1.0 (40 kDa)_ vs. HA-ADH_1.0_HA-Ox_1.67 (100 kDa)_ | ns | >0.9999 |
| HA-ADH_1.67_HA-Ox_1.0 (40 kDa)_ vs. HA-ADH_1.0_HA-Ox_1.0 (100 kDa)_ | ns | 0.1043 |
| HA-ADH_1.0_HA-Ox_3.0 (40 kDa)_ vs. HA-ADH_1.0_HA-Ox_1.0 (40 kDa)_ | ns | >0.9999 |
| HA-ADH_1.0_HA-Ox_3.0 (40 kDa)_ vs. HA-ADH_3.0_HA-Ox_3.0 (100 kDa)_ | ns | 0.8693 |
| HA-ADH_1.0_HA-Ox_3.0 (40 kDa)_ vs. HA-ADH_3.0_HA-Ox_2.33 (100 kDa)_ | ns | 0.9709 |
| HA-ADH_1.0_HA-Ox_3.0 (40 kDa)_ vs. HA-ADH_3.0_HA-Ox_1.0 (100 kDa)_ | ns | 0.998 |
| HA-ADH_1.0_HA-Ox_3.0 (40 kDa)_ vs. HA-ADH_2.33_HA-Ox_1.0 (100 kDa)_ | ns | 0.9846 |
| HA-ADH_1.0_HA-Ox_3.0 (40 kDa)_ vs. HA-ADH_2.0_HA-Ox_2.0 (100 kDa)_ | ns | 0.9557 |
| HA-ADH_1.0_HA-Ox_3.0 (40 kDa)_ vs. HA-ADH_1.67_HA-Ox_3.0 (100 kDa)_ | ns | 0.9997 |
| HA-ADH_1.0_HA-Ox_3.0 (40 kDa)_ vs. HA-ADH_1.0_HA-Ox_3.0 (100 kDa)_ | ns | >0.9999 |
| HA-ADH_1.0_HA-Ox_3.0 (40 kDa)_ vs. HA-ADH_1.0_HA-Ox_1.67 (100 kDa)_ | ns | 0.9997 |
| HA-ADH_1.0_HA-Ox_3.0 (40 kDa)_ vs. HA-ADH_1.0_HA-Ox_1.0 (100 kDa)_ | ns | 0.9713 |
| HA-ADH_1.0_HA-Ox_1.0 (40 kDa)_ vs. HA-ADH_3.0_HA-Ox_3.0 (100 kDa)_ | * | 0.0491 |
| HA-ADH_1.0_HA-Ox_1.0 (40 kDa)_ vs. HA-ADH_3.0_HA-Ox_2.33 (100 kDa)_ | ns | 0.1617 |
| HA-ADH_1.0_HA-Ox_1.0 (40 kDa)_ vs. HA-ADH_3.0_HA-Ox_1.0 (100 kDa)_ | ns | 0.4526 |
| HA-ADH_1.0_HA-Ox_1.0 (40 kDa)_ vs. HA-ADH_2.33_HA-Ox_1.0 (100 kDa)_ | ns | 0.2208 |
| HA-ADH_1.0_HA-Ox_1.0 (40 kDa)_ vs. HA-ADH_2.0_HA-Ox_2.0 (100 kDa)_ | ns | 0.2375 |
| HA-ADH_1.0_HA-Ox_1.0 (40 kDa)_ vs. HA-ADH_1.67_HA-Ox_3.0 (100 kDa)_ | ns | 0.8852 |
| HA-ADH_1.0_HA-Ox_1.0 (40 kDa)_ vs. HA-ADH_1.0_HA-Ox_3.0 (100 kDa)_ | ns | >0.9999 |
| HA-ADH_1.0_HA-Ox_1.0 (40 kDa)_ vs. HA-ADH_1.0_HA-Ox_1.67 (100 kDa)_ | ns | 0.9225 |
| HA-ADH_1.0_HA-Ox_1.0 (40 kDa)_ vs. HA-ADH_1.0_HA-Ox_1.0 (100 kDa)_ | ns | 0.1607 |
| HA-ADH_3.0_HA-Ox_3.0 (100 kDa)_ vs. HA-ADH_3.0_HA-Ox_2.33 (100 kDa)_ | ns | 0.6167 |
| HA-ADH_3.0_HA-Ox_3.0 (100 kDa)_ vs. HA-ADH_3.0_HA-Ox_1.0 (100 kDa)_ | ns | 0.1228 |
| HA-ADH_3.0_HA-Ox_3.0 (100 kDa)_ vs. HA-ADH_2.33_HA-Ox_1.0 (100 kDa)_ | ns | 0.3806 |
| HA-ADH_3.0_HA-Ox_3.0 (100 kDa)_ vs. HA-ADH_2.0_HA-Ox_2.0 (100 kDa)_ | ns | 0.9736 |
| HA-ADH_3.0_HA-Ox_3.0 (100 kDa)_ vs. HA-ADH_1.67_HA-Ox_3.0 (100 kDa)_ | ns | 0.3941 |
| HA-ADH_3.0_HA-Ox_3.0 (100 kDa)_ vs. HA-ADH_1.0_HA-Ox_3.0 (100 kDa)_ | ns | 0.1959 |
| HA-ADH_3.0_HA-Ox_3.0 (100 kDa)_ vs. HA-ADH_1.0_HA-Ox_1.67 (100 kDa)_ | ns | 0.457 |
| HA-ADH_3.0_HA-Ox_3.0 (100 kDa)_ vs. HA-ADH_1.0_HA-Ox_1.0 (100 kDa)_ | ns | 0.3744 |
| HA-ADH_3.0_HA-Ox_2.33 (100 kDa)_ vs. HA-ADH_3.0_HA-Ox_1.0 (100 kDa)_ | ns | 0.7645 |
| HA-ADH_3.0_HA-Ox_2.33 (100 kDa)_ vs. HA-ADH_2.33_HA-Ox_1.0 (100 kDa)_ | ns | >0.9999 |
| HA-ADH_3.0_HA-Ox_2.33 (100 kDa)_ vs. HA-ADH_2.0_HA-Ox_2.0 (100 kDa)_ | ns | >0.9999 |
| HA-ADH_3.0_HA-Ox_2.33 (100 kDa)_ vs. HA-ADH_1.67_HA-Ox_3.0 (100 kDa)_ | ns | 0.8888 |
| HA-ADH_3.0_HA-Ox_2.33 (100 kDa)_ vs. HA-ADH_1.0_HA-Ox_3.0 (100 kDa)_ | ns | 0.4145 |
| HA-ADH_3.0_HA-Ox_2.33 (100 kDa)_ vs. HA-ADH_1.0_HA-Ox_1.67 (100 kDa)_ | ns | 0.9128 |
| HA-ADH_3.0_HA-Ox_2.33 (100 kDa)_ vs. HA-ADH_1.0_HA-Ox_1.0 (100 kDa)_ | ns | >0.9999 |
| HA-ADH_3.0_HA-Ox_1.0 (100 kDa)_ vs. HA-ADH_2.33_HA-Ox_1.0 (100 kDa)_ | ns | 0.9491 |
| HA-ADH_3.0_HA-Ox_1.0 (100 kDa)_ vs. HA-ADH_2.0_HA-Ox_2.0 (100 kDa)_ | ns | 0.7916 |
| HA-ADH_3.0_HA-Ox_1.0 (100 kDa)_ vs. HA-ADH_1.67_HA-Ox_3.0 (100 kDa)_ | ns | >0.9999 |
| HA-ADH_3.0_HA-Ox_1.0 (100 kDa)_ vs. HA-ADH_1.0_HA-Ox_3.0 (100 kDa)_ | ns | 0.7551 |
| HA-ADH_3.0_HA-Ox_1.0 (100 kDa)_ vs. HA-ADH_1.0_HA-Ox_1.67 (100 kDa)_ | ns | >0.9999 |
| HA-ADH_3.0_HA-Ox_1.0 (100 kDa)_ vs. HA-ADH_1.0_HA-Ox_1.0 (100 kDa)_ | ns | 0.5763 |
| HA-ADH_2.33_HA-Ox_1.0 (100 kDa)_ vs. HA-ADH_2.0_HA-Ox_2.0 (100 kDa)_ | ns | 0.9963 |
| HA-ADH_2.33_HA-Ox_1.0 (100 kDa)_ vs. HA-ADH_1.67_HA-Ox_3.0 (100 kDa)_ | ns | 0.9713 |
| HA-ADH_2.33_HA-Ox_1.0 (100 kDa)_ vs. HA-ADH_1.0_HA-Ox_3.0 (100 kDa)_ | ns | 0.5067 |
| HA-ADH_2.33_HA-Ox_1.0 (100 kDa)_ vs. HA-ADH_1.0_HA-Ox_1.67 (100 kDa)_ | ns | 0.9785 |
| HA-ADH_2.33_HA-Ox_1.0 (100 kDa)_ vs. HA-ADH_1.0_HA-Ox_1.0 (100 kDa)_ | ns | 0.9998 |
| HA-ADH_2.0_HA-Ox_2.0 (100 kDa)_ vs. HA-ADH_1.67_HA-Ox_3.0 (100 kDa)_ | ns | 0.8588 |
| HA-ADH_2.0_HA-Ox_2.0 (100 kDa)_ vs. HA-ADH_1.0_HA-Ox_3.0 (100 kDa)_ | ns | 0.3959 |
| HA-ADH_2.0_HA-Ox_2.0 (100 kDa)_ vs. HA-ADH_1.0_HA-Ox_1.67 (100 kDa)_ | ns | 0.8818 |
| HA-ADH_2.0_HA-Ox_2.0 (100 kDa)_ vs. HA-ADH_1.0_HA-Ox_1.0 (100 kDa)_ | ns | 0.9999 |
| HA-ADH_1.67_HA-Ox_3.0 (100 kDa)_ vs. HA-ADH_1.0_HA-Ox_3.0 (100 kDa)_ | ns | 0.9574 |
| HA-ADH_1.67_HA-Ox_3.0 (100 kDa)_ vs. HA-ADH_1.0_HA-Ox_1.67 (100 kDa)_ | ns | >0.9999 |
| HA-ADH_1.67_HA-Ox_3.0 (100 kDa)_ vs. HA-ADH_1.0_HA-Ox_1.0 (100 kDa)_ | ns | 0.8511 |
| HA-ADH_1.0_HA-Ox_3.0 (100 kDa)_ vs. HA-ADH_1.0_HA-Ox_1.67 (100 kDa)_ | ns | 0.9718 |
| HA-ADH_1.0_HA-Ox_3.0 (100 kDa)_ vs. HA-ADH_1.0_HA-Ox_1.0 (100 kDa)_ | ns | 0.4109 |
| HA-ADH_1.0_HA-Ox_1.67 (100 kDa)_ vs. HA-ADH_1.0_HA-Ox_1.0 (100 kDa)_ | ns | 0.886 |
| **Day 7** | | |
| HA-ADH_3.0_HA-Ox_3.0 (40 kDa)_ vs. HA-ADH_3.0_HA-Ox_1.0 (40 kDa)_ | ns | >0.9999 |
| HA-ADH_3.0_HA-Ox_3.0 (40 kDa)_ vs. HA-ADH_2.0_HA-Ox_2.0 (40 kDa)_ | ns | >0.9999 |
| HA-ADH_3.0_HA-Ox_3.0 (40 kDa)_ vs. HA-ADH_1.67_HA-Ox_1.0 (40 kDa)_ | ns | >0.9999 |
| HA-ADH_3.0_HA-Ox_3.0 (40 kDa)_ vs. HA-ADH_1.0_HA-Ox_3.0 (40 kDa)_ | ns | 0.988 |
| HA-ADH_3.0_HA-Ox_3.0 (40 kDa)_ vs. HA-ADH_1.0_HA-Ox_1.0 (40 kDa)_ | ns | 0.1866 |
| HA-ADH_3.0_HA-Ox_3.0 (40 kDa)_ vs. HA-ADH_3.0_HA-Ox_3.0 (100 kDa)_ | ns | 0.6369 |
| HA-ADH_3.0_HA-Ox_3.0 (40 kDa)_ vs. HA-ADH_3.0_HA-Ox_2.33 (100 kDa)_ | ns | 0.9882 |
| HA-ADH_3.0_HA-Ox_3.0 (40 kDa)_ vs. HA-ADH_3.0_HA-Ox_1.0 (100 kDa)_ | ns | 0.925 |
| HA-ADH_3.0_HA-Ox_3.0 (40 kDa)_ vs. HA-ADH_2.33_HA-Ox_1.0 (100 kDa)_ | ns | 0.8254 |
| HA-ADH_3.0_HA-Ox_3.0 (40 kDa)_ vs. HA-ADH_2.0_HA-Ox_2.0 (100 kDa)_ | ns | 0.6777 |
| HA-ADH_3.0_HA-Ox_3.0 (40 kDa)_ vs. HA-ADH_1.67_HA-Ox_3.0 (100 kDa)_ | ns | 0.7135 |
| HA-ADH_3.0_HA-Ox_3.0 (40 kDa)_ vs. HA-ADH_1.0_HA-Ox_3.0 (100 kDa)_ | ns | 0.6983 |
| HA-ADH_3.0_HA-Ox_3.0 (40 kDa)_ vs. HA-ADH_1.0_HA-Ox_1.67 (100 kDa)_ | ns | >0.9999 |
| HA-ADH_3.0_HA-Ox_3.0 (40 kDa)_ vs. HA-ADH_1.0_HA-Ox_1.0 (100 kDa)_ | ns | 0.9999 |
| HA-ADH_3.0_HA-Ox_1.0 (40 kDa)_ vs. HA-ADH_2.0_HA-Ox_2.0 (40 kDa)_ | ns | >0.9999 |
| HA-ADH_3.0_HA-Ox_1.0 (40 kDa)_ vs. HA-ADH_1.67_HA-Ox_1.0 (40 kDa)_ | ns | >0.9999 |
| HA-ADH_3.0_HA-Ox_1.0 (40 kDa)_ vs. HA-ADH_1.0_HA-Ox_3.0 (40 kDa)_ | ns | >0.9999 |
| HA-ADH_3.0_HA-Ox_1.0 (40 kDa)_ vs. HA-ADH_1.0_HA-Ox_1.0 (40 kDa)_ | ns | 0.9675 |
| HA-ADH_3.0_HA-Ox_1.0 (40 kDa)_ vs. HA-ADH_3.0_HA-Ox_3.0 (100 kDa)_ | ns | 0.9998 |
| HA-ADH_3.0_HA-Ox_1.0 (40 kDa)_ vs. HA-ADH_3.0_HA-Ox_2.33 (100 kDa)_ | ns | >0.9999 |
| HA-ADH_3.0_HA-Ox_1.0 (40 kDa)_ vs. HA-ADH_3.0_HA-Ox_1.0 (100 kDa)_ | ns | >0.9999 |
| HA-ADH_3.0_HA-Ox_1.0 (40 kDa)_ vs. HA-ADH_2.33_HA-Ox_1.0 (100 kDa)_ | ns | >0.9999 |
| HA-ADH_3.0_HA-Ox_1.0 (40 kDa)_ vs. HA-ADH_2.0_HA-Ox_2.0 (100 kDa)_ | ns | >0.9999 |
| HA-ADH_3.0_HA-Ox_1.0 (40 kDa)_ vs. HA-ADH_1.67_HA-Ox_3.0 (100 kDa)_ | ns | 0.9995 |
| HA-ADH_3.0_HA-Ox_1.0 (40 kDa)_ vs. HA-ADH_1.0_HA-Ox_3.0 (100 kDa)_ | ns | 0.9984 |
| HA-ADH_3.0_HA-Ox_1.0 (40 kDa)_ vs. HA-ADH_1.0_HA-Ox_1.67 (100 kDa)_ | ns | >0.9999 |
| HA-ADH_3.0_HA-Ox_1.0 (40 kDa)_ vs. HA-ADH_1.0_HA-Ox_1.0 (100 kDa)_ | ns | >0.9999 |
| HA-ADH_2.0_HA-Ox_2.0 (40 kDa)_ vs. HA-ADH_1.67_HA-Ox_1.0 (40 kDa)_ | ns | >0.9999 |
| HA-ADH_2.0_HA-Ox_2.0 (40 kDa)_ vs. HA-ADH_1.0_HA-Ox_3.0 (40 kDa)_ | ns | >0.9999 |
| HA-ADH_2.0_HA-Ox_2.0 (40 kDa)_ vs. HA-ADH_1.0_HA-Ox_1.0 (40 kDa)_ | ns | 0.5016 |
| HA-ADH_2.0_HA-Ox_2.0 (40 kDa)_ vs. HA-ADH_3.0_HA-Ox_3.0 (100 kDa)_ | ns | 0.9642 |
| HA-ADH_2.0_HA-Ox_2.0 (40 kDa)_ vs. HA-ADH_3.0_HA-Ox_2.33 (100 kDa)_ | ns | >0.9999 |
| HA-ADH_2.0_HA-Ox_2.0 (40 kDa)_ vs. HA-ADH_3.0_HA-Ox_1.0 (100 kDa)_ | ns | >0.9999 |
| HA-ADH_2.0_HA-Ox_2.0 (40 kDa)_ vs. HA-ADH_2.33_HA-Ox_1.0 (100 kDa)_ | ns | 0.9982 |
| HA-ADH_2.0_HA-Ox_2.0 (40 kDa)_ vs. HA-ADH_2.0_HA-Ox_2.0 (100 kDa)_ | ns | 0.9838 |
| HA-ADH_2.0_HA-Ox_2.0 (40 kDa)_ vs. HA-ADH_1.67_HA-Ox_3.0 (100 kDa)_ | ns | 0.9643 |
| HA-ADH_2.0_HA-Ox_2.0 (40 kDa)_ vs. HA-ADH_1.0_HA-Ox_3.0 (100 kDa)_ | ns | 0.9371 |
| HA-ADH_2.0_HA-Ox_2.0 (40 kDa)_ vs. HA-ADH_1.0_HA-Ox_1.67 (100 kDa)_ | ns | >0.9999 |
| HA-ADH_2.0_HA-Ox_2.0 (40 kDa)_ vs. HA-ADH_1.0_HA-Ox_1.0 (100 kDa)_ | ns | >0.9999 |
| HA-ADH_1.67_HA-Ox_1.0 (40 kDa)_ vs. HA-ADH_1.0_HA-Ox_3.0 (40 kDa)_ | ns | 0.9815 |
| HA-ADH_1.67_HA-Ox_1.0 (40 kDa)_ vs. HA-ADH_1.0_HA-Ox_1.0 (40 kDa)_ | ns | 0.2282 |
| HA-ADH_1.67_HA-Ox_1.0 (40 kDa)_ vs. HA-ADH_3.0_HA-Ox_3.0 (100 kDa)_ | ns | 0.6514 |
| HA-ADH_1.67_HA-Ox_1.0 (40 kDa)_ vs. HA-ADH_3.0_HA-Ox_2.33 (100 kDa)_ | ns | 0.9773 |
| HA-ADH_1.67_HA-Ox_1.0 (40 kDa)_ vs. HA-ADH_3.0_HA-Ox_1.0 (100 kDa)_ | ns | 0.9105 |
| HA-ADH_1.67_HA-Ox_1.0 (40 kDa)_ vs. HA-ADH_2.33_HA-Ox_1.0 (100 kDa)_ | ns | 0.8193 |
| HA-ADH_1.67_HA-Ox_1.0 (40 kDa)_ vs. HA-ADH_2.0_HA-Ox_2.0 (100 kDa)_ | ns | 0.6974 |
| HA-ADH_1.67_HA-Ox_1.0 (40 kDa)_ vs. HA-ADH_1.67_HA-Ox_3.0 (100 kDa)_ | ns | 0.7032 |
| HA-ADH_1.67_HA-Ox_1.0 (40 kDa)_ vs. HA-ADH_1.0_HA-Ox_3.0 (100 kDa)_ | ns | 0.6834 |
| HA-ADH_1.67_HA-Ox_1.0 (40 kDa)_ vs. HA-ADH_1.0_HA-Ox_1.67 (100 kDa)_ | ns | >0.9999 |
| HA-ADH_1.67_HA-Ox_1.0 (40 kDa)_ vs. HA-ADH_1.0_HA-Ox_1.0 (100 kDa)_ | ns | 0.9988 |
| HA-ADH_1.0_HA-Ox_3.0 (40 kDa)_ vs. HA-ADH_1.0_HA-Ox_1.0 (40 kDa)_ | ns | 0.9026 |
| HA-ADH_1.0_HA-Ox_3.0 (40 kDa)_ vs. HA-ADH_3.0_HA-Ox_3.0 (100 kDa)_ | ns | >0.9999 |
| HA-ADH_1.0_HA-Ox_3.0 (40 kDa)_ vs. HA-ADH_3.0_HA-Ox_2.33 (100 kDa)_ | ns | >0.9999 |
| HA-ADH_1.0_HA-Ox_3.0 (40 kDa)_ vs. HA-ADH_3.0_HA-Ox_1.0 (100 kDa)_ | ns | >0.9999 |
| HA-ADH_1.0_HA-Ox_3.0 (40 kDa)_ vs. HA-ADH_2.33_HA-Ox_1.0 (100 kDa)_ | ns | >0.9999 |
| HA-ADH_1.0_HA-Ox_3.0 (40 kDa)_ vs. HA-ADH_2.0_HA-Ox_2.0 (100 kDa)_ | ns | >0.9999 |
| HA-ADH_1.0_HA-Ox_3.0 (40 kDa)_ vs. HA-ADH_1.67_HA-Ox_3.0 (100 kDa)_ | ns | >0.9999 |
| HA-ADH_1.0_HA-Ox_3.0 (40 kDa)_ vs. HA-ADH_1.0_HA-Ox_3.0 (100 kDa)_ | ns | 0.9999 |
| HA-ADH_1.0_HA-Ox_3.0 (40 kDa)_ vs. HA-ADH_1.0_HA-Ox_1.67 (100 kDa)_ | ns | 0.9997 |
| HA-ADH_1.0_HA-Ox_3.0 (40 kDa)_ vs. HA-ADH_1.0_HA-Ox_1.0 (100 kDa)_ | ns | 0.9974 |
| HA-ADH_1.0_HA-Ox_1.0 (40 kDa)_ vs. HA-ADH_3.0_HA-Ox_3.0 (100 kDa)_ | ns | 0.5108 |
| HA-ADH_1.0_HA-Ox_1.0 (40 kDa)_ vs. HA-ADH_3.0_HA-Ox_2.33 (100 kDa)_ | ns | 0.1558 |
| HA-ADH_1.0_HA-Ox_1.0 (40 kDa)_ vs. HA-ADH_3.0_HA-Ox_1.0 (100 kDa)_ | ns | 0.1621 |
| HA-ADH_1.0_HA-Ox_1.0 (40 kDa)_ vs. HA-ADH_2.33_HA-Ox_1.0 (100 kDa)_ | ns | 0.2364 |
| HA-ADH_1.0_HA-Ox_1.0 (40 kDa)_ vs. HA-ADH_2.0_HA-Ox_2.0 (100 kDa)_ | ns | 0.2819 |
| HA-ADH_1.0_HA-Ox_1.0 (40 kDa)_ vs. HA-ADH_1.67_HA-Ox_3.0 (100 kDa)_ | ns | 0.9047 |
| HA-ADH_1.0_HA-Ox_1.0 (40 kDa)_ vs. HA-ADH_1.0_HA-Ox_3.0 (100 kDa)_ | ns | 0.9944 |
| HA-ADH_1.0_HA-Ox_1.0 (40 kDa)_ vs. HA-ADH_1.0_HA-Ox_1.67 (100 kDa)_ | ns | 0.3217 |
| HA-ADH_1.0_HA-Ox_1.0 (40 kDa)_ vs. HA-ADH_1.0_HA-Ox_1.0 (100 kDa)_ | ns | 0.0907 |
| HA-ADH_3.0_HA-Ox_3.0 (100 kDa)_ vs. HA-ADH_3.0_HA-Ox_2.33 (100 kDa)_ | ns | 0.8528 |
| HA-ADH_3.0_HA-Ox_3.0 (100 kDa)_ vs. HA-ADH_3.0_HA-Ox_1.0 (100 kDa)_ | ns | 0.8863 |
| HA-ADH_3.0_HA-Ox_3.0 (100 kDa)_ vs. HA-ADH_2.33_HA-Ox_1.0 (100 kDa)_ | ns | 0.9957 |
| HA-ADH_3.0_HA-Ox_3.0 (100 kDa)_ vs. HA-ADH_2.0_HA-Ox_2.0 (100 kDa)_ | ns | 0.9992 |
| HA-ADH_3.0_HA-Ox_3.0 (100 kDa)_ vs. HA-ADH_1.67_HA-Ox_3.0 (100 kDa)_ | ns | >0.9999 |
| HA-ADH_3.0_HA-Ox_3.0 (100 kDa)_ vs. HA-ADH_1.0_HA-Ox_3.0 (100 kDa)_ | ns | 0.9998 |
| HA-ADH_3.0_HA-Ox_3.0 (100 kDa)_ vs. HA-ADH_1.0_HA-Ox_1.67 (100 kDa)_ | ns | 0.8928 |
| HA-ADH_3.0_HA-Ox_3.0 (100 kDa)_ vs. HA-ADH_1.0_HA-Ox_1.0 (100 kDa)_ | ns | 0.5199 |
| HA-ADH_3.0_HA-Ox_2.33 (100 kDa)_ vs. HA-ADH_3.0_HA-Ox_1.0 (100 kDa)_ | ns | >0.9999 |
| HA-ADH_3.0_HA-Ox_2.33 (100 kDa)_ vs. HA-ADH_2.33_HA-Ox_1.0 (100 kDa)_ | ns | 0.9949 |
| HA-ADH_3.0_HA-Ox_2.33 (100 kDa)_ vs. HA-ADH_2.0_HA-Ox_2.0 (100 kDa)_ | ns | 0.8492 |
| HA-ADH_3.0_HA-Ox_2.33 (100 kDa)_ vs. HA-ADH_1.67_HA-Ox_3.0 (100 kDa)_ | ns | 0.9104 |
| HA-ADH_3.0_HA-Ox_2.33 (100 kDa)_ vs. HA-ADH_1.0_HA-Ox_3.0 (100 kDa)_ | ns | 0.8787 |
| HA-ADH_3.0_HA-Ox_2.33 (100 kDa)_ vs. HA-ADH_1.0_HA-Ox_1.67 (100 kDa)_ | ns | >0.9999 |
| HA-ADH_3.0_HA-Ox_2.33 (100 kDa)_ vs. HA-ADH_1.0_HA-Ox_1.0 (100 kDa)_ | ns | 0.9996 |
| HA-ADH_3.0_HA-Ox_1.0 (100 kDa)_ vs. HA-ADH_2.33_HA-Ox_1.0 (100 kDa)_ | ns | 0.9997 |
| HA-ADH_3.0_HA-Ox_1.0 (100 kDa)_ vs. HA-ADH_2.0_HA-Ox_2.0 (100 kDa)_ | ns | 0.8427 |
| HA-ADH_3.0_HA-Ox_1.0 (100 kDa)_ vs. HA-ADH_1.67_HA-Ox_3.0 (100 kDa)_ | ns | 0.9397 |
| HA-ADH_3.0_HA-Ox_1.0 (100 kDa)_ vs. HA-ADH_1.0_HA-Ox_3.0 (100 kDa)_ | ns | 0.9102 |
| HA-ADH_3.0_HA-Ox_1.0 (100 kDa)_ vs. HA-ADH_1.0_HA-Ox_1.67 (100 kDa)_ | ns | 0.9995 |
| HA-ADH_3.0_HA-Ox_1.0 (100 kDa)_ vs. HA-ADH_1.0_HA-Ox_1.0 (100 kDa)_ | ns | 0.9477 |
| HA-ADH_2.33_HA-Ox_1.0 (100 kDa)_ vs. HA-ADH_2.0_HA-Ox_2.0 (100 kDa)_ | ns | 0.9999 |
| HA-ADH_2.33_HA-Ox_1.0 (100 kDa)_ vs. HA-ADH_1.67_HA-Ox_3.0 (100 kDa)_ | ns | 0.9929 |
| HA-ADH_2.33_HA-Ox_1.0 (100 kDa)_ vs. HA-ADH_1.0_HA-Ox_3.0 (100 kDa)_ | ns | 0.9743 |
| HA-ADH_2.33_HA-Ox_1.0 (100 kDa)_ vs. HA-ADH_1.0_HA-Ox_1.67 (100 kDa)_ | ns | 0.9875 |
| HA-ADH_2.33_HA-Ox_1.0 (100 kDa)_ vs. HA-ADH_1.0_HA-Ox_1.0 (100 kDa)_ | ns | 0.7805 |
| HA-ADH_2.0_HA-Ox_2.0 (100 kDa)_ vs. HA-ADH_1.67_HA-Ox_3.0 (100 kDa)_ | ns | 0.998 |
| HA-ADH_2.0_HA-Ox_2.0 (100 kDa)_ vs. HA-ADH_1.0_HA-Ox_3.0 (100 kDa)_ | ns | 0.9868 |
| HA-ADH_2.0_HA-Ox_2.0 (100 kDa)_ vs. HA-ADH_1.0_HA-Ox_1.67 (100 kDa)_ | ns | 0.9256 |
| HA-ADH_2.0_HA-Ox_2.0 (100 kDa)_ vs. HA-ADH_1.0_HA-Ox_1.0 (100 kDa)_ | ns | 0.4203 |
| HA-ADH_1.67_HA-Ox_3.0 (100 kDa)_ vs. HA-ADH_1.0_HA-Ox_3.0 (100 kDa)_ | ns | >0.9999 |
| HA-ADH_1.67_HA-Ox_3.0 (100 kDa)_ vs. HA-ADH_1.0_HA-Ox_1.67 (100 kDa)_ | ns | 0.9126 |
| HA-ADH_1.67_HA-Ox_3.0 (100 kDa)_ vs. HA-ADH_1.0_HA-Ox_1.0 (100 kDa)_ | ns | 0.7366 |
| HA-ADH_1.0_HA-Ox_3.0 (100 kDa)_ vs. HA-ADH_1.0_HA-Ox_1.67 (100 kDa)_ | ns | 0.8762 |
| HA-ADH_1.0_HA-Ox_3.0 (100 kDa)_ vs. HA-ADH_1.0_HA-Ox_1.0 (100 kDa)_ | ns | 0.7393 |
| HA-ADH_1.0_HA-Ox_1.67 (100 kDa)_ vs. HA-ADH_1.0_HA-Ox_1.0 (100 kDa)_ | ns | >0.9999 |
| **Day 14** | | |
| HA-ADH_3.0_HA-Ox_3.0 (40 kDa)_ vs. HA-ADH_3.0_HA-Ox_1.0 (40 kDa)_ | ns | 0.4582 |
| HA-ADH_3.0_HA-Ox_3.0 (40 kDa)_ vs. HA-ADH_2.0_HA-Ox_2.0 (40 kDa)_ | ns | >0.9999 |
| HA-ADH_3.0_HA-Ox_3.0 (40 kDa)_ vs. HA-ADH_1.67_HA-Ox_1.0 (40 kDa)_ | ns | 0.8718 |
| HA-ADH_3.0_HA-Ox_3.0 (40 kDa)_ vs. HA-ADH_1.0_HA-Ox_3.0 (40 kDa)_ | ns | 0.1187 |
| HA-ADH_3.0_HA-Ox_3.0 (40 kDa)_ vs. HA-ADH_1.0_HA-Ox_1.0 (40 kDa)_ | ns | 0.1181 |
| HA-ADH_3.0_HA-Ox_3.0 (40 kDa)_ vs. HA-ADH_3.0_HA-Ox_3.0 (100 kDa)_ | ns | 0.5724 |
| HA-ADH_3.0_HA-Ox_3.0 (40 kDa)_ vs. HA-ADH_3.0_HA-Ox_2.33 (100 kDa)_ | ns | 0.9227 |
| HA-ADH_3.0_HA-Ox_3.0 (40 kDa)_ vs. HA-ADH_3.0_HA-Ox_1.0 (100 kDa)_ | ns | 0.991 |
| HA-ADH_3.0_HA-Ox_3.0 (40 kDa)_ vs. HA-ADH_2.33_HA-Ox_1.0 (100 kDa)_ | ns | 0.9455 |
| HA-ADH_3.0_HA-Ox_3.0 (40 kDa)_ vs. HA-ADH_2.0_HA-Ox_2.0 (100 kDa)_ | ns | 0.5572 |
| HA-ADH_3.0_HA-Ox_3.0 (40 kDa)_ vs. HA-ADH_1.67_HA-Ox_3.0 (100 kDa)_ | ns | 0.6568 |
| HA-ADH_3.0_HA-Ox_3.0 (40 kDa)_ vs. HA-ADH_1.0_HA-Ox_3.0 (100 kDa)_ | ns | 0.2582 |
| HA-ADH_3.0_HA-Ox_3.0 (40 kDa)_ vs. HA-ADH_1.0_HA-Ox_1.67 (100 kDa)_ | ns | 0.5764 |
| HA-ADH_3.0_HA-Ox_3.0 (40 kDa)_ vs. HA-ADH_1.0_HA-Ox_1.0 (100 kDa)_ | ns | 0.737 |
| HA-ADH_3.0_HA-Ox_1.0 (40 kDa)_ vs. HA-ADH_2.0_HA-Ox_2.0 (40 kDa)_ | ns | 0.3856 |
| HA-ADH_3.0_HA-Ox_1.0 (40 kDa)_ vs. HA-ADH_1.67_HA-Ox_1.0 (40 kDa)_ | ns | 0.7673 |
| HA-ADH_3.0_HA-Ox_1.0 (40 kDa)_ vs. HA-ADH_1.0_HA-Ox_3.0 (40 kDa)_ | ns | 0.0613 |
| HA-ADH_3.0_HA-Ox_1.0 (40 kDa)_ vs. HA-ADH_1.0_HA-Ox_1.0 (40 kDa)_ | ns | 0.0607 |
| HA-ADH_3.0_HA-Ox_1.0 (40 kDa)_ vs. HA-ADH_3.0_HA-Ox_3.0 (100 kDa)_ | ns | 0.1396 |
| HA-ADH_3.0_HA-Ox_1.0 (40 kDa)_ vs. HA-ADH_3.0_HA-Ox_2.33 (100 kDa)_ | ns | 0.2126 |
| HA-ADH_3.0_HA-Ox_1.0 (40 kDa)_ vs. HA-ADH_3.0_HA-Ox_1.0 (100 kDa)_ | ns | 0.2473 |
| HA-ADH_3.0_HA-Ox_1.0 (40 kDa)_ vs. HA-ADH_2.33_HA-Ox_1.0 (100 kDa)_ | ns | 0.1941 |
| HA-ADH_3.0_HA-Ox_1.0 (40 kDa)_ vs. HA-ADH_2.0_HA-Ox_2.0 (100 kDa)_ | ns | 0.1354 |
| HA-ADH_3.0_HA-Ox_1.0 (40 kDa)_ vs. HA-ADH_1.67_HA-Ox_3.0 (100 kDa)_ | ns | 0.1381 |
| HA-ADH_3.0_HA-Ox_1.0 (40 kDa)_ vs. HA-ADH_1.0_HA-Ox_3.0 (100 kDa)_ | ns | 0.0589 |
| HA-ADH_3.0_HA-Ox_1.0 (40 kDa)_ vs. HA-ADH_1.0_HA-Ox_1.67 (100 kDa)_ | ns | 0.0993 |
| HA-ADH_3.0_HA-Ox_1.0 (40 kDa)_ vs. HA-ADH_1.0_HA-Ox_1.0 (100 kDa)_ | ns | 0.1289 |
| HA-ADH_2.0_HA-Ox_2.0 (40 kDa)_ vs. HA-ADH_1.67_HA-Ox_1.0 (40 kDa)_ | ns | 0.641 |
| HA-ADH_2.0_HA-Ox_2.0 (40 kDa)_ vs. HA-ADH_1.0_HA-Ox_3.0 (40 kDa)_ | * | 0.031 |
| HA-ADH_2.0_HA-Ox_2.0 (40 kDa)_ vs. HA-ADH_1.0_HA-Ox_1.0 (40 kDa)_ | * | 0.0291 |
| HA-ADH_2.0_HA-Ox_2.0 (40 kDa)_ vs. HA-ADH_3.0_HA-Ox_3.0 (100 kDa)_ | ns | 0.1527 |
| HA-ADH_2.0_HA-Ox_2.0 (40 kDa)_ vs. HA-ADH_3.0_HA-Ox_2.33 (100 kDa)_ | ns | 0.4779 |
| HA-ADH_2.0_HA-Ox_2.0 (40 kDa)_ vs. HA-ADH_3.0_HA-Ox_1.0 (100 kDa)_ | ns | 0.7365 |
| HA-ADH_2.0_HA-Ox_2.0 (40 kDa)_ vs. HA-ADH_2.33_HA-Ox_1.0 (100 kDa)_ | ns | 0.5914 |
| HA-ADH_2.0_HA-Ox_2.0 (40 kDa)_ vs. HA-ADH_2.0_HA-Ox_2.0 (100 kDa)_ | ns | 0.1439 |
| HA-ADH_2.0_HA-Ox_2.0 (40 kDa)_ vs. HA-ADH_1.67_HA-Ox_3.0 (100 kDa)_ | ns | 0.208 |
| HA-ADH_2.0_HA-Ox_2.0 (40 kDa)_ vs. HA-ADH_1.0_HA-Ox_3.0 (100 kDa)_ | ns | 0.0852 |
| HA-ADH_2.0_HA-Ox_2.0 (40 kDa)_ vs. HA-ADH_1.0_HA-Ox_1.67 (100 kDa)_ | ns | 0.2629 |
| HA-ADH_2.0_HA-Ox_2.0 (40 kDa)_ vs. HA-ADH_1.0_HA-Ox_1.0 (100 kDa)_ | ns | 0.3485 |
| HA-ADH_1.67_HA-Ox_1.0 (40 kDa)_ vs. HA-ADH_1.0_HA-Ox_3.0 (40 kDa)_ | * | 0.0156 |
| HA-ADH_1.67_HA-Ox_1.0 (40 kDa)_ vs. HA-ADH_1.0_HA-Ox_1.0 (40 kDa)_ | * | 0.0141 |
| HA-ADH_1.67_HA-Ox_1.0 (40 kDa)_ vs. HA-ADH_3.0_HA-Ox_3.0 (100 kDa)_ | * | 0.0325 |
| HA-ADH_1.67_HA-Ox_1.0 (40 kDa)_ vs. HA-ADH_3.0_HA-Ox_2.33 (100 kDa)_ | ns | 0.0959 |
| HA-ADH_1.67_HA-Ox_1.0 (40 kDa)_ vs. HA-ADH_3.0_HA-Ox_1.0 (100 kDa)_ | ns | 0.1362 |
| HA-ADH_1.67_HA-Ox_1.0 (40 kDa)_ vs. HA-ADH_2.33_HA-Ox_1.0 (100 kDa)_ | ns | 0.0983 |
| HA-ADH_1.67_HA-Ox_1.0 (40 kDa)_ vs. HA-ADH_2.0_HA-Ox_2.0 (100 kDa)_ | * | 0.03 |
| HA-ADH_1.67_HA-Ox_1.0 (40 kDa)_ vs. HA-ADH_1.67_HA-Ox_3.0 (100 kDa)_ | * | 0.0385 |
| HA-ADH_1.67_HA-Ox_1.0 (40 kDa)_ vs. HA-ADH_1.0_HA-Ox_3.0 (100 kDa)_ | * | 0.0326 |
| HA-ADH_1.67_HA-Ox_1.0 (40 kDa)_ vs. HA-ADH_1.0_HA-Ox_1.67 (100 kDa)_ | ns | 0.083 |
| HA-ADH_1.67_HA-Ox_1.0 (40 kDa)_ vs. HA-ADH_1.0_HA-Ox_1.0 (100 kDa)_ | ns | 0.0859 |
| HA-ADH_1.0_HA-Ox_3.0 (40 kDa)_ vs. HA-ADH_1.0_HA-Ox_1.0 (40 kDa)_ | ns | >0.9999 |
| HA-ADH_1.0_HA-Ox_3.0 (40 kDa)_ vs. HA-ADH_3.0_HA-Ox_3.0 (100 kDa)_ | * | 0.0231 |
| HA-ADH_1.0_HA-Ox_3.0 (40 kDa)_ vs. HA-ADH_3.0_HA-Ox_2.33 (100 kDa)_ | ** | 0.0054 |
| HA-ADH_1.0_HA-Ox_3.0 (40 kDa)_ vs. HA-ADH_3.0_HA-Ox_1.0 (100 kDa)_ | ** | 0.0057 |
| HA-ADH_1.0_HA-Ox_3.0 (40 kDa)_ vs. HA-ADH_2.33_HA-Ox_1.0 (100 kDa)_ | * | 0.0362 |
| HA-ADH_1.0_HA-Ox_3.0 (40 kDa)_ vs. HA-ADH_2.0_HA-Ox_2.0 (100 kDa)_ | * | 0.0282 |
| HA-ADH_1.0_HA-Ox_3.0 (40 kDa)_ vs. HA-ADH_1.67_HA-Ox_3.0 (100 kDa)_ | * | 0.0481 |
| HA-ADH_1.0_HA-Ox_3.0 (40 kDa)_ vs. HA-ADH_1.0_HA-Ox_3.0 (100 kDa)_ | ns | 0.362 |
| HA-ADH_1.0_HA-Ox_3.0 (40 kDa)_ vs. HA-ADH_1.0_HA-Ox_1.67 (100 kDa)_ | ns | 0.1778 |
| HA-ADH_1.0_HA-Ox_3.0 (40 kDa)_ vs. HA-ADH_1.0_HA-Ox_1.0 (100 kDa)_ | ns | 0.1041 |
| HA-ADH_1.0_HA-Ox_1.0 (40 kDa)_ vs. HA-ADH_3.0_HA-Ox_3.0 (100 kDa)_ | * | 0.0188 |
| HA-ADH_1.0_HA-Ox_1.0 (40 kDa)_ vs. HA-ADH_3.0_HA-Ox_2.33 (100 kDa)_ | ** | 0.0027 |
| HA-ADH_1.0_HA-Ox_1.0 (40 kDa)_ vs. HA-ADH_3.0_HA-Ox_1.0 (100 kDa)_ | ** | 0.0031 |
| HA-ADH_1.0_HA-Ox_1.0 (40 kDa)_ vs. HA-ADH_2.33_HA-Ox_1.0 (100 kDa)_ | * | 0.0333 |
| HA-ADH_1.0_HA-Ox_1.0 (40 kDa)_ vs. HA-ADH_2.0_HA-Ox_2.0 (100 kDa)_ | * | 0.0237 |
| HA-ADH_1.0_HA-Ox_1.0 (40 kDa)_ vs. HA-ADH_1.67_HA-Ox_3.0 (100 kDa)_ | * | 0.0444 |
| HA-ADH_1.0_HA-Ox_1.0 (40 kDa)_ vs. HA-ADH_1.0_HA-Ox_3.0 (100 kDa)_ | ns | 0.3645 |
| HA-ADH_1.0_HA-Ox_1.0 (40 kDa)_ vs. HA-ADH_1.0_HA-Ox_1.67 (100 kDa)_ | ns | 0.1769 |
| HA-ADH_1.0_HA-Ox_1.0 (40 kDa)_ vs. HA-ADH_1.0_HA-Ox_1.0 (100 kDa)_ | ns | 0.1021 |
| HA-ADH_3.0_HA-Ox_3.0 (100 kDa)_ vs. HA-ADH_3.0_HA-Ox_2.33 (100 kDa)_ | ns | 0.3062 |
| HA-ADH_3.0_HA-Ox_3.0 (100 kDa)_ vs. HA-ADH_3.0_HA-Ox_1.0 (100 kDa)_ | ns | 0.1386 |
| HA-ADH_3.0_HA-Ox_3.0 (100 kDa)_ vs. HA-ADH_2.33_HA-Ox_1.0 (100 kDa)_ | ns | 0.6462 |
| HA-ADH_3.0_HA-Ox_3.0 (100 kDa)_ vs. HA-ADH_2.0_HA-Ox_2.0 (100 kDa)_ | ns | >0.9999 |
| HA-ADH_3.0_HA-Ox_3.0 (100 kDa)_ vs. HA-ADH_1.67_HA-Ox_3.0 (100 kDa)_ | ns | >0.9999 |
| HA-ADH_3.0_HA-Ox_3.0 (100 kDa)_ vs. HA-ADH_1.0_HA-Ox_3.0 (100 kDa)_ | ns | 0.6112 |
| HA-ADH_3.0_HA-Ox_3.0 (100 kDa)_ vs. HA-ADH_1.0_HA-Ox_1.67 (100 kDa)_ | ns | >0.9999 |
| HA-ADH_3.0_HA-Ox_3.0 (100 kDa)_ vs. HA-ADH_1.0_HA-Ox_1.0 (100 kDa)_ | ns | >0.9999 |
| HA-ADH_3.0_HA-Ox_2.33 (100 kDa)_ vs. HA-ADH_3.0_HA-Ox_1.0 (100 kDa)_ | ns | 0.9148 |
| HA-ADH_3.0_HA-Ox_2.33 (100 kDa)_ vs. HA-ADH_2.33_HA-Ox_1.0 (100 kDa)_ | ns | >0.9999 |
| HA-ADH_3.0_HA-Ox_2.33 (100 kDa)_ vs. HA-ADH_2.0_HA-Ox_2.0 (100 kDa)_ | ns | 0.3058 |
| HA-ADH_3.0_HA-Ox_2.33 (100 kDa)_ vs. HA-ADH_1.67_HA-Ox_3.0 (100 kDa)_ | ns | 0.6844 |
| HA-ADH_3.0_HA-Ox_2.33 (100 kDa)_ vs. HA-ADH_1.0_HA-Ox_3.0 (100 kDa)_ | ns | 0.2558 |
| HA-ADH_3.0_HA-Ox_2.33 (100 kDa)_ vs. HA-ADH_1.0_HA-Ox_1.67 (100 kDa)_ | ns | 0.7057 |
| HA-ADH_3.0_HA-Ox_2.33 (100 kDa)_ vs. HA-ADH_1.0_HA-Ox_1.0 (100 kDa)_ | ns | 0.9137 |
| HA-ADH_3.0_HA-Ox_1.0 (100 kDa)_ vs. HA-ADH_2.33_HA-Ox_1.0 (100 kDa)_ | ns | 0.9914 |
| HA-ADH_3.0_HA-Ox_1.0 (100 kDa)_ vs. HA-ADH_2.0_HA-Ox_2.0 (100 kDa)_ | ns | 0.145 |
| HA-ADH_3.0_HA-Ox_1.0 (100 kDa)_ vs. HA-ADH_1.67_HA-Ox_3.0 (100 kDa)_ | ns | 0.3953 |
| HA-ADH_3.0_HA-Ox_1.0 (100 kDa)_ vs. HA-ADH_1.0_HA-Ox_3.0 (100 kDa)_ | ns | 0.1926 |
| HA-ADH_3.0_HA-Ox_1.0 (100 kDa)_ vs. HA-ADH_1.0_HA-Ox_1.67 (100 kDa)_ | ns | 0.5219 |
| HA-ADH_3.0_HA-Ox_1.0 (100 kDa)_ vs. HA-ADH_1.0_HA-Ox_1.0 (100 kDa)_ | ns | 0.6955 |
| HA-ADH_2.33_HA-Ox_1.0 (100 kDa)_ vs. HA-ADH_2.0_HA-Ox_2.0 (100 kDa)_ | ns | 0.6174 |
| HA-ADH_2.33_HA-Ox_1.0 (100 kDa)_ vs. HA-ADH_1.67_HA-Ox_3.0 (100 kDa)_ | ns | 0.8849 |
| HA-ADH_2.33_HA-Ox_1.0 (100 kDa)_ vs. HA-ADH_1.0_HA-Ox_3.0 (100 kDa)_ | ns | 0.2652 |
| HA-ADH_2.33_HA-Ox_1.0 (100 kDa)_ vs. HA-ADH_1.0_HA-Ox_1.67 (100 kDa)_ | ns | 0.8038 |
| HA-ADH_2.33_HA-Ox_1.0 (100 kDa)_ vs. HA-ADH_1.0_HA-Ox_1.0 (100 kDa)_ | ns | 0.974 |
| HA-ADH_2.0_HA-Ox_2.0 (100 kDa)_ vs. HA-ADH_1.67_HA-Ox_3.0 (100 kDa)_ | ns | >0.9999 |
| HA-ADH_2.0_HA-Ox_2.0 (100 kDa)_ vs. HA-ADH_1.0_HA-Ox_3.0 (100 kDa)_ | ns | 0.6462 |
| HA-ADH_2.0_HA-Ox_2.0 (100 kDa)_ vs. HA-ADH_1.0_HA-Ox_1.67 (100 kDa)_ | ns | >0.9999 |
| HA-ADH_2.0_HA-Ox_2.0 (100 kDa)_ vs. HA-ADH_1.0_HA-Ox_1.0 (100 kDa)_ | ns | >0.9999 |
| HA-ADH_1.67_HA-Ox_3.0 (100 kDa)_ vs. HA-ADH_1.0_HA-Ox_3.0 (100 kDa)_ | ns | 0.5666 |
| HA-ADH_1.67_HA-Ox_3.0 (100 kDa)_ vs. HA-ADH_1.0_HA-Ox_1.67 (100 kDa)_ | ns | 0.9997 |
| HA-ADH_1.67_HA-Ox_3.0 (100 kDa)_ vs. HA-ADH_1.0_HA-Ox_1.0 (100 kDa)_ | ns | >0.9999 |
| HA-ADH_1.0_HA-Ox_3.0 (100 kDa)_ vs. HA-ADH_1.0_HA-Ox_1.67 (100 kDa)_ | ns | 0.928 |
| HA-ADH_1.0_HA-Ox_3.0 (100 kDa)_ vs. HA-ADH_1.0_HA-Ox_1.0 (100 kDa)_ | ns | 0.6496 |
| HA-ADH_1.0_HA-Ox_1.67 (100 kDa)_ vs. HA-ADH_1.0_HA-Ox_1.0 (100 kDa)_ | ns | 0.9998 |
| **Day 21** | | |
| HA-ADH_3.0_HA-Ox_3.0 (40 kDa)_ vs. HA-ADH_3.0_HA-Ox_1.0 (40 kDa)_ | ns | 0.2215 |
| HA-ADH_3.0_HA-Ox_3.0 (40 kDa)_ vs. HA-ADH_2.0_HA-Ox_2.0 (40 kDa)_ | ns | >0.9999 |
| HA-ADH_3.0_HA-Ox_3.0 (40 kDa)_ vs. HA-ADH_1.67_HA-Ox_1.0 (40 kDa)_ | ns | 0.8054 |
| HA-ADH_3.0_HA-Ox_3.0 (40 kDa)_ vs. HA-ADH_1.0_HA-Ox_3.0 (40 kDa)_ | ns | 0.0537 |
| HA-ADH_3.0_HA-Ox_3.0 (40 kDa)_ vs. HA-ADH_1.0_HA-Ox_1.0 (40 kDa)_ | ns | 0.052 |
| HA-ADH_3.0_HA-Ox_3.0 (40 kDa)_ vs. HA-ADH_3.0_HA-Ox_3.0 (100 kDa)_ | ns | 0.2209 |
| HA-ADH_3.0_HA-Ox_3.0 (40 kDa)_ vs. HA-ADH_3.0_HA-Ox_2.33 (100 kDa)_ | ns | 0.4186 |
| HA-ADH_3.0_HA-Ox_3.0 (40 kDa)_ vs. HA-ADH_3.0_HA-Ox_1.0 (100 kDa)_ | ns | 0.8093 |
| HA-ADH_3.0_HA-Ox_3.0 (40 kDa)_ vs. HA-ADH_2.33_HA-Ox_1.0 (100 kDa)_ | ns | 0.5724 |
| HA-ADH_3.0_HA-Ox_3.0 (40 kDa)_ vs. HA-ADH_2.0_HA-Ox_2.0 (100 kDa)_ | ns | 0.3462 |
| HA-ADH_3.0_HA-Ox_3.0 (40 kDa)_ vs. HA-ADH_1.67_HA-Ox_3.0 (100 kDa)_ | ns | 0.6414 |
| HA-ADH_3.0_HA-Ox_3.0 (40 kDa)_ vs. HA-ADH_1.0_HA-Ox_3.0 (100 kDa)_ | * | 0.05 |
| HA-ADH_3.0_HA-Ox_3.0 (40 kDa)_ vs. HA-ADH_1.0_HA-Ox_1.67 (100 kDa)_ | ns | 0.8314 |
| HA-ADH_3.0_HA-Ox_3.0 (40 kDa)_ vs. HA-ADH_1.0_HA-Ox_1.0 (100 kDa)_ | ns | 0.4346 |
| HA-ADH_3.0_HA-Ox_1.0 (40 kDa)_ vs. HA-ADH_2.0_HA-Ox_2.0 (40 kDa)_ | ns | 0.2288 |
| HA-ADH_3.0_HA-Ox_1.0 (40 kDa)_ vs. HA-ADH_1.67_HA-Ox_1.0 (40 kDa)_ | ns | 0.4165 |
| HA-ADH_3.0_HA-Ox_1.0 (40 kDa)_ vs. HA-ADH_1.0_HA-Ox_3.0 (40 kDa)_ | ns | 0.0523 |
| HA-ADH_3.0_HA-Ox_1.0 (40 kDa)_ vs. HA-ADH_1.0_HA-Ox_1.0 (40 kDa)_ | ns | 0.0517 |
| HA-ADH_3.0_HA-Ox_1.0 (40 kDa)_ vs. HA-ADH_3.0_HA-Ox_3.0 (100 kDa)_ | ns | 0.0994 |
| HA-ADH_3.0_HA-Ox_1.0 (40 kDa)_ vs. HA-ADH_3.0_HA-Ox_2.33 (100 kDa)_ | ns | 0.0962 |
| HA-ADH_3.0_HA-Ox_1.0 (40 kDa)_ vs. HA-ADH_3.0_HA-Ox_1.0 (100 kDa)_ | ns | 0.133 |
| HA-ADH_3.0_HA-Ox_1.0 (40 kDa)_ vs. HA-ADH_2.33_HA-Ox_1.0 (100 kDa)_ | ns | 0.132 |
| HA-ADH_3.0_HA-Ox_1.0 (40 kDa)_ vs. HA-ADH_2.0_HA-Ox_2.0 (100 kDa)_ | ns | 0.0968 |
| HA-ADH_3.0_HA-Ox_1.0 (40 kDa)_ vs. HA-ADH_1.67_HA-Ox_3.0 (100 kDa)_ | ns | 0.1098 |
| HA-ADH_3.0_HA-Ox_1.0 (40 kDa)_ vs. HA-ADH_1.0_HA-Ox_3.0 (100 kDa)_ | * | 0.0495 |
| HA-ADH_3.0_HA-Ox_1.0 (40 kDa)_ vs. HA-ADH_1.0_HA-Ox_1.67 (100 kDa)_ | ns | 0.1034 |
| HA-ADH_3.0_HA-Ox_1.0 (40 kDa)_ vs. HA-ADH_1.0_HA-Ox_1.0 (100 kDa)_ | ns | 0.0965 |
| HA-ADH_2.0_HA-Ox_2.0 (40 kDa)_ vs. HA-ADH_1.67_HA-Ox_1.0 (40 kDa)_ | ns | 0.5479 |
| HA-ADH_2.0_HA-Ox_2.0 (40 kDa)_ vs. HA-ADH_1.0_HA-Ox_3.0 (40 kDa)_ | * | 0.0148 |
| HA-ADH_2.0_HA-Ox_2.0 (40 kDa)_ vs. HA-ADH_1.0_HA-Ox_1.0 (40 kDa)_ | * | 0.013 |
| HA-ADH_2.0_HA-Ox_2.0 (40 kDa)_ vs. HA-ADH_3.0_HA-Ox_3.0 (100 kDa)_ | * | 0.0415 |
| HA-ADH_2.0_HA-Ox_2.0 (40 kDa)_ vs. HA-ADH_3.0_HA-Ox_2.33 (100 kDa)_ | ns | 0.2038 |
| HA-ADH_2.0_HA-Ox_2.0 (40 kDa)_ vs. HA-ADH_3.0_HA-Ox_1.0 (100 kDa)_ | ns | 0.5384 |
| HA-ADH_2.0_HA-Ox_2.0 (40 kDa)_ vs. HA-ADH_2.33_HA-Ox_1.0 (100 kDa)_ | ns | 0.2089 |
| HA-ADH_2.0_HA-Ox_2.0 (40 kDa)_ vs. HA-ADH_2.0_HA-Ox_2.0 (100 kDa)_ | ns | 0.1285 |
| HA-ADH_2.0_HA-Ox_2.0 (40 kDa)_ vs. HA-ADH_1.67_HA-Ox_3.0 (100 kDa)_ | ns | 0.397 |
| HA-ADH_2.0_HA-Ox_2.0 (40 kDa)_ vs. HA-ADH_1.0_HA-Ox_3.0 (100 kDa)_ | ** | 0.0085 |
| HA-ADH_2.0_HA-Ox_2.0 (40 kDa)_ vs. HA-ADH_1.0_HA-Ox_1.67 (100 kDa)_ | ns | 0.7396 |
| HA-ADH_2.0_HA-Ox_2.0 (40 kDa)_ vs. HA-ADH_1.0_HA-Ox_1.0 (100 kDa)_ | ns | 0.2194 |
| HA-ADH_1.67_HA-Ox_1.0 (40 kDa)_ vs. HA-ADH_1.0_HA-Ox_3.0 (40 kDa)_ | * | 0.0173 |
| HA-ADH_1.67_HA-Ox_1.0 (40 kDa)_ vs. HA-ADH_1.0_HA-Ox_1.0 (40 kDa)_ | * | 0.0161 |
| HA-ADH_1.67_HA-Ox_1.0 (40 kDa)_ vs. HA-ADH_3.0_HA-Ox_3.0 (100 kDa)_ | * | 0.0362 |
| HA-ADH_1.67_HA-Ox_1.0 (40 kDa)_ vs. HA-ADH_3.0_HA-Ox_2.33 (100 kDa)_ | ns | 0.0602 |
| HA-ADH_1.67_HA-Ox_1.0 (40 kDa)_ vs. HA-ADH_3.0_HA-Ox_1.0 (100 kDa)_ | ns | 0.1296 |
| HA-ADH_1.67_HA-Ox_1.0 (40 kDa)_ vs. HA-ADH_2.33_HA-Ox_1.0 (100 kDa)_ | ns | 0.0746 |
| HA-ADH_1.67_HA-Ox_1.0 (40 kDa)_ vs. HA-ADH_2.0_HA-Ox_2.0 (100 kDa)_ | * | 0.0433 |
| HA-ADH_1.67_HA-Ox_1.0 (40 kDa)_ vs. HA-ADH_1.67_HA-Ox_3.0 (100 kDa)_ | ns | 0.1067 |
| HA-ADH_1.67_HA-Ox_1.0 (40 kDa)_ vs. HA-ADH_1.0_HA-Ox_3.0 (100 kDa)_ | ** | 0.0062 |
| HA-ADH_1.67_HA-Ox_1.0 (40 kDa)_ vs. HA-ADH_1.0_HA-Ox_1.67 (100 kDa)_ | ns | 0.3496 |
| HA-ADH_1.67_HA-Ox_1.0 (40 kDa)_ vs. HA-ADH_1.0_HA-Ox_1.0 (100 kDa)_ | ns | 0.064 |
| HA-ADH_1.0_HA-Ox_3.0 (40 kDa)_ vs. HA-ADH_1.0_HA-Ox_1.0 (40 kDa)_ | ns | >0.9999 |
| HA-ADH_1.0_HA-Ox_3.0 (40 kDa)_ vs. HA-ADH_3.0_HA-Ox_3.0 (100 kDa)_ | * | 0.0145 |
| HA-ADH_1.0_HA-Ox_3.0 (40 kDa)_ vs. HA-ADH_3.0_HA-Ox_2.33 (100 kDa)_ | ns | 0.065 |
| HA-ADH_1.0_HA-Ox_3.0 (40 kDa)_ vs. HA-ADH_3.0_HA-Ox_1.0 (100 kDa)_ | * | 0.0391 |
| HA-ADH_1.0_HA-Ox_3.0 (40 kDa)_ vs. HA-ADH_2.33_HA-Ox_1.0 (100 kDa)_ | * | 0.0171 |
| HA-ADH_1.0_HA-Ox_3.0 (40 kDa)_ vs. HA-ADH_2.0_HA-Ox_2.0 (100 kDa)_ | ns | 0.0531 |
| HA-ADH_1.0_HA-Ox_3.0 (40 kDa)_ vs. HA-ADH_1.67_HA-Ox_3.0 (100 kDa)_ | ns | 0.0611 |
| HA-ADH_1.0_HA-Ox_3.0 (40 kDa)_ vs. HA-ADH_1.0_HA-Ox_3.0 (100 kDa)_ | ns | 0.2976 |
| HA-ADH_1.0_HA-Ox_3.0 (40 kDa)_ vs. HA-ADH_1.0_HA-Ox_1.67 (100 kDa)_ | ns | 0.2014 |
| HA-ADH_1.0_HA-Ox_3.0 (40 kDa)_ vs. HA-ADH_1.0_HA-Ox_1.0 (100 kDa)_ | ns | 0.0664 |
| HA-ADH_1.0_HA-Ox_1.0 (40 kDa)_ vs. HA-ADH_3.0_HA-Ox_3.0 (100 kDa)_ | * | 0.0104 |
| HA-ADH_1.0_HA-Ox_1.0 (40 kDa)_ vs. HA-ADH_3.0_HA-Ox_2.33 (100 kDa)_ | ns | 0.0618 |
| HA-ADH_1.0_HA-Ox_1.0 (40 kDa)_ vs. HA-ADH_3.0_HA-Ox_1.0 (100 kDa)_ | * | 0.0367 |
| HA-ADH_1.0_HA-Ox_1.0 (40 kDa)_ vs. HA-ADH_2.33_HA-Ox_1.0 (100 kDa)_ | * | 0.0143 |
| HA-ADH_1.0_HA-Ox_1.0 (40 kDa)_ vs. HA-ADH_2.0_HA-Ox_2.0 (100 kDa)_ | * | 0.0497 |
| HA-ADH_1.0_HA-Ox_1.0 (40 kDa)_ vs. HA-ADH_1.67_HA-Ox_3.0 (100 kDa)_ | ns | 0.0584 |
| HA-ADH_1.0_HA-Ox_1.0 (40 kDa)_ vs. HA-ADH_1.0_HA-Ox_3.0 (100 kDa)_ | ns | 0.2844 |
| HA-ADH_1.0_HA-Ox_1.0 (40 kDa)_ vs. HA-ADH_1.0_HA-Ox_1.67 (100 kDa)_ | ns | 0.1981 |
| HA-ADH_1.0_HA-Ox_1.0 (40 kDa)_ vs. HA-ADH_1.0_HA-Ox_1.0 (100 kDa)_ | ns | 0.0633 |
| HA-ADH_3.0_HA-Ox_3.0 (100 kDa)_ vs. HA-ADH_3.0_HA-Ox_2.33 (100 kDa)_ | ns | 0.9289 |
| HA-ADH_3.0_HA-Ox_3.0 (100 kDa)_ vs. HA-ADH_3.0_HA-Ox_1.0 (100 kDa)_ | ns | 0.3451 |
| HA-ADH_3.0_HA-Ox_3.0 (100 kDa)_ vs. HA-ADH_2.33_HA-Ox_1.0 (100 kDa)_ | ns | 0.2547 |
| HA-ADH_3.0_HA-Ox_3.0 (100 kDa)_ vs. HA-ADH_2.0_HA-Ox_2.0 (100 kDa)_ | ns | 0.9641 |
| HA-ADH_3.0_HA-Ox_3.0 (100 kDa)_ vs. HA-ADH_1.67_HA-Ox_3.0 (100 kDa)_ | ns | 0.6719 |
| HA-ADH_3.0_HA-Ox_3.0 (100 kDa)_ vs. HA-ADH_1.0_HA-Ox_3.0 (100 kDa)_ | ns | 0.1406 |
| HA-ADH_3.0_HA-Ox_3.0 (100 kDa)_ vs. HA-ADH_1.0_HA-Ox_1.67 (100 kDa)_ | ns | 0.9736 |
| HA-ADH_3.0_HA-Ox_3.0 (100 kDa)_ vs. HA-ADH_1.0_HA-Ox_1.0 (100 kDa)_ | ns | 0.9185 |
| HA-ADH_3.0_HA-Ox_2.33 (100 kDa)_ vs. HA-ADH_3.0_HA-Ox_1.0 (100 kDa)_ | ns | 0.9381 |
| HA-ADH_3.0_HA-Ox_2.33 (100 kDa)_ vs. HA-ADH_2.33_HA-Ox_1.0 (100 kDa)_ | ns | 0.9901 |
| HA-ADH_3.0_HA-Ox_2.33 (100 kDa)_ vs. HA-ADH_2.0_HA-Ox_2.0 (100 kDa)_ | ns | >0.9999 |
| HA-ADH_3.0_HA-Ox_2.33 (100 kDa)_ vs. HA-ADH_1.67_HA-Ox_3.0 (100 kDa)_ | ns | 0.9997 |
| HA-ADH_3.0_HA-Ox_2.33 (100 kDa)_ vs. HA-ADH_1.0_HA-Ox_3.0 (100 kDa)_ | ns | 0.1191 |
| HA-ADH_3.0_HA-Ox_2.33 (100 kDa)_ vs. HA-ADH_1.0_HA-Ox_1.67 (100 kDa)_ | ns | >0.9999 |
| HA-ADH_3.0_HA-Ox_2.33 (100 kDa)_ vs. HA-ADH_1.0_HA-Ox_1.0 (100 kDa)_ | ns | >0.9999 |
| HA-ADH_3.0_HA-Ox_1.0 (100 kDa)_ vs. HA-ADH_2.33_HA-Ox_1.0 (100 kDa)_ | ns | 0.9998 |
| HA-ADH_3.0_HA-Ox_1.0 (100 kDa)_ vs. HA-ADH_2.0_HA-Ox_2.0 (100 kDa)_ | ns | 0.8315 |
| HA-ADH_3.0_HA-Ox_1.0 (100 kDa)_ vs. HA-ADH_1.67_HA-Ox_3.0 (100 kDa)_ | ns | 0.9999 |
| HA-ADH_3.0_HA-Ox_1.0 (100 kDa)_ vs. HA-ADH_1.0_HA-Ox_3.0 (100 kDa)_ | * | 0.0408 |
| HA-ADH_3.0_HA-Ox_1.0 (100 kDa)_ vs. HA-ADH_1.0_HA-Ox_1.67 (100 kDa)_ | ns | >0.9999 |
| HA-ADH_3.0_HA-Ox_1.0 (100 kDa)_ vs. HA-ADH_1.0_HA-Ox_1.0 (100 kDa)_ | ns | 0.9517 |
| HA-ADH_2.33_HA-Ox_1.0 (100 kDa)_ vs. HA-ADH_2.0_HA-Ox_2.0 (100 kDa)_ | ns | 0.9314 |
| HA-ADH_2.33_HA-Ox_1.0 (100 kDa)_ vs. HA-ADH_1.67_HA-Ox_3.0 (100 kDa)_ | ns | >0.9999 |
| HA-ADH_2.33_HA-Ox_1.0 (100 kDa)_ vs. HA-ADH_1.0_HA-Ox_3.0 (100 kDa)_ | * | 0.0341 |
| HA-ADH_2.33_HA-Ox_1.0 (100 kDa)_ vs. HA-ADH_1.0_HA-Ox_1.67 (100 kDa)_ | ns | >0.9999 |
| HA-ADH_2.33_HA-Ox_1.0 (100 kDa)_ vs. HA-ADH_1.0_HA-Ox_1.0 (100 kDa)_ | ns | 0.9941 |
| HA-ADH_2.0_HA-Ox_2.0 (100 kDa)_ vs. HA-ADH_1.67_HA-Ox_3.0 (100 kDa)_ | ns | 0.9946 |
| HA-ADH_2.0_HA-Ox_2.0 (100 kDa)_ vs. HA-ADH_1.0_HA-Ox_3.0 (100 kDa)_ | ns | 0.1065 |
| HA-ADH_2.0_HA-Ox_2.0 (100 kDa)_ vs. HA-ADH_1.0_HA-Ox_1.67 (100 kDa)_ | ns | >0.9999 |
| HA-ADH_2.0_HA-Ox_2.0 (100 kDa)_ vs. HA-ADH_1.0_HA-Ox_1.0 (100 kDa)_ | ns | >0.9999 |
| HA-ADH_1.67_HA-Ox_3.0 (100 kDa)_ vs. HA-ADH_1.0_HA-Ox_3.0 (100 kDa)_ | ns | 0.0881 |
| HA-ADH_1.67_HA-Ox_3.0 (100 kDa)_ vs. HA-ADH_1.0_HA-Ox_1.67 (100 kDa)_ | ns | >0.9999 |
| HA-ADH_1.67_HA-Ox_3.0 (100 kDa)_ vs. HA-ADH_1.0_HA-Ox_1.0 (100 kDa)_ | ns | 0.9999 |
| HA-ADH_1.0_HA-Ox_3.0 (100 kDa)_ vs. HA-ADH_1.0_HA-Ox_1.67 (100 kDa)_ | ns | 0.3893 |
| HA-ADH_1.0_HA-Ox_3.0 (100 kDa)_ vs. HA-ADH_1.0_HA-Ox_1.0 (100 kDa)_ | ns | 0.1199 |
| HA-ADH_1.0_HA-Ox_1.67 (100 kDa)_ vs. HA-ADH_1.0_HA-Ox_1.0 (100 kDa)_ | ns | >0.9999 |
| **Day 28** | | |
| HA-ADH_3.0_HA-Ox_3.0 (40 kDa)_ vs. HA-ADH_3.0_HA-Ox_1.0 (40 kDa)_ | ns | 0.0887 |
| HA-ADH_3.0_HA-Ox_3.0 (40 kDa)_ vs. HA-ADH_2.0_HA-Ox_2.0 (40 kDa)_ | ns | 0.8973 |
| HA-ADH_3.0_HA-Ox_3.0 (40 kDa)_ vs. HA-ADH_1.67_HA-Ox_1.0 (40 kDa)_ | ns | 0.4655 |
| HA-ADH_3.0_HA-Ox_3.0 (40 kDa)_ vs. HA-ADH_1.0_HA-Ox_3.0 (40 kDa)_ | ** | 0.0025 |
| HA-ADH_3.0_HA-Ox_3.0 (40 kDa)_ vs. HA-ADH_1.0_HA-Ox_1.0 (40 kDa)_ | ** | 0.0015 |
| HA-ADH_3.0_HA-Ox_3.0 (40 kDa)_ vs. HA-ADH_3.0_HA-Ox_3.0 (100 kDa)_ | * | 0.0108 |
| HA-ADH_3.0_HA-Ox_3.0 (40 kDa)_ vs. HA-ADH_3.0_HA-Ox_2.33 (100 kDa)_ | * | 0.013 |
| HA-ADH_3.0_HA-Ox_3.0 (40 kDa)_ vs. HA-ADH_3.0_HA-Ox_1.0 (100 kDa)_ | ns | 0.9982 |
| HA-ADH_3.0_HA-Ox_3.0 (40 kDa)_ vs. HA-ADH_2.33_HA-Ox_1.0 (100 kDa)_ | ns | 0.2129 |
| HA-ADH_3.0_HA-Ox_3.0 (40 kDa)_ vs. HA-ADH_2.0_HA-Ox_2.0 (100 kDa)_ | ns | 0.2207 |
| HA-ADH_3.0_HA-Ox_3.0 (40 kDa)_ vs. HA-ADH_1.67_HA-Ox_3.0 (100 kDa)_ | ** | 0.0032 |
| HA-ADH_3.0_HA-Ox_3.0 (40 kDa)_ vs. HA-ADH_1.0_HA-Ox_3.0 (100 kDa)_ | * | 0.0127 |
| HA-ADH_3.0_HA-Ox_3.0 (40 kDa)_ vs. HA-ADH_1.0_HA-Ox_1.67 (100 kDa)_ | ns | 0.1384 |
| HA-ADH_3.0_HA-Ox_3.0 (40 kDa)_ vs. HA-ADH_1.0_HA-Ox_1.0 (100 kDa)_ | ns | 0.2095 |
| HA-ADH_3.0_HA-Ox_1.0 (40 kDa)_ vs. HA-ADH_2.0_HA-Ox_2.0 (40 kDa)_ | ns | 0.2687 |
| HA-ADH_3.0_HA-Ox_1.0 (40 kDa)_ vs. HA-ADH_1.67_HA-Ox_1.0 (40 kDa)_ | ns | 0.2282 |
| HA-ADH_3.0_HA-Ox_1.0 (40 kDa)_ vs. HA-ADH_1.0_HA-Ox_3.0 (40 kDa)_ | * | 0.018 |
| HA-ADH_3.0_HA-Ox_1.0 (40 kDa)_ vs. HA-ADH_1.0_HA-Ox_1.0 (40 kDa)_ | * | 0.0173 |
| HA-ADH_3.0_HA-Ox_1.0 (40 kDa)_ vs. HA-ADH_3.0_HA-Ox_3.0 (100 kDa)_ | * | 0.0224 |
| HA-ADH_3.0_HA-Ox_1.0 (40 kDa)_ vs. HA-ADH_3.0_HA-Ox_2.33 (100 kDa)_ | * | 0.0442 |
| HA-ADH_3.0_HA-Ox_1.0 (40 kDa)_ vs. HA-ADH_3.0_HA-Ox_1.0 (100 kDa)_ | ns | 0.0557 |
| HA-ADH_3.0_HA-Ox_1.0 (40 kDa)_ vs. HA-ADH_2.33_HA-Ox_1.0 (100 kDa)_ | * | 0.0148 |
| HA-ADH_3.0_HA-Ox_1.0 (40 kDa)_ vs. HA-ADH_2.0_HA-Ox_2.0 (100 kDa)_ | * | 0.0164 |
| HA-ADH_3.0_HA-Ox_1.0 (40 kDa)_ vs. HA-ADH_1.67_HA-Ox_3.0 (100 kDa)_ | * | 0.025 |
| HA-ADH_3.0_HA-Ox_1.0 (40 kDa)_ vs. HA-ADH_1.0_HA-Ox_3.0 (100 kDa)_ | ** | 0.0064 |
| HA-ADH_3.0_HA-Ox_1.0 (40 kDa)_ vs. HA-ADH_1.0_HA-Ox_1.67 (100 kDa)_ | ** | 0.0096 |
| HA-ADH_3.0_HA-Ox_1.0 (40 kDa)_ vs. HA-ADH_1.0_HA-Ox_1.0 (100 kDa)_ | * | 0.0133 |
| HA-ADH_2.0_HA-Ox_2.0 (40 kDa)_ vs. HA-ADH_1.67_HA-Ox_1.0 (40 kDa)_ | ns | >0.9999 |
| HA-ADH_2.0_HA-Ox_2.0 (40 kDa)_ vs. HA-ADH_1.0_HA-Ox_3.0 (40 kDa)_ | ns | 0.061 |
| HA-ADH_2.0_HA-Ox_2.0 (40 kDa)_ vs. HA-ADH_1.0_HA-Ox_1.0 (40 kDa)_ | ns | 0.0601 |
| HA-ADH_2.0_HA-Ox_2.0 (40 kDa)_ vs. HA-ADH_3.0_HA-Ox_3.0 (100 kDa)_ | ns | 0.1631 |
| HA-ADH_2.0_HA-Ox_2.0 (40 kDa)_ vs. HA-ADH_3.0_HA-Ox_2.33 (100 kDa)_ | ns | 0.2372 |
| HA-ADH_2.0_HA-Ox_2.0 (40 kDa)_ vs. HA-ADH_3.0_HA-Ox_1.0 (100 kDa)_ | ns | 0.8603 |
| HA-ADH_2.0_HA-Ox_2.0 (40 kDa)_ vs. HA-ADH_2.33_HA-Ox_1.0 (100 kDa)_ | ns | 0.224 |
| HA-ADH_2.0_HA-Ox_2.0 (40 kDa)_ vs. HA-ADH_2.0_HA-Ox_2.0 (100 kDa)_ | ns | 0.2533 |
| HA-ADH_2.0_HA-Ox_2.0 (40 kDa)_ vs. HA-ADH_1.67_HA-Ox_3.0 (100 kDa)_ | ns | 0.1493 |
| HA-ADH_2.0_HA-Ox_2.0 (40 kDa)_ vs. HA-ADH_1.0_HA-Ox_3.0 (100 kDa)_ | ns | 0.052 |
| HA-ADH_2.0_HA-Ox_2.0 (40 kDa)_ vs. HA-ADH_1.0_HA-Ox_1.67 (100 kDa)_ | ns | 0.0979 |
| HA-ADH_2.0_HA-Ox_2.0 (40 kDa)_ vs. HA-ADH_1.0_HA-Ox_1.0 (100 kDa)_ | ns | 0.1733 |
| HA-ADH_1.67_HA-Ox_1.0 (40 kDa)_ vs. HA-ADH_1.0_HA-Ox_3.0 (40 kDa)_ | * | 0.0253 |
| HA-ADH_1.67_HA-Ox_1.0 (40 kDa)_ vs. HA-ADH_1.0_HA-Ox_1.0 (40 kDa)_ | * | 0.0242 |
| HA-ADH_1.67_HA-Ox_1.0 (40 kDa)_ vs. HA-ADH_3.0_HA-Ox_3.0 (100 kDa)_ | * | 0.0452 |
| HA-ADH_1.67_HA-Ox_1.0 (40 kDa)_ vs. HA-ADH_3.0_HA-Ox_2.33 (100 kDa)_ | ns | 0.0936 |
| HA-ADH_1.67_HA-Ox_1.0 (40 kDa)_ vs. HA-ADH_3.0_HA-Ox_1.0 (100 kDa)_ | ns | 0.5128 |
| HA-ADH_1.67_HA-Ox_1.0 (40 kDa)_ vs. HA-ADH_2.33_HA-Ox_1.0 (100 kDa)_ | ns | 0.0679 |
| HA-ADH_1.67_HA-Ox_1.0 (40 kDa)_ vs. HA-ADH_2.0_HA-Ox_2.0 (100 kDa)_ | ns | 0.0741 |
| HA-ADH_1.67_HA-Ox_1.0 (40 kDa)_ vs. HA-ADH_1.67_HA-Ox_3.0 (100 kDa)_ | * | 0.0459 |
| HA-ADH_1.67_HA-Ox_1.0 (40 kDa)_ vs. HA-ADH_1.0_HA-Ox_3.0 (100 kDa)_ | ** | 0.0095 |
| HA-ADH_1.67_HA-Ox_1.0 (40 kDa)_ vs. HA-ADH_1.0_HA-Ox_1.67 (100 kDa)_ | * | 0.0396 |
| HA-ADH_1.67_HA-Ox_1.0 (40 kDa)_ vs. HA-ADH_1.0_HA-Ox_1.0 (100 kDa)_ | ns | 0.0633 |
| HA-ADH_1.0_HA-Ox_3.0 (40 kDa)_ vs. HA-ADH_1.0_HA-Ox_1.0 (40 kDa)_ | ns | >0.9999 |
| HA-ADH_1.0_HA-Ox_3.0 (40 kDa)_ vs. HA-ADH_3.0_HA-Ox_3.0 (100 kDa)_ | * | 0.0301 |
| HA-ADH_1.0_HA-Ox_3.0 (40 kDa)_ vs. HA-ADH_3.0_HA-Ox_2.33 (100 kDa)_ | **** | <0.0001 |
| HA-ADH_1.0_HA-Ox_3.0 (40 kDa)_ vs. HA-ADH_3.0_HA-Ox_1.0 (100 kDa)_ | ns | 0.0639 |
| HA-ADH_1.0_HA-Ox_3.0 (40 kDa)_ vs. HA-ADH_2.33_HA-Ox_1.0 (100 kDa)_ | ns | 0.0914 |
| HA-ADH_1.0_HA-Ox_3.0 (40 kDa)_ vs. HA-ADH_2.0_HA-Ox_2.0 (100 kDa)_ | ns | 0.0732 |
| HA-ADH_1.0_HA-Ox_3.0 (40 kDa)_ vs. HA-ADH_1.67_HA-Ox_3.0 (100 kDa)_ | * | 0.0216 |
| HA-ADH_1.0_HA-Ox_3.0 (40 kDa)_ vs. HA-ADH_1.0_HA-Ox_3.0 (100 kDa)_ | ns | 0.4367 |
| HA-ADH_1.0_HA-Ox_3.0 (40 kDa)_ vs. HA-ADH_1.0_HA-Ox_1.67 (100 kDa)_ | ns | 0.3021 |
| HA-ADH_1.0_HA-Ox_3.0 (40 kDa)_ vs. HA-ADH_1.0_HA-Ox_1.0 (100 kDa)_ | ns | 0.1566 |
| HA-ADH_1.0_HA-Ox_1.0 (40 kDa)_ vs. HA-ADH_3.0_HA-Ox_3.0 (100 kDa)_ | * | 0.0248 |
| HA-ADH_1.0_HA-Ox_1.0 (40 kDa)_ vs. HA-ADH_3.0_HA-Ox_2.33 (100 kDa)_ | **** | <0.0001 |
| HA-ADH_1.0_HA-Ox_1.0 (40 kDa)_ vs. HA-ADH_3.0_HA-Ox_1.0 (100 kDa)_ | ns | 0.0625 |
| HA-ADH_1.0_HA-Ox_1.0 (40 kDa)_ vs. HA-ADH_2.33_HA-Ox_1.0 (100 kDa)_ | ns | 0.0891 |
| HA-ADH_1.0_HA-Ox_1.0 (40 kDa)_ vs. HA-ADH_2.0_HA-Ox_2.0 (100 kDa)_ | ns | 0.0707 |
| HA-ADH_1.0_HA-Ox_1.0 (40 kDa)_ vs. HA-ADH_1.67_HA-Ox_3.0 (100 kDa)_ | * | 0.0156 |
| HA-ADH_1.0_HA-Ox_1.0 (40 kDa)_ vs. HA-ADH_1.0_HA-Ox_3.0 (100 kDa)_ | ns | 0.4361 |
| HA-ADH_1.0_HA-Ox_1.0 (40 kDa)_ vs. HA-ADH_1.0_HA-Ox_1.67 (100 kDa)_ | ns | 0.3009 |
| HA-ADH_1.0_HA-Ox_1.0 (40 kDa)_ vs. HA-ADH_1.0_HA-Ox_1.0 (100 kDa)_ | ns | 0.1547 |
| HA-ADH_3.0_HA-Ox_3.0 (100 kDa)_ vs. HA-ADH_3.0_HA-Ox_2.33 (100 kDa)_ | ns | 0.6017 |
| HA-ADH_3.0_HA-Ox_3.0 (100 kDa)_ vs. HA-ADH_3.0_HA-Ox_1.0 (100 kDa)_ | ns | 0.2775 |
| HA-ADH_3.0_HA-Ox_3.0 (100 kDa)_ vs. HA-ADH_2.33_HA-Ox_1.0 (100 kDa)_ | ns | 0.9758 |
| HA-ADH_3.0_HA-Ox_3.0 (100 kDa)_ vs. HA-ADH_2.0_HA-Ox_2.0 (100 kDa)_ | ns | 0.8782 |
| HA-ADH_3.0_HA-Ox_3.0 (100 kDa)_ vs. HA-ADH_1.67_HA-Ox_3.0 (100 kDa)_ | ns | 0.9973 |
| HA-ADH_3.0_HA-Ox_3.0 (100 kDa)_ vs. HA-ADH_1.0_HA-Ox_3.0 (100 kDa)_ | ns | 0.1094 |
| HA-ADH_3.0_HA-Ox_3.0 (100 kDa)_ vs. HA-ADH_1.0_HA-Ox_1.67 (100 kDa)_ | ns | 0.9713 |
| HA-ADH_3.0_HA-Ox_3.0 (100 kDa)_ vs. HA-ADH_1.0_HA-Ox_1.0 (100 kDa)_ | ns | >0.9999 |
| HA-ADH_3.0_HA-Ox_2.33 (100 kDa)_ vs. HA-ADH_3.0_HA-Ox_1.0 (100 kDa)_ | ns | 0.4403 |
| HA-ADH_3.0_HA-Ox_2.33 (100 kDa)_ vs. HA-ADH_2.33_HA-Ox_1.0 (100 kDa)_ | ns | >0.9999 |
| HA-ADH_3.0_HA-Ox_2.33 (100 kDa)_ vs. HA-ADH_2.0_HA-Ox_2.0 (100 kDa)_ | ns | >0.9999 |
| HA-ADH_3.0_HA-Ox_2.33 (100 kDa)_ vs. HA-ADH_1.67_HA-Ox_3.0 (100 kDa)_ | ns | 0.23 |
| HA-ADH_3.0_HA-Ox_2.33 (100 kDa)_ vs. HA-ADH_1.0_HA-Ox_3.0 (100 kDa)_ | ns | 0.0885 |
| HA-ADH_3.0_HA-Ox_2.33 (100 kDa)_ vs. HA-ADH_1.0_HA-Ox_1.67 (100 kDa)_ | ns | 0.6834 |
| HA-ADH_3.0_HA-Ox_2.33 (100 kDa)_ vs. HA-ADH_1.0_HA-Ox_1.0 (100 kDa)_ | ns | 0.998 |
| HA-ADH_3.0_HA-Ox_1.0 (100 kDa)_ vs. HA-ADH_2.33_HA-Ox_1.0 (100 kDa)_ | ns | 0.5763 |
| HA-ADH_3.0_HA-Ox_1.0 (100 kDa)_ vs. HA-ADH_2.0_HA-Ox_2.0 (100 kDa)_ | ns | 0.6545 |
| HA-ADH_3.0_HA-Ox_1.0 (100 kDa)_ vs. HA-ADH_1.67_HA-Ox_3.0 (100 kDa)_ | ns | 0.2314 |
| HA-ADH_3.0_HA-Ox_1.0 (100 kDa)_ vs. HA-ADH_1.0_HA-Ox_3.0 (100 kDa)_ | ns | 0.0562 |
| HA-ADH_3.0_HA-Ox_1.0 (100 kDa)_ vs. HA-ADH_1.0_HA-Ox_1.67 (100 kDa)_ | ns | 0.2151 |
| HA-ADH_3.0_HA-Ox_1.0 (100 kDa)_ vs. HA-ADH_1.0_HA-Ox_1.0 (100 kDa)_ | ns | 0.4379 |
| HA-ADH_2.33_HA-Ox_1.0 (100 kDa)_ vs. HA-ADH_2.0_HA-Ox_2.0 (100 kDa)_ | ns | >0.9999 |
| HA-ADH_2.33_HA-Ox_1.0 (100 kDa)_ vs. HA-ADH_1.67_HA-Ox_3.0 (100 kDa)_ | ns | 0.8422 |
| HA-ADH_2.33_HA-Ox_1.0 (100 kDa)_ vs. HA-ADH_1.0_HA-Ox_3.0 (100 kDa)_ | ns | 0.1372 |
| HA-ADH_2.33_HA-Ox_1.0 (100 kDa)_ vs. HA-ADH_1.0_HA-Ox_1.67 (100 kDa)_ | ns | 0.8311 |
| HA-ADH_2.33_HA-Ox_1.0 (100 kDa)_ vs. HA-ADH_1.0_HA-Ox_1.0 (100 kDa)_ | ns | >0.9999 |
| HA-ADH_2.0_HA-Ox_2.0 (100 kDa)_ vs. HA-ADH_1.67_HA-Ox_3.0 (100 kDa)_ | ns | 0.6715 |
| HA-ADH_2.0_HA-Ox_2.0 (100 kDa)_ vs. HA-ADH_1.0_HA-Ox_3.0 (100 kDa)_ | ns | 0.0946 |
| HA-ADH_2.0_HA-Ox_2.0 (100 kDa)_ vs. HA-ADH_1.0_HA-Ox_1.67 (100 kDa)_ | ns | 0.7148 |
| HA-ADH_2.0_HA-Ox_2.0 (100 kDa)_ vs. HA-ADH_1.0_HA-Ox_1.0 (100 kDa)_ | ns | 0.9969 |
| HA-ADH_1.67_HA-Ox_3.0 (100 kDa)_ vs. HA-ADH_1.0_HA-Ox_3.0 (100 kDa)_ | ns | 0.1566 |
| HA-ADH_1.67_HA-Ox_3.0 (100 kDa)_ vs. HA-ADH_1.0_HA-Ox_1.67 (100 kDa)_ | ns | 0.998 |
| HA-ADH_1.67_HA-Ox_3.0 (100 kDa)_ vs. HA-ADH_1.0_HA-Ox_1.0 (100 kDa)_ | ns | 0.9992 |
| HA-ADH_1.0_HA-Ox_3.0 (100 kDa)_ vs. HA-ADH_1.0_HA-Ox_1.67 (100 kDa)_ | ns | 0.7591 |
| HA-ADH_1.0_HA-Ox_3.0 (100 kDa)_ vs. HA-ADH_1.0_HA-Ox_1.0 (100 kDa)_ | ns | 0.3106 |
| HA-ADH_1.0_HA-Ox_1.67 (100 kDa)_ vs. HA-ADH_1.0_HA-Ox_1.0 (100 kDa)_ | ns | 0.9872 |

**Table 4. Summary of significance for 40 kDa HA mass change and 100 kDa HA mass change.** A summary of significance for the mass change for both 40 kDa and 100 kDa is listed, a two-way ANOVA post-hoc Tukey’s multiple comparisons was performed. n = 3; * p < 0.05, ** < p 0.01, *** p < 0.001, **** p < 0.0001.

*Affibody Bioconjugation*

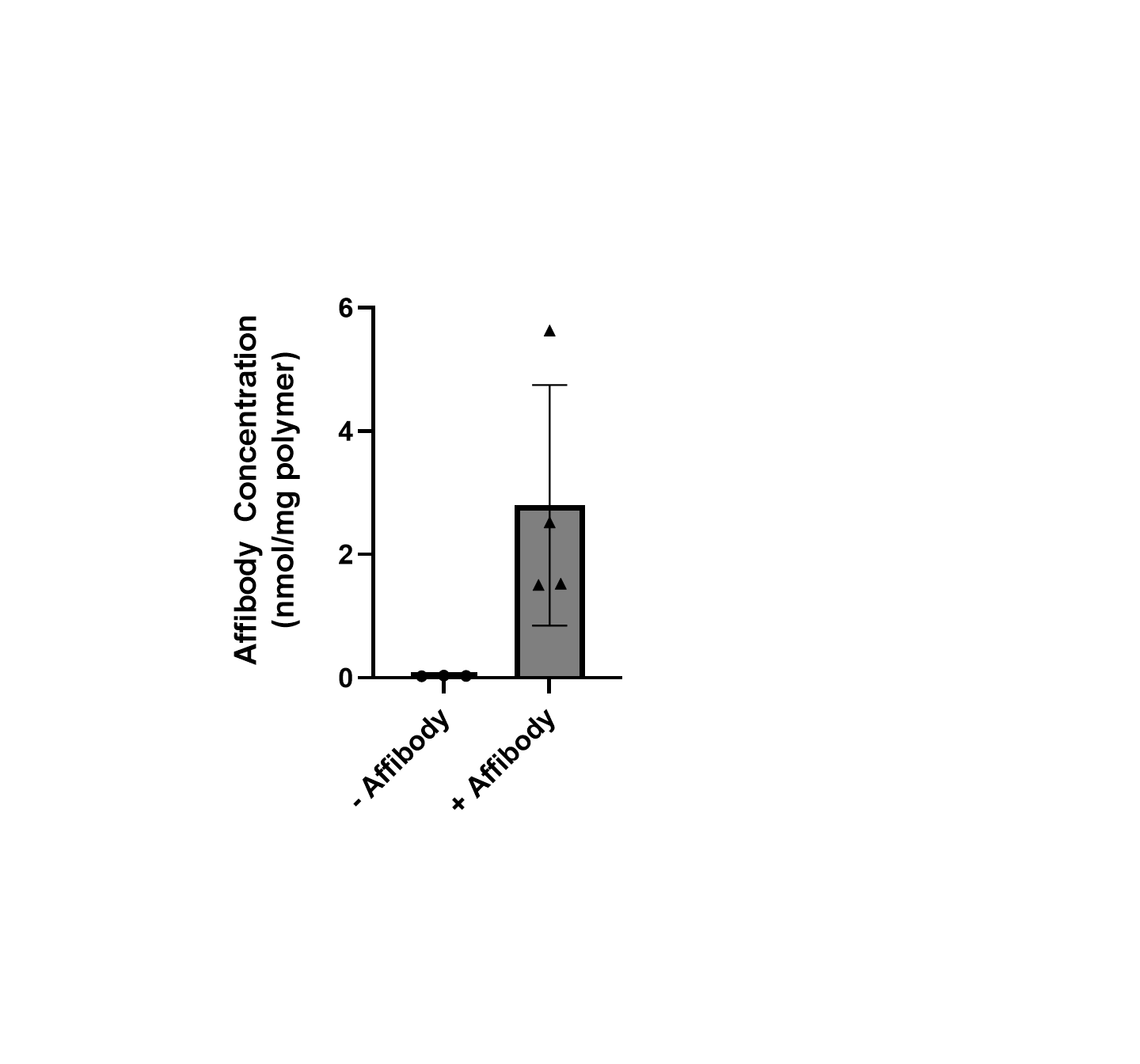

**Figure S3. Quantification of affibody conjugation on HA-Nor-Ox-Affibody polymer.** Pierce 660 Protein Assay was used to measure the amount of affibody conjugated per mg of HA-Nor-Ox-Affibody polymer and compared to HA-Nor-Ox polymer without conjugated affibodies as a control.
